# Integrative optical genome mapping and long-read sequencing resolve constitutional complex rearrangements at nucleotide resolution

**DOI:** 10.64898/2026.08.27.747510

**Authors:** Bruna Burssed, Bart van der Sanden, Wolfram Höps, Kornelia Neveling, Eveline Kamping, Ronald van Beek, Amber den Ouden, Ronny Derks, Raoul Timmermans, Eduardo Perrone, Marco Antonio Ramos, Fernanda Teixeira Bellucco, Alexander Hoischen, Maria Isabel Melaragno

## Abstract

Complex rearrangements are one of the rarest types of structural variants (SVs) and can be divided into two categories: complex chromosomal rearrangements (CCRs) and complex genomic rearrangements (CGRs). CCRs include structural rearrangements that present at least three breakpoints and show exchange of genetic material between more than two chromosomes and CGRs are rearrangements that present more than one junction and/or more than one SV in *cis*. They are usually formed by one of the chromoanagenesis mechanisms, where a massive disruptive cellular event leads to multiple structural rearrangements. Classical cytogenomic techniques have been commonly applied for their characterization, but methodologies that involve longer DNA molecules, namely optical genome mapping (OGM) and long-read genome sequencing (lrGS), present a considerably higher SV detection resolution, revealing more details about the rearrangements, including precise breakpoint location. Here, we describe six patients with complex rearrangements investigated through a combination of different techniques: karyotyping, chromosomal microarray, and OGM were performed to characterize the rearrangements. Subsequently, lrGS was used to further resolve the alterations, refine their breakpoints’ location, and sequence their junction points. Three patients presented CCRs involving three, four, and six chromosomes, while three exhibited CGRs involving one different chromosome each, providing a variety of complex SVs to show the importance of each technique and their combination in rearrangement resolution. In total, the complex rearrangements presented 127 breakpoints, 66 junction points and involved 14 of the 24 chromosomes. Higher-resolution techniques revealed additional complexity in all cases. Despite the advances provided by OGM and lrGS, conventional karyotyping remained indispensable for complete rearrangement resolution. In two patients, the findings supported a novel mechanism combining features of the different chromoanagenesis processes. Furthermore, evidence of inherited alterations was identified, and the comprehensive characterization of the rearrangements enabled more accurate genotype–phenotype correlations. Our findings indicate that an integrated approach combining karyotyping, OGM, and lrGS can completely resolve SVs, including complex rearrangements.

## INTRODUCTION

Structural variants (SVs) are abnormalities in the chromosome structure known to cause chromosomal and genomic disorders (Lupski, 1998; Shaw and Lupski, 2004; Lupski and Stankiewicz, 2005; Weckselblatt and Rudd, 2015). They may affect the phenotype due to abnormal dosage of genes, caused by copy number variants (CNVs), gene rupture, and gene fusion, among other mechanisms (Lupski, 1998; Shaffer and Lupski, 2000; Gu, Zhang and Lupski, 2008). Among the types of SV, complex rearrangements are among the rarest (Czakó *et al*., 2025; Kocagil *et al*., 2026) and they can be divided into two categories (Schuy *et al*., 2022). Complex chromosomal rearrangements (CCRs) include structural rearrangements that present at least three breakpoints and show exchange of genetic material between more than two chromosomes. Complex genomic rearrangements (CGRs) are rearrangements that present more than one junction and/or more than one SV in *cis*. The most evident difference between the two is the number of involved chromosomes: CCRs involve at least three chromosomes, as in translocations with multiple breakpoints affecting various chromosomes (Priya *et al*., 2018), while CGRs involve a single chromosome, such as inverted duplications associated with terminal deletions (inv-dup-del) (Burssed *et al*., 2023).

SVs result from different mutational mechanisms that include DNA recombination, repair, and replication processes (Carvalho and Lupski, 2016). The study of these mechanisms is important as it can help us understand how cells respond to DNA damage and the nature of the DNA sequences involved in these rearrangements. The study of the rearrangement’s breakpoints through DNA sequencing allows us to recognize signatures of these processes and to identify risk factors for such rearrangements, thus guiding us in inferring their mechanisms of formation (Weckselblatt and Rudd, 2015).

Several mechanisms of formation of constitutional chromosomal alterations have been described, such as Non-Allelic Homologous Recombination (NAHR), Non-Homologous End-Joining (NHEJ), Microhomology-Mediated End-Joining (MMEJ), Fork Stalling and Template Switching (FoSTeS), and Microhomology-Mediated Break-Induced Replication (MMBIR) (Burssed *et al*., 2022). CCRs are likely to be formed in a single catastrophic event that leads to multiple structural rearrangements in one or more chromosomes (Pellestor and Gatinois, 2018; Zepeda-Mendoza and Morton, 2019; Burssed *et al*., 2022). Three mutational events leading to CCRs, collectively known as chromoanagenesis, have been reported. Chromothripsis involves a single chromosome that shatters and then is reassembled through NHEJ; chromoanasynthesis also involves a single chromosome that undergoes DNA segment re-synthesis via FoSTeS or MMBIR; and chromoplexy involves more than two chromosomes that are broken and rejoined through NHEJ (Pellestor and Gatinois, 2018; Zepeda-Mendoza and Morton, 2019; Burssed et al., 2022).

Structural rearrangements can be studied through many techniques. G-banding karyotyping allows for the detection of balanced and unbalanced alterations; however, it has a low diagnostic rate of around 10%, a low resolution (5–10 Mb), and requires a trained cytogeneticist for the analysis (Stankiewicz, Pursley and Cheung, 2010; Mantere *et al*., 2021). Chromosomal microarray is a technique with much higher resolution (∼50-150 kb, though it varies per platform), which can assess copy number throughout the genome, although not being able to identify balanced alterations or locate extra copies of genomic regions (Miller *et al*., 2010; Ciuladaite *et al*., 2014; Mantere *et al*., 2021). Fluorescence in situ hybridization (FISH) can reveal the alterations’ location and orientation by using fluorescent-labeled DNA probes specific to certain regions of the chromosome, therefore it is harder to be performed on its own since a region of interest must be known *a priori* (Weckselblatt and Rudd, 2015; Neveling *et al*., 2021). Optical genome mapping (OGM) uses linearized ultra-high molecular weight DNA molecules labeled at specific sites, which allows for complicated and repetitive regions to be spanned more easily than with short molecules, thus detecting balanced and unbalanced SVs across the genome (Dremsek *et al*., 2021; Mantere *et al*., 2021). OGM’s analysis has two pipelines: the CNV pipeline relies on coverage depth to detect copy number while the SV pipeline looks at fusions seen in the patient’s maps to encounter alterations in chromosome structure and also copy number changes when they occur with abnormal fusion (Bionano, 2024). OGM appears as an alternative that can overcome the limitations presented by the classical cytogenetics techniques: it has a much higher resolution than karyotype (500 base pairs against 5 Mb) and is able to detect balanced alterations, locate additional material, and find breakpoints with more precision than microarray (Dremsek *et al*., 2021). Also, it does not require any cell cultivation before processing, therefore allowing the analysis to occur much faster than karyotyping or FISH (Dremsek *et al*., 2021). Whole genome sequencing (WGS) allows for the detection of single nucleotide variants (SNVs) as well as SVs (Weckselblatt and Rudd, 2015). Short-read sequencing is becoming increasingly affordable and emerges as a first-tier test in modern laboratories. However, it is limited in read length (usually 2x150 base pairs) and therefore has challenges in resolving SVs, especially in the presence of repetitive regions (Mantere *et al*., 2021). Long-read genome sequencing (lrGS), on the other hand, uses longer reads originating from single DNA molecules which are more likely to span breakpoints and can more confidently be aligned to repetitive sequences that are often involved in the formation of SVs, including segmental duplications (SDs) and repetitive elements (REs) such as Alu and LINE elements (Weckselblatt and Rudd, 2015; Mantere, Kersten and Hoischen, 2019; Eisfeldt *et al*., 2023).

Often, one requires multiple of the above-mentioned techniques can assist in the detection and characterization of the rearrangement and location of its breakpoints (Burssed *et al*., 2022). A comprehensive analysis of breakpoints and junctions is necessary in order to define with more certainty the mechanism of formation of the rearrangement. Sequencing of the breakpoint at the nucleotide level, either with WGS or with Sanger sequencing, allows the analysis of the presence of information scars from NHEJ, microhomology from MMEJ, FoSTeS, and MMBIR, and inserted segments from the replication-based mechanisms (Burssed *et al*., 2022; Eisfeldt *et al*., 2023). The full characterization of the rearrangement’s junction points is also crucial since it assists in positioning the rearranged genomic segments, phasing multiple breakpoints, and pinpointing interrupted gene(s) or regulatory interaction(s), which are important for the correlation with the phenotype (Schuy et al., 2022).

Here, we describe complex constitutional rearrangements in six patients, three harboring CCRs and three presenting CGRs. Karyotyping, chromosomal microarray, and OGM were used to characterize the rearrangements while lrGS was performed to further resolve the alterations, refine their breakpoints’ location, and sequence their junction points. This approach allowed us to fully resolve all breakpoints of the complex rearrangements, enabling us to infer their mechanisms of formation and identify genomic alterations affecting coding or regulatory regions that may underlie the patients’ phenotypes.

## MATERIALS AND METHODS

### Enrollment following karyotype analysis

We studied six patients with complex rearrangements. The patients were selected based on their G-banding karyotype at 550-band resolution performed from lymphocyte cultures following their evaluation by the geneticists of the Medical Genetics Center of the Universidade Federal de São Paulo, Brazil. Peripheral blood of patients and their parents (when available) was collected after written informed consent and approval of the local ethics committee (CAAE 40846114.2.0000.5505, CEP 0028/2015; CAAE 78269424.4.0000.5505, CEP 0215/2024).

### Chromosomal Microarray Analysis (CMA)

DNA extraction was performed using Gentra Puregene Blood Kit (Qiagen Sciences, MD, USA) according to the manufacturer’s protocol. The DNA quality and quantification were assessed with NanoDrop ND 1000 (Thermo Technologies, Haute Savoie, France). Chromosomal microarrays were performed according to the manufacturer’s protocol using different platforms: KaryoNIM^®^ 180K (NIMGenetics, Madrid, Spain and Agilent Technologies, CA, USA) for patient 1, CytoScan^®^ 750 K Array for patients 2–5 (Affymetrix, CA, USA), and CGH+SNP-Array 180K (Agilent) for patient 6. The analysis was carried out using the Cytogenomics (Agilent) and ChAS (Affymetrix) software with GRCh37/hg19 annotation. Genomic coordinates were subsequently converted to GRCh38/hg38 using the LiftOver tool available through the UCSC Genome Browser (Casper *et al*., 2026).

### Optical Genome Mapping (OGM)

OGM was performed as previously described (Mantere *et al*., 2021; Neveling *et al*., 2021). In short, ultra-high molecular weight (UHMW) DNA was isolated from 650 µL of whole peripheral blood (EDTA) using the SP Blood and Cell Culture DNA Isolation Kit according to manufacturers’ instructions (Bionano Genomics, San Diego, CA, USA). UHMW genomic DNA (gDNA) molecules were labeled with the DLS (Direct Label and Stain) DNA Labeling Kit (Bionano Genomic). We used Direct Label Enzyme (DLE-1) and DL-green fluorophores to label 750 ng of gDNA. After a wash-out of the DL-green fluorophores excess, the DNA backbone was counterstained overnight before quantitation. Labeled UHMW gDNA was loaded on a Saphyr chip for linearization and imaging on the Saphyr instrument (Bionano Genomics). The *de novo* assembly and variant annotation pipeline were executed with Bionano Solve software v.3.7 based on the GRCh38/hg38 reference genome. Results were analyzed through two distinct pipelines: a CNV pipeline that allows for the detection of large unbalanced aberrations based on normalized molecule coverage and an SV pipeline that compares the labeling patterns between the constructed sample genome maps and a reference genome map. Reporting and direct visualization of structural variants were performed using Bionano Access software v.1.7.

Since OGM is not a sequencing technique, it is not able to precisely locate the breakpoints. A chimeric map may present an uncertain region highlighted in purple that has not been assigned to either of the regions involved in the junction due to a lack of alignment or labels. The precise breakpoints should be located within this region.

### Long-read Genome Sequencing (lrGS)

Library preparation was performed according to the manufacturer’s instructions. Briefly, 7 µg of DNA was sheared on Megaruptor 3 (Diagenode, Liège, Belgium) to a target size of ± 15–18 kb. Libraries were then prepared using SMRTbell prep kit 3.0 (PacBio, Menlo Park, CA, USA), followed by size selection for fragments >10 kb on the BluePippin system (Sage Science, Beverly, MA, USA). Samples were equimolarly pooled and sequencing polymerase was bound to SMRTbell library using the Revio polymerase kit (PacBio, Menlo Park, CA, USA). Finished libraries were loaded onto a 25M Revio SMRT cell and sequenced on a Revio instrument (PacBio, Menlo Park, CA, USA), according to the manufacturer’s instructions. Alignment (pbmm2 v1.10.0) of High Fidelity (HiFi) reads was performed against the GRCh38/hg38 reference genome for all cases and also the T2T-CHM13 reference genome patients 3, 5, and 6. Sample of the patients’ parents were also submitted to lrGS when available to investigate inheritance. SVs were called using pbsv (v2.9.0) and small variants using DeepVariant (v1.5.0). Copy number variants were detected using variation in depth of coverages using HifiCNV (v0.1.6). Finally, short tandem repeats were called using TRGT (v0.4.0). All variants were annotated using in-house pipelines and publicly available databases.

### Breakpoints and junction points assessment

Rearrangement breakpoints correspond to the unaffected nucleotide closest to the break of a region and junction point refer to the joining of different breakpoints that link regions together. Therefore, breakpoints have a coordinate and can be located while junction points are formed by at least two breakpoints and can be characterized (e.g. as presenting insertions or micro-homology). Breakpoint location and junction point characterization was based on annotations and on visual inspection of chimeric reads using the software Integrative Genomics Viewer (IGV) 2.4.14 (Broad Institute and the Regents of the University of California). Read sequences were analyzed with the BLAT tool on the UCSC Genome Browser. Repetitive elements (REs) were annotated within a 2 kb range surrounding each breakpoint using the RepeatMasker track of the UCSC Genome Browser. Breakpoints were numbered according to the order that their coordinates appear in the normal chromosomes. Junction points were numbered according to the order that they appear in the derivative chromosomes.

### Karyotype-phenotype correlation

Annotation and ranking of structural variants and affected genes were assessed using Franklin by Genoox (https://franklin.genoox.com) along with the AnnotSV tool (Geoffroy *et al*., 2018) following proper rearrangement characterization. The AnnotSV tool compiles regulatory and clinically relevant information with data from the Database of Genomic Variants (DGV), Deciphering Developmental Disorders (DDD) Study, Database of Genomic Structural Variation (dbVar), Genome Aggregation Database (gnomAD), Clinical Genome Resource (ClinGen), and Online Mendelian Inheritance in Man (OMIM) databases. The DatabasE of genomiC varIation and Phenotype in Humans using Ensembl Resources (DECIPHER, https://decipher.sanger.ac.uk/) as well as previous studies from the literature were used to perform a karyotype-phenotype correlation.

## RESULTS

All six rearrangements were fully resolved to the nucleotide level, with three patients presenting CCRs and three harboring CGRs. In total, 127 breakpoints were detected and 66 junction points were characterized. Fourteen of the 24 chromosomes were affected, with chromosomes 3 and 14 being involved in two different CCRs. Insertions and micro-homology at the junction points allowed for the inference of chromoanasynthesis as the mechanism of formation of two patients while its combination with chromoplexy was proposed for another two. With a precise breakpoint location, ruptured genes and disrupted regulatory elements were identified and correlated with the patients’ phenotypes. Results of each patient are presented below and on their respective supplementary files.

### Patient 1

Patient 1 was a 6-year-old girl and the second daughter of a non-consanguineous couple. Both parents had normal karyotypes, and their first daughter was healthy. The patient presented with neuropsychomotor developmental delay, intellectual disability, microcephaly, plagiocephaly, hypoplasia of the corpus callosum, hypotonia, strabismus, widely spaced nipples, pes cavus, lower limb asymmetry, protruding ears, and facial dysmorphisms, including prominent forehead, bitemporal narrowness, synophrys, depressed nasal bridge, smooth philtrum, thin lips with downturned corners of mouth, and widely spaced teeth (Supplementary File 1 – Supplementary Table 1). She was admitted to the hospital due to respiratory issues and had corrective surgery for femur fracture, bilateral tympanoplasty, as well as adenoid removal.

Patient 1’s initial karyotype (Figure 1A) analysis revealed a possible alteration in chromosome 12, with it being either a deletion or a derivative chromosome, while chromosomal microarray analysis unveiled a *de novo* 2.4 Mb deletion in chromosome 4 (Supplementary File 1 – Supplementary Figure 1, Supplementary Table 2). The difference in these results between karyotyping and CMA prompted us to perform OGM to resolve the patient’s alterations.

**Figure 1.**
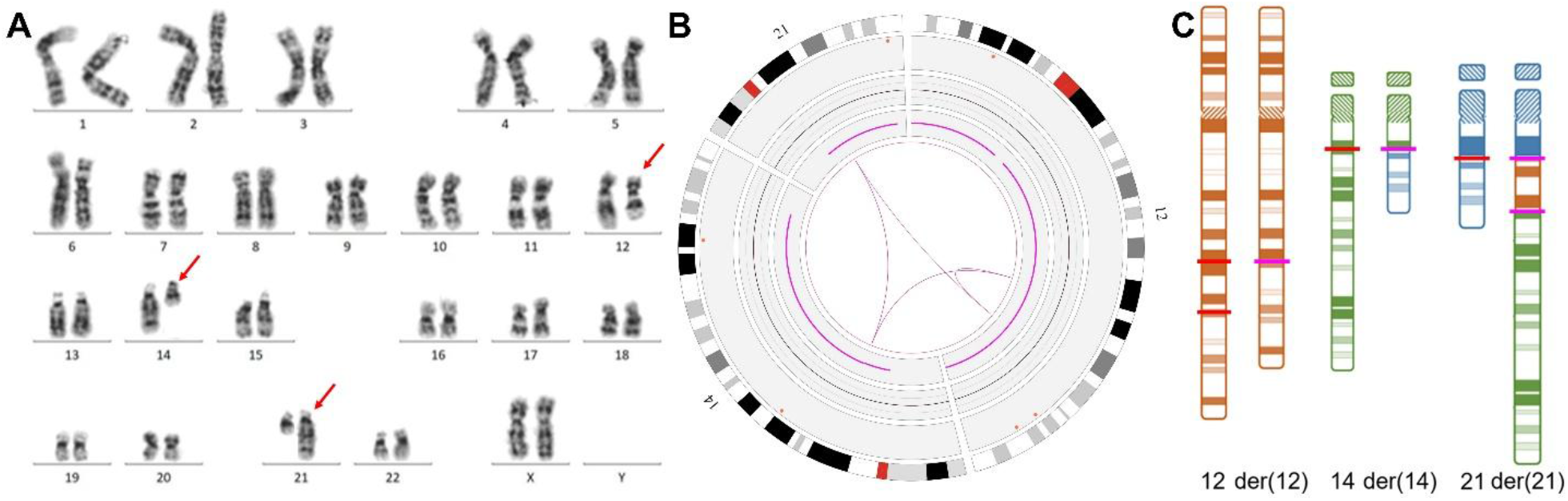
Patient 1’s results. **(A) G-banding karyotype with red arrows showing the der(12), der(14), and der(21)**. Initially, only der(12) was identified while der(14) and der(21)’s places were switched. **(B) Circos plot showing chromosomes 12, 14, and 21**. The chromosomes’ idiograms can be seen in the periphery while the pink lines in the middle represent the translocations. **(C) Idiogram of the chromosomes involved in the complex chromosomal rearrangement**. Red lines show the breakpoints in the normal chromosomes and pink lines show the junction points in the derivative chromosomes. Patient 1 presents eight breakpoints and four junctions.

A similar ∼2.4 Mb deletion in chromosome 4 was detected by OGM, while it also revealed a complex chromosomal rearrangement between chromosomes 12, 14, and 21 (Figure 1B, Supplementary File 1 – Supplementary Table 2, Supplementary Figure 2). The rearrangement was resolved with a thorough evaluation of the patient’s molecule maps, which led to an updated karyotype of 46,XX,t(12;14;21)(12pter→12q21.32::12q23.3→12qter;14pter→14q12:: 21q21.1→21qter;21pter→21q21.1::12q23.3→12q21.32::14q12→14qter)dn (Figure 1C, Supplementary File 1 – Supplementary Figure 3).

Based on the OGM results, we analyzed the long-read sequencing in order to better characterize the breakpoints. All eight breakpoints and four junction points of the complex rearrangement were found and fully characterized (Figure 2, Supplementary File 1 – Supplementary Table 3, Supplementary Table 4). The three junction points from der(14) and der(21) show a simple ligation between the breakpoints. The junction point from der(12), however, presents an insertion between both chromosome 12 breakpoints which corresponds to a repetition of the final 12 nucleotides of the upstream breakpoint (Figure 2B). Upon closer analysis of the breakpoints, we could see a deletion of 1 base pair ([GRCh38] chr12:86,717,791) and a duplication of 3 base pairs ([GRCh38] chr21:22,080,569-22,080,571). The deletion in chromosome 4 was called by the HiFi CNV variant caller but no chimeric reads were found to precisely characterize the rearrangement with nucleotide-level resolution. The patient’s mother was investigated through lrGS and presenting normal results. The patient’s father is deceased therefore we cannot affirm that the CCR is *de novo*.

**Figure 2.**
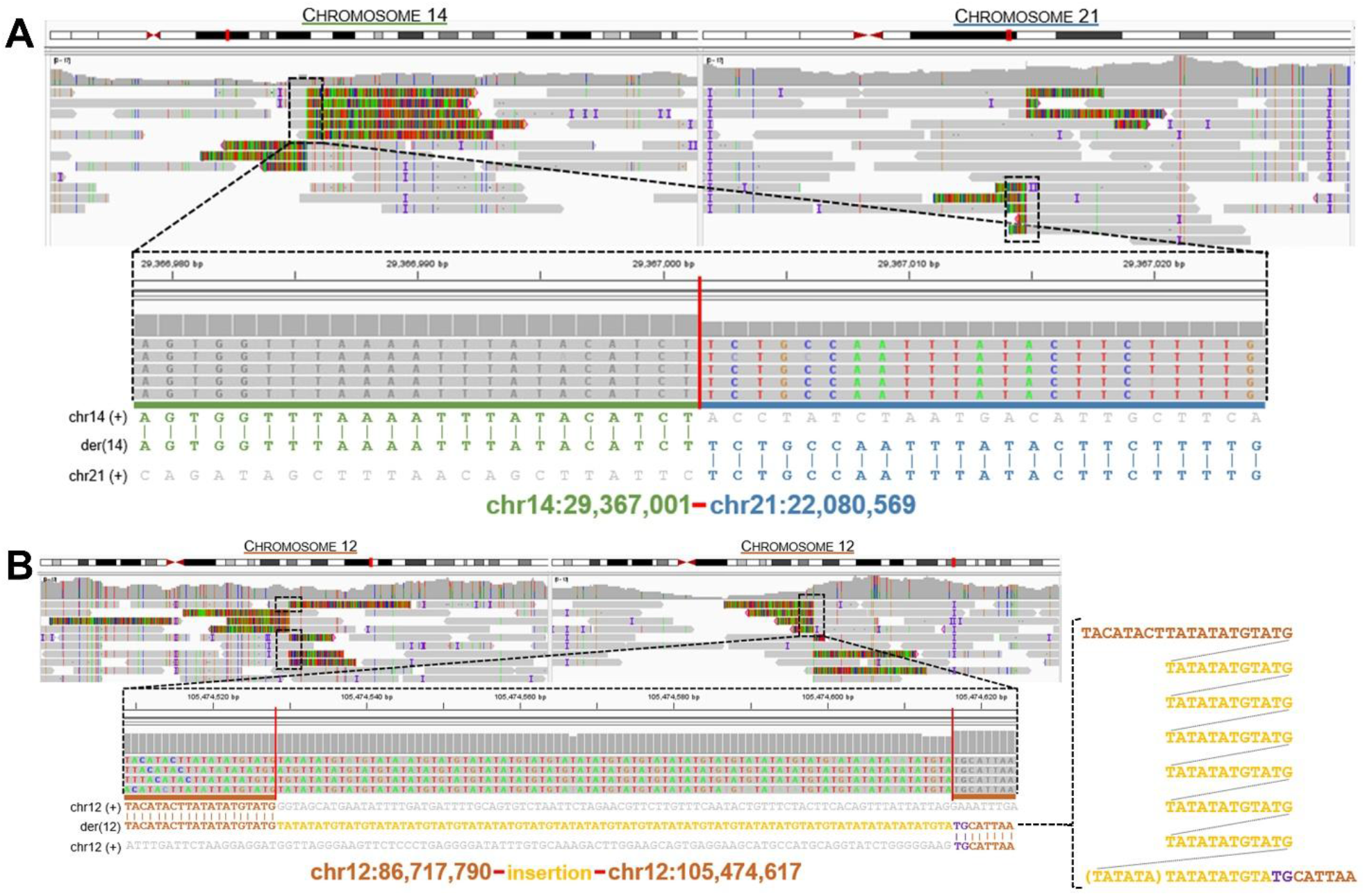
Two of Patient 1’s lrGS junction points seen in chimeric reads through IGV. At the top, IGV view of both involved regions. At the bottom, zoom in on the junction point at the nucleotide level with the sequence alignment and breakpoints below. **(A)** Junction point between chromosomes 14 and 21 in der(14) shows simple ligation of the broken ends through NHEJ with no editing. **(B)** Junction point in der(12) with an insertion between both chromosome 12 regions. To the right, a highlight in the der(12) sequence shows that the insertion corresponds to a repetition of the final 12 nucleotides of the upstream breakpoint. The patient’s other two lrGS breakpoints can be seen in Supplementary Figure 2.

OGM analysis was able to identify eight breakpoints and four junctions, the same as lrGS. The average distance between the actual breakpoint found by lrGS and the one found by OGM was 2,641 (±1,256) base pairs (bp) (Supplementary File 1 – Supplementary Table 5). All but one junction (JP1) presented a shorter sum of the breakpoint difference than the size of the uncertain OGM region.

The patient’s chromosome 4 deletion encompassed five OMIM genes (*NEK1*, *CLCN3*, *HPF1*, *MFAP3L*, and *AADAT*) and was classified as likely pathogenic (class 4) according to the American College of Medical Genetics and Genomics (ACMG) guidelines (Riggs *et al*., 2020). The complex rearrangement led to the rupture of one gene (*MGAT4C*) in chromosome 12 as well as regulatory elements’ interactions of the *PRKD1* gene in chromosome 14.

### Patient 2

Patient 2 was a 14-year-old boy and the only child of a non-consanguineous couple. The patient presented multiple congenital alterations, including neurodevelopmental delay, speech delay, dandy-walker malformation, hypoplasia of the corpus callosum, bell-shaped thorax, decreased calvarial ossification, sclerosis of skull base, abnormal vena cava physiology, dilatation of the renal pelvis, retinal thinning, bilateral conductive hearing loss, and facial dysmorphisms, such as high and prominent forehead, large fontanelles, proptosis, hypertelorism, low-set ears, neck muscle hypoplasia, and dysphagia (Supplementary File 2 – Supplementary Table 1).

The patient was initially investigated through cytogenomic techniques when he was a newborn (Guilherme *et al*., 2013). Karyotyping revealed a complex rearrangement involving four chromosomes: 3, 6, 8, and 14 (Figure 3A), and CMA showed a 585 kb deletion on chromosome 14q24.1 (Supplementary File 2 – Supplementary Table 2, Supplementary Figure 1). By employing FISH with different probes, as described in Guilherme et al, 2013, the rearrangement was identified as involving four chromosomes, nine breakpoints, one insertion, and one deletion. Both of his parents presented normal results in karyotyping.

**Figure 3.**
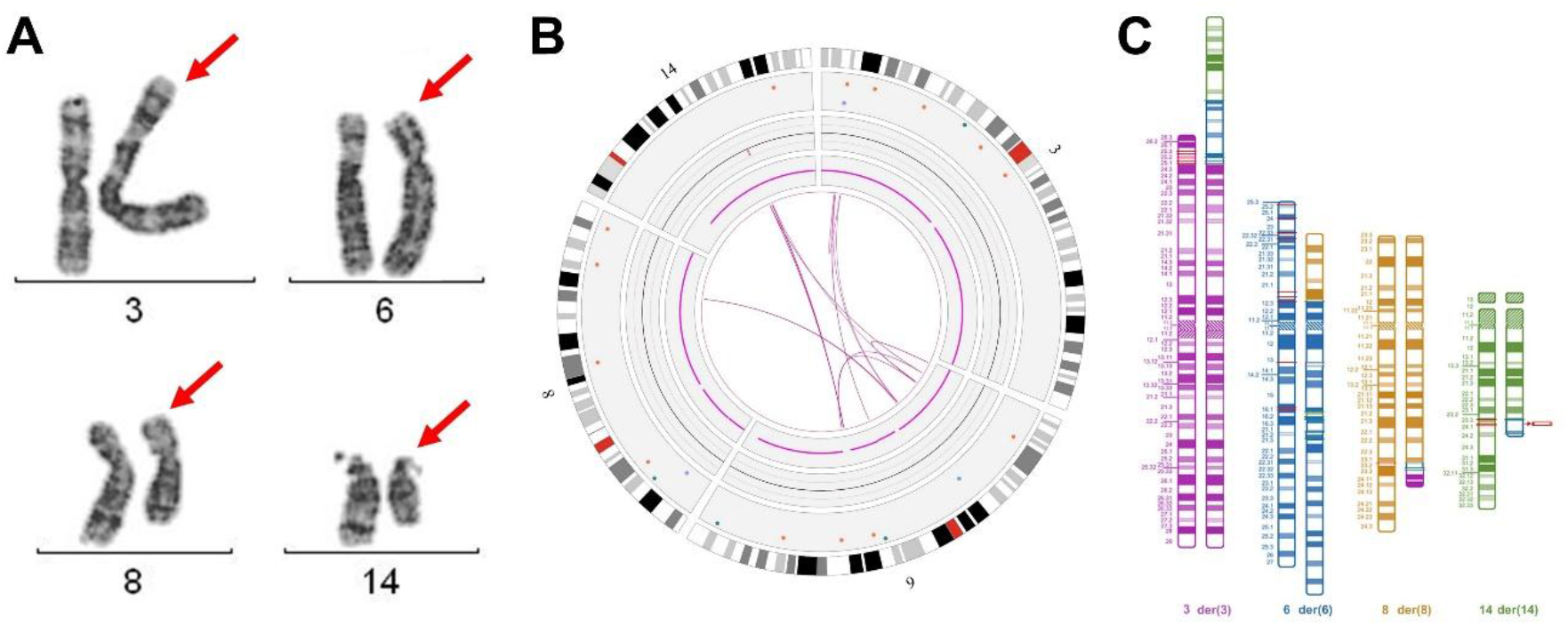
Patient 2’s results. (A) Partial karyotype showing the chromosomes involved in the CCR. Red arrows show the der(3), der (6), der(8), and der(14). **(B) Circos plot showing chromosomes 3, 6, 8, and 14.** The chromosomes’ idiograms can be seen in the periphery while the pink lines in the middle represent the intra- and inter-fusions. **(C) Idiogram of the chromosomes involved in the complex chromosomal rearrangement.** Red lines show the breakpoints in the normal chromosomes and cyan lines show the junction points in the derivative chromosomes. Patient 2 presents 31 breakpoints and 18 junctions.

OGM showed a 757 kb deletion and revealed many intra- and inter-fusions between the four chromosomes discovered by the previous techniques (Figure 3B; Supplementary File 2 – Supplementary Table 2, Supplementary Figure 2), providing an approximate location of 13 breakpoints. lrGS confirmed the involvement of the same four chromosomes, 3, 6, 8, and 14, and the combination of visual inspection between OGM and lrGS allowed us to completely resolve the rearrangement. The patient presents a CCR involving four chromosomes, 31 breakpoints, and 18 junction points, as well as a 592 kb del(14) and a 1.8 kb del(3) (Figure 3C; Supplementary File 2 – Supplementary Table 3, Supplementary Table 4, Supplementary Figure 3). Half of the junction points exhibited microhomology, three showed 1-3 nucleotide insertions and six presented no mechanistic information scars (Supplementary File 2 – Supplementary Table 4). Both parents were investigated through lrGS and presented normal results, therefore the patient’s CCR is *de novo*.

OGM analysis showed 31 breakpoints and 18 junctions, the same as lrGS. However, it is important to note that the OGM detection of some of them relied on visual inspection and interpretation. Four breakpoints were detected as start/end of CNVs – JP4 (fus(3).1) was a duplication and JP5 (fus(3).2) was a deletion – while the two breakpoints of JP18 were called as start/end of an inversion. The average distance between the OGM breakpoints and the lrGS breakpoints was 5,989 (±6,920) bp (Supplementary File 2 – Supplementary Table 5). The sum of the breakpoint difference was shorter than the size of the uncertain OGM region for 11/18 junctions, indicating that these breakpoints were indeed within the uncertain region, as expected.

The patient’s chromosome 14 deletion overlaps four OMIM genes (*PIGH*, *RDH11*, *RDH12*, and *ZFYVE26*) and was classified as pathogenic (class 5) according to the ACMG guidelines (Riggs *et al*., 2020). The chromosome 3 deletion involves two OMIM genes (*SYN2* and *TIMP4*) and was classified as a variant of uncertain significance (VUS) (class 3) according to the ACMG guidelines (Riggs *et al*., 2020).

### Patient 3

Patient 3 was a 6-year-old boy and the second child of a non-consanguineous couple, who also has an older healthy daughter. The patient presented with neuropsychomotor developmental delay, speech delay, irritability, genu valgum, laryngomalacia, atopic dermatitis, poor eye contact, brachycephaly, plagiocephaly, and facial dysmorphisms, including triangular face, prominent forehead, facial edema, long eyelashes, epicanthal folds, anteverted nostrils, and low-set ears (Supplementary File 3 – Supplementary Table 1). Both parents presented normal genomic results.

Patient 3 exhibited a difficult karyogram to analyze due to a complex rearrangement likely involving five chromosomes: 1, 3, 9, 11, and 15 (Figure 4A; Supplementary File 3 – Supplementary Table 2, Supplementary Figure 1). Despite this, CMA revealed only a ∼2.2 Mb deletion in chromosome 11q23.3 (Supplementary File 3 – Supplementary Table 2). OGM corroborated with the CCR and actually uncovered the involvement of chromosome 10 as well as the other five previously discovered, resulting in a six-chromosome rearrangement (Figure 4B; Supplementary File 3 – Supplementary Figure 2). The technique also called two inversions in chromosome 3 and one in chromosome 9, one duplication in chromosome 3, one deletion in chromosomes 3 and the deletion in chromosome 11 (Supplementary File 3 – Supplementary Table 2).

**Figure 4.**
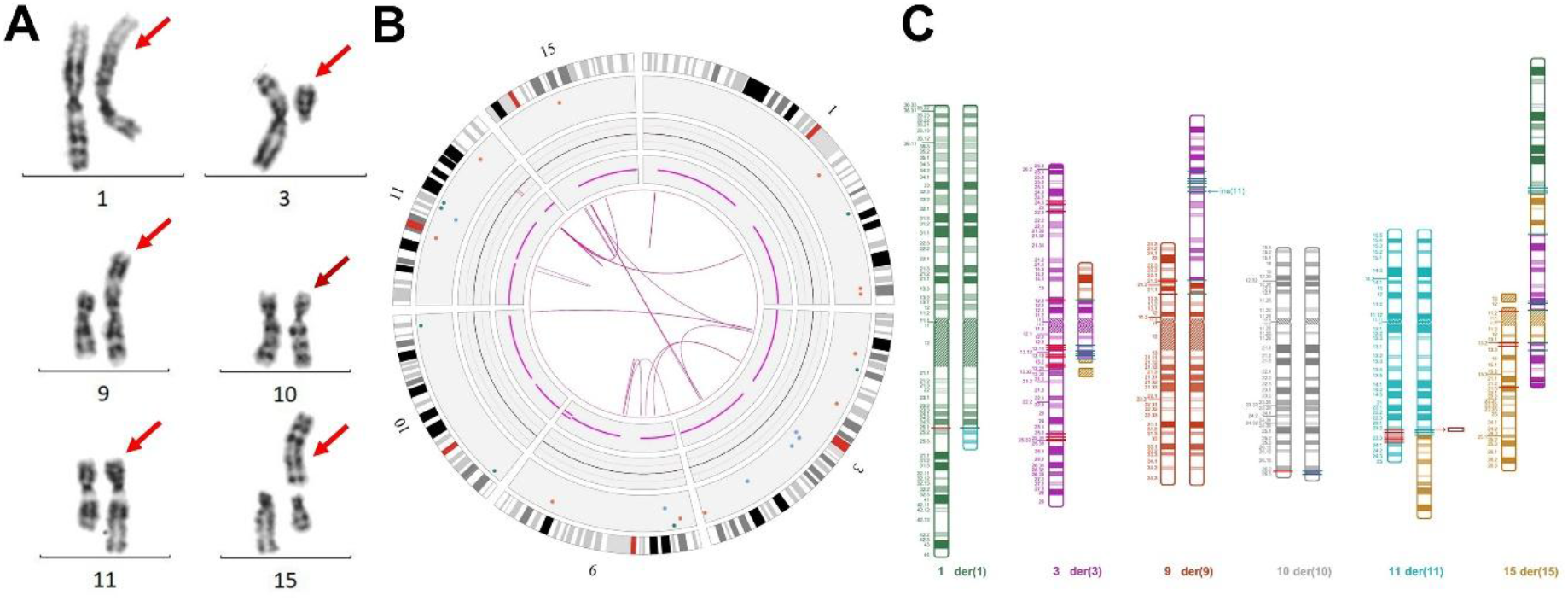
Patient 3’s results. (A) Partial karyotype showing the chromosomes involved in the CCR. Red arrows show the der(1), der(3), der(9), der(11), and der(15). Burgundy arrow show the der(10), which was identified as affected through OGM. **(B) Circos plot showing chromosomes 1, 3, 9, 10, 11, and 14.** The chromosomes’ idiograms can be seen in the periphery while the pink lines in the middle represent the intra- and inter-fusions. **(C) Idiogram of the chromosomes involved in the complex chromosomal rearrangement.** Red lines show the breakpoints in the normal chromosomes and cyan lines show the junction points in the derivative chromosomes. Patient 2 presents 42 breakpoints and 28 junctions.

LrGS analysis confirmed the complexity of the rearrangement by presenting multiple chimeric reads throughout the six involved chromosomes. The complete resolution of the CCR relied on a combined visual analysis of OGM and lrGS results (aligned against references GRCh38/hg38 and T2T-CHM13) as well as a return to the karyotyping result to properly assemble all derivative chromosomes. T2T-CHM13-alignment and karyotyping were vital for the resolution since the first showed the involvement of an acrocentric chromosome short arm and later identified it as chromosome 15p. The final CCR involves 42 breakpoints and 28 junctions, which were all characterized (Figure 4C; Supplementary File 3 – Supplementary Table 3, Supplementary Table 4, Supplementary Figure 3). The majority of the junctions exhibited no information scars at their junctions, while seven junctions showed microhomology and one involved an insertion of chromosome 11 between both breakpoints of chromosome 3 (JP12). The CCR was classified as *de novo* since neither parent presented genetic alterations when investigated through lrGS.

Since rearrangement resolution relied on both OGM and lrGS, more breakpoints and junctions were discovered in OGM after their finding in lrGS. As a result, OGM presented 30 breakpoints and 15 junctions while lrGS detected 42 breakpoints and 28 junctions. Importantly, OGM analysis required considerable interpretation, with lrGS assistance, given that six junctions were called as CNVs, and three inter-fusions were not called and required chimeric map investigation, which also happened for seven intra-fusions. The involvement of chr15p was also not detected since the OGM result was not aligned against T2T. The average distance between the OGM breakpoints and the lrGS breakpoints was 6,451 (±6,325) bp (Supplementary File 2 – Supplementary Table 5). The majority of junctions showed a smaller sum of the breakpoint difference than the size of the OGM uncertain region. Ten junctions exhibited a great sum of the difference between these two detected breakpoints, which corroborated with the size of the uncertain OGM region. Upon OGM reference map investigation, this is due to large unlabeled regions that hampered the identification of a closer breakpoint.

The patient’s chromosome 11 deletion overlaps ten OMIM morbid genes (*APOA1*, *APOA4*, *APOA5*, *APOC3*, *BUD13*, *CEP164*, *FXYD2*, *IL10RA*, *SIK3*, and *ZPR1*) and was classified as pathogenic (class 5) according to the ACMG guidelines (Riggs *et al*., 2020).

### Patient 4

Patient 4 was a 6-year-old boy and the first child of a non-consanguineous couple, who also had a younger healthy son. The patient presented with neuropsychomotor developmental delay, autism, aggressive behavior, lack of interest in peers, agitation, auditory hypersensitivity, microcephaly, brachycephaly, cerebellar hypoplasia, mega cisterna magna, cryptorchidism (now surgically corrected), and facial dysmorphisms, including sparse and thin eyebrows, widow’s peak, slender nose with long nasal bridge and anteverted hypoplastic nostrils (Supplementary File 4 – Supplementary Table 1).

Patient 4 presented a normal karyotype (46,XY) (Figure 5A; Supplementary File 4 – Supplementary Figure 1), as did his parents, and was further investigated due to his phenotype, which was indicative of a genetic alteration. Chromosomal microarray analysis unveiled a 3.7 Mb triplication in chromosome 2 (arr[GRCh38] 2q37.2q37.3(236068385_239769318)×4) (Supplementary File 4 – Supplementary Table 2). OGM detected a similar triplication in 2q while also revealing that the three copies of this region were in tandem with the middle one being inverted (Figure 5B; Supplementary File 4 – Supplementary Figure 2).

**Figure 5.**
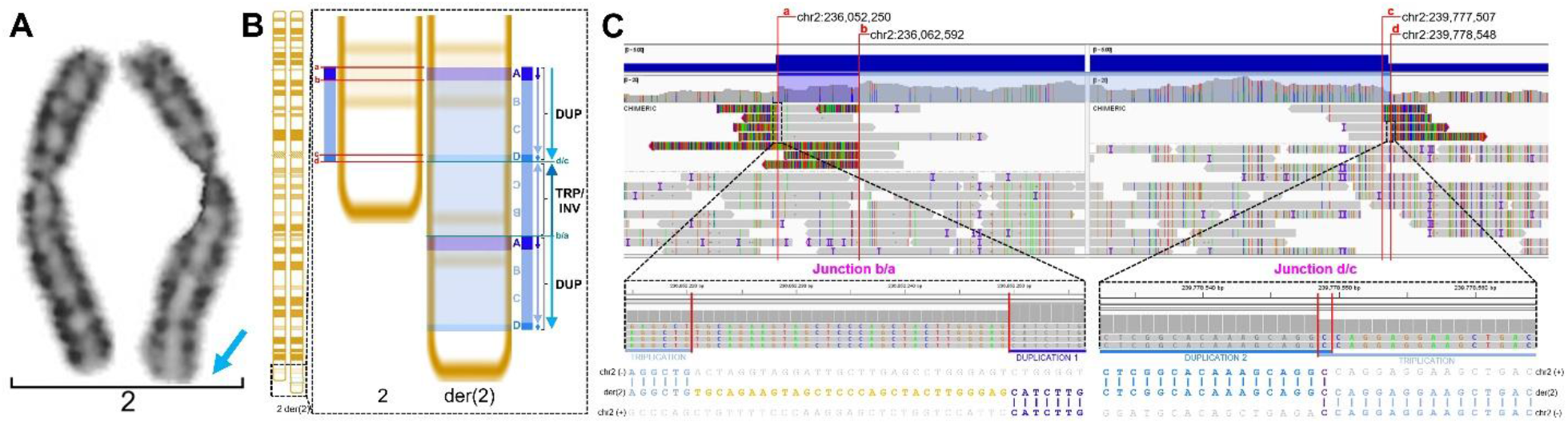
Patient 2’s results. **(A) Partial karyotype showing the chromosome 2 pair.** Initially, a normal karyotype (46,XY) was reported. After the resolution of the rearrangement with the other methodologies, a closer look at the karyotype allowed for the identification of a lighter chromosome band at the end of chromosome 2 that corresponds to the alteration (blue arrow). **(B) Idiogram of the chromosome involved in the complex chromosomal rearrangement.** In the rectangle, to the left, a depiction of the breakpoints’ location in red (a, b, c, and d) and a representation of the first duplication between a and b, the triplication between b and c, and the second duplication between c and d. To the right, the characterization of the final DUP-TRP/INV-DUP rearrangement highlighting, in pink lines, junction d/c between the second duplication and the inverted triplication and junction b/a between the inverted triplication and the first duplication. **(C) LrGS junction points.** At the top, IGV view of the chimeric reads in the regions involved in the CCR. The breakpoints (a, b, c, and d) are indicated with red lines. At the bottom, zoom in on the junction points at the nucleotide level with the sequence alignment and breakpoints below. In junction b/a, a 29-nucleotide insertion is revealed while in junction d/c, a 1-nucleotide microhomology is present.

In the lrGS analysis, two breakpoints were identified upon the inspection of the surrounding region of each breakpoint found in OGM (Supplementary File 4 – Supplementary Table 2, Supplementary Figure 3). A close investigation of this result resolved the patient’s rearrangement: in the 2q37.2q37.3 region, the patient presents a ∼10 kb duplication, a ∼3.7 Mb triplication, and a ∼1 kb duplication (Supplementary File 4 – Supplementary Table 3), leading to the formation of a complex rearrangement known as duplication-inverted/triplication- duplication (DUP-TRP/INV-DUP) (Figure 5B; Supplementary File 4 – Supplementary Figure 4). All breakpoints and both junction points were identified and fully characterized (Supplementary File 4 – Supplementary Table 4, Supplementary Table 5). Junction point 1 (between the end of the second duplication and the start of the inverted triplication) exhibited 1 nucleotide of microhomology between the two regions while junction point 2 (between the end of the inverted triplication and the start of the first duplication) presented an insertion of 29 nucleotides that did not map anywhere in the genome (Figure 5C). Both parents presented normal results in the lrGS analysis, therefore the patient’s CGR is *de novo*.

OGM showed two breakpoints and no junction between them, while lrGS identified four breakpoints forming two junctions. The distance between the OGM and the lrGS breakpoints was 3,962 bp for the upstream breakpoint and 23,281 bp for the downstream one (Supplementary File 4 – Supplementary Table 6).

The patient’s 10 kb duplication overlapped a part of the *AGAP1* (OMIM *608651) gene, while the 3.7 Mb triplication encompassed 21 OMIM genes (*AGAP1, HDAC4, ILKAP, PER2, UBE2F, COL6A3, COPS8, GBX2, ACKR3, LRRFIP1, TRAF3IP1, MLPH, ASB1, ESPNL, RAB17, RAMP1, SCLY, TWIST2, HES6, PRLH,* and *ERFE*), and the 1 kb duplication did not involve any genes. Both duplications were classified as VUS (class 3) and the triplication as likely pathogenic (class 4) according to the ACMG guidelines (Riggs *et al*., 2020).

### Patient 5

Patient 5 was a 7-year-old boy and the second child of a non-consanguineous couple. Both parents presented normal karyotypes, and their first daughter was healthy. The patient presented with seizures usually precipitated by febrile ear infection or gastroenteritis, facial hypotonia with drooling (sialorrhea), neurodevelopmental delay, hypoplasia of the cerebellum, cranial asymmetry, hypoplasia of the frontal lobes, Blake’s pouch cyst, posterior plagiocephaly, abnormal retinal morphology, renal insufficiency, positive CMV urine nucleic acid test, decreased palmar creases, clinodactyly of the 5th finger, fibular deviation of the 3^rd^ and 4^th^ toes, small low-set ears with auricular pit and prominent crus of helix, and facial characteristics, including hypertelorism, downslanted palpebral fissures, depressed nasal bridge, deep philtrum, and thin upper lip vermilion (Supplementary File 5 – Supplementary Table 1).

Karyotype analysis showed that Patient 5 exhibited additional material in chromosome 16: 46,XY,add(16)(q13) (Figure 6A; Supplementary File 5 – Supplementary Figure 1) while his parents present normal results. CMA revealed a complex genomic rearrangement with a ∼514 kb 16p deletion and three 16q duplications of ∼4.7 Mb, ∼6.7 Mb, and ∼11.7 Mb (Supplementary File 5 – Supplementary Table 2). OGM detected four similar CNVs and revealed that the extra copies of the two downstream 16q duplications were in tandem with each original copy and in the same orientation while the extra copy of the first duplication was not localized. Besides that, OGM also uncovered an additional ∼45 kb 16q deletion through the SV pipeline (Figure 6B; Supplementary File 5 – Supplementary Table 2, Supplementary Figure 2).

**Figure 6.**
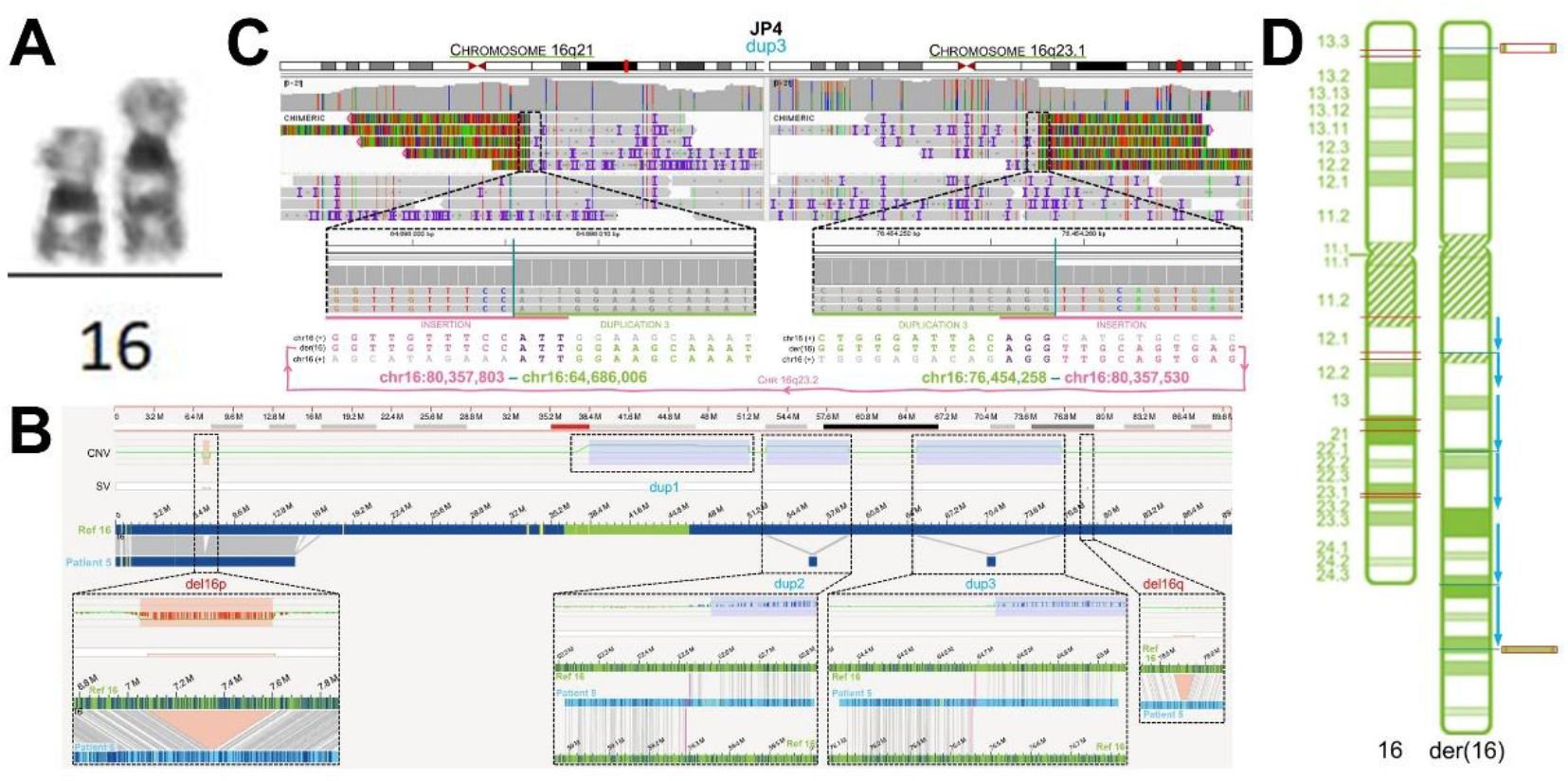
Patient 5’s results. **(A) Partial karyotype showing the chromosome 16 pair.** The der(16) presents additional material in 16q. **(B) OGM genome browser view of chromosome 16.** From top to bottom, chromosome 16 idiogram, CNV call track, SV call track, chromosome 16 reference map, patient maps. Five CNVs were called by the CNV pipeline but only four were identified in the maps and called by the SV pipeline. **(C) Example of lrGS junction point.** At the top, IGV view of the chimeric reads in the regions involved in the junction. At the bottom, zoom in on the junction points at the nucleotide level with the sequence alignment and breakpoints below. This junction refers to the distal duplication of 16q, which presented an insertion of a different region of chromosome 16 in between its breakpoints. **(D) Idiogram of the chromosome involved in the complex chromosomal rearrangement.** To the left, the normal chromosome 16 with breaks shown in pink. To the right, the rearranged chromosome 16 with junctions represented in cyan. Blue arrows indicate the location and orientation of the copies of the duplications and regions in red indicate the deletions. Patient 5 presents 14 breakpoints and five junctions.

The long-read sequencing analysis identified both deletions’ and the three duplications’ breakpoints at the nucleotide level and characterized all five junctions (Figure 6C; Supplementary File 5 – Supplementary Table 3, Supplementary Table 4, Supplementary Figures 3-7). Similarly to OGM, lrGS confirmed the location and orientation of the two downstream 16q duplications. Alignment to the GRCh38/hg38 reference genome did not reveal this information for the proximal 16q duplication, however, using the T2T-CHM13, lrGS found the same pattern for all three duplications (Figure 6D; Supplementary File 5 – Supplementary Table 5, Supplementary Figure 8). Junction 4 and Junction 5 presented higher complexity revealed by the higher resolution analysis, leading to the formation of four subjunctions due to the insertion of other regions of chromosome 16 in between the original breakpoints. All junctions, beside JP2, present microhomology between the breakpoints (Supplementary File 5 – Supplementary Table 4). Patient 5’s parents were not available for lrGS investigation.

OGM detected 10 breakpoints and five junctions, while lrGS uncovered four more breakpoints, totaling 14 breakpoints present in five junctions. The average difference between the breakpoints found by OGM and the ones found by lrGS was 7,832 (±7,002) bp (Supplementary File 5 – Supplementary Table 6). All breakpoints fell within the OGM uncertain region of their respective junction.

The patients’ ∼509 kb 16p deletion, ∼4.8 Mb, ∼6.7 Mb, and ∼11.7 Mb 16q duplications, and 33kb 16q deletion together encompass over 70 OMIM morbid genes.

### Patient 6

Patient 6 was an 11-year-old boy and the only child of a non-consanguineous couple. The patient presented with mild intellectual disability, autism, and epilepsy (Supplementary File 6 – Supplementary Table 1). Karyotype analysis revealed additional material in chromosome 18: 46,XY,add(18)(p11.2) (Figure 7A; Supplementary File 6 – Supplementary Figure 1). Investigation into the parents’ karyotype showed a normal result for the mother and a rearranged chromosome 18 for the father: 46,XY,der(18)?add(18)(p11.2)?add(18)(q11.2). CMA of Patient 6 revealed an 8.6 Mb duplication and a 6.7 Mb deletion in 18q (Supplementary File 6 – Supplementary Table 2). His father’s CMA results exposed the occurrence of 11 microduplications across chromosome 18, ranging from 26 kb to 191 kb (Supplementary File 6 – Supplementary Table 3). Unfortunately, the father is absent from in his son’s life, therefore it was not possible to continue studying his genome. According to information given by the patient’s paternal grandmother, the patient has a younger half-sister who presents altered phenotypes (uninformed), and his aunt’s family is also likely affected, since his cousin also presents autism.

**Figure 7.**
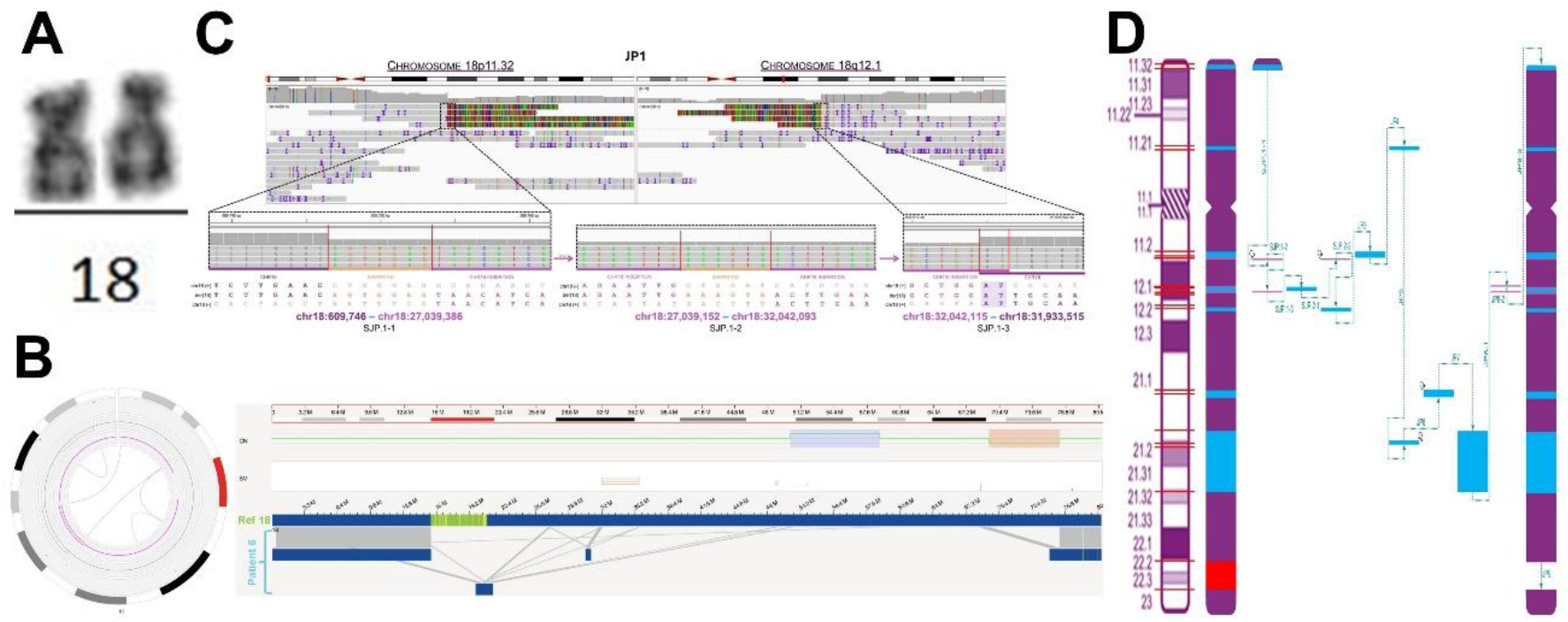
Patient 6’s results. **(A) Partial karyotype showing the chromosome 18 pair.** The der(18) presents additional material. **(B) OGM results.** To the left, circos plot showing chromosome 18’s idiogram in the periphery with pink lines in the middle representing the intra-fusions. To the left, genome browser view of chromosome 18. From top to bottom, chromosome 18 idiogram, CNV call track, SV call track, chromosome 18 reference map, and patient maps. The CNV pipeline detected the large 16q duplication and deletion while we can observe each patient maps mapping to various regions of the chromosome. **(C) Example of lrGS junction point.** At the top, IGV view of the chimeric reads in the regions involved in the junction. At the bottom, zoom in on the junction points at the nucleotide level with the sequence alignment and breakpoints below. This junction presents the insertion of two different regions of chromosome between its breakpoints, thus forming three subjunctions. **(D) Idiogram of the chromosome involved in the complex chromosomal rearrangement.** To the left, the normal chromosome 18 has the breaks shown in pink. To the left, a schematic representation of how the fragments rearranged to form the patient’s der(18). Dotted cyan lines represent junctions and subjunctions, blue regions refer to duplications and microduplications while the red one show the deletion. Patient 6 presents 28 breakpoints, nine junctions, and eight subjunctions.

OGM detected both of the patient’s previously known 18q CNVs as well as a 199.6 kb 18q11.2 duplication, 3.5 Mb 18q12.1q12.2 deletion, a 276 kb 18q21.1inversion, and six fusions involving both 18p and 18q (Figure 7B; Supplementary File 6 – Supplementary Table 2, Supplementary Figure 2). LrGS uncovered the CGR as involving 28 breakpoints and nine junctions, with an extra eight subjunctions due to the insertion of other regions of chromosome 18 (Supplementary File 6 – Supplementary Table 4, Supplementary Table 5). Seven junctions presented insertions, three of regions in chromosome 18 (Figure 7C; Supplementary File 6 – Supplementary Figure 3) and four of nucleotides that did not map anywhere in the genome. The latter was also identified in five subjunctions. Microhomology was detected in one junction and one subjunction. Based on this, the technique revealed 12 microduplications in Patient 6’s chromosome 18, of which nine overlap with his father’s microduplications (Supplementary File 6 – Supplementary Table 6, Supplementary Figure 4). The complete resolution of the CGR was only achieved by the association of OGM and lrGS visual inspections (Figure 7D; Supplementary File 6 – Supplementary Figure 5). The patient’s mother was investigated through lrGS and presented normal results.

OGM identified 16 breakpoints and nine junctions, while lrGS revealed 28 breakpoints, forming the nine junctions. The average difference between the breakpoints found by OGM and the ones found by lrGS was 5,290 (±2,317) base pairs (Supplementary File 6 – Supplementary Table 7). Some junctions presented a large difference between the sum of the difference of their breakpoints and the size of the uncertain OGM region, while one had the exact same value.

Patient 6’s 8.7 Mb 18q duplication and 6.7 Mb 18q deletion overlap nine and four OMIM genes, respectively, and were both classified as pathogenic (class 5) according to the ACMG guidelines (Riggs *et al*., 2020). Concerning the 12 microduplications, classified as VUS (class 3), two OMIM genes are duplicated and two morbid OMIM genes are ruptured.

## DISCUSSION

In six patients, three CCRs and three CRGs were initially suggested from classical cytogenetic assessment, including karyotyping and CMA. Here, the combination of OGM and lrGS allowed us to completely characterize all patients’ complex rearrangements to breakpoint resolution, thus enabling the inference of their mechanisms of formation and the correlation of precise breakpoints with the phenotype.

### Mechanisms of Formation

Complex rearrangements can be divided into four groups based on the number of breakpoints that they present compared to the quantity of chromosomes involved (Madan, 2012; Aynaci *et al*., 2025; Kocagil *et al*., 2026). Type I rearrangements are the most common and possess the same number of breakpoints and chromosomes involved, presenting as reciprocal translocations where each chromosome harbors one breakpoint. Type II rearrangements are more complicated, with a greater number of breakpoints than chromosomes involved, usually due to inverted segments. Type III and Type IV rearrangements also show more breakpoints than involved chromosomes; however, they are formed due to a massive disruptive cellular event that leads to multiple structural rearrangements in one or more chromosomes with numerous breakpoints and severe phenotypical alterations. The difference between them is that, while Type III rearrangements commonly exhibit insertions, Type IV rearrangements present fragments of other chromosomes in at least one derivative chromosome, therefore being the most complex type (Madan, 2012; Aynaci *et al*., 2025; Kocagil *et al*., 2026). Given the characterization of the six complex rearrangements described here, they all fall into Types III and IV, with the chromoanagenesis mechanisms likely being responsible for their formation.

The term chromoanagenesis contemplates three previously described mechanisms: chromothripsis, chromoanasynthesis, and chromoplexy (Figure 8) (Pellestor and Gatinois, 2018; Zepeda-Mendoza and Morton, 2019; Burssed *et al*., 2022). Chromothripsis involves a single chromosome, which can be encapsulated inside a micronucleus, where it breaks into small fragments, and is then reassembled through NHEJ in a different order or orientation than the original one with the possibility of loss of fragments, leading to deletions. Chromoanasynthesis also comprises a single chromosome but, in this case, it relies on FoSTeS or MMBIR for re-synthesis of DNA segments to form a new rearranged chromosome, which can present deletions, duplications, inversions, and triplications. Chromoplexy is the only one that encompasses two or more chromosomes, which undergo a chain of rearrangements consisting of DNA segments from distinct chromosomes breaking and rejoining through c-NHEJ or MMEJ, leading to the formation of usually balanced complex translocations. Importantly, chromoplexy creates fewer breakpoints than chromothripsis. The rearrangements of the patients in this study can have one (or two) of these three mechanisms as responsible for their formation; however, some of them present characteristics that do not fit entirely with these mechanisms’ known description.

**Figure 8.**
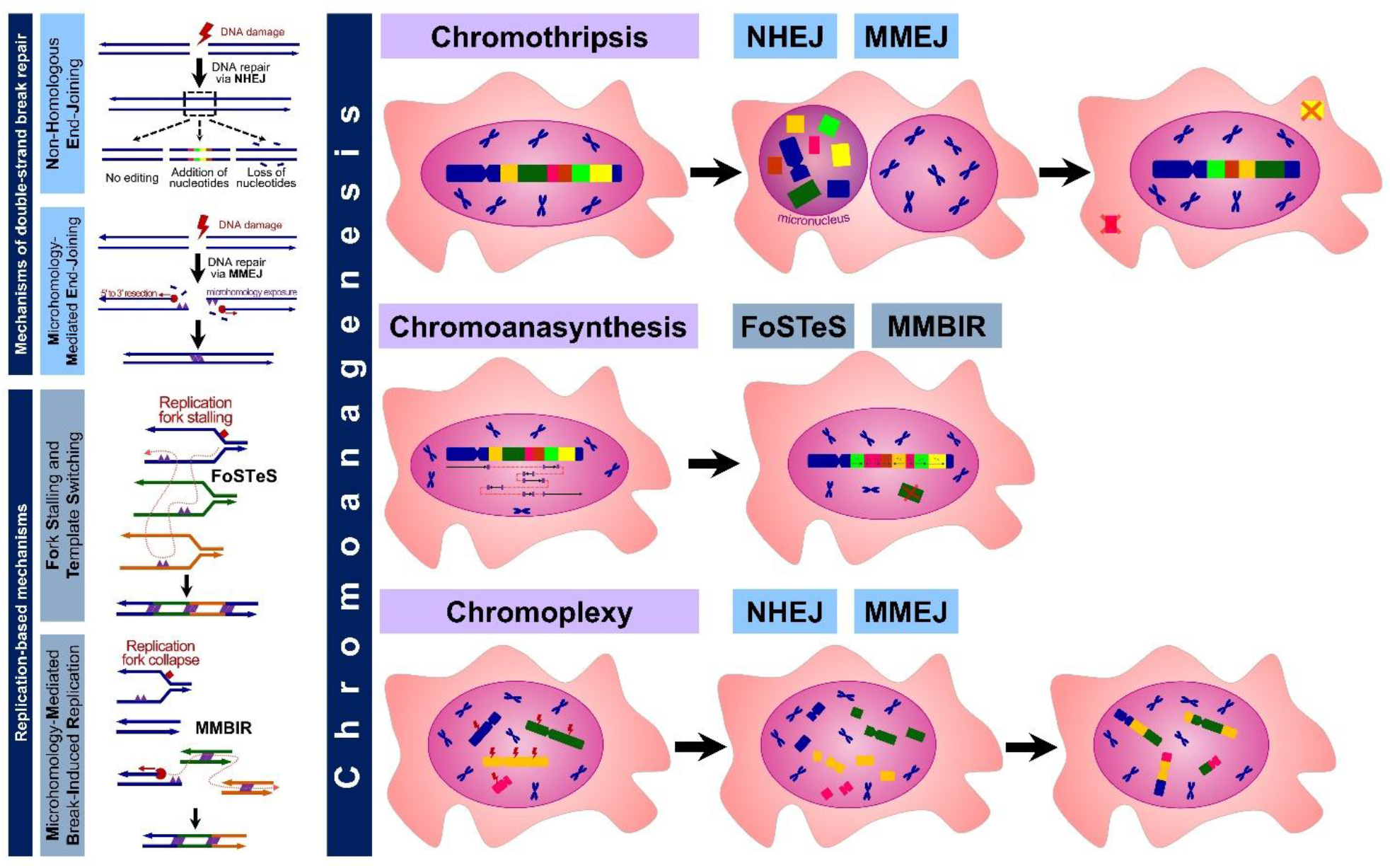
Mechanisms of formation of complex rearrangements. **To the left, at the top, mechanisms of double-strand break repair.** Non-Homologous End-Joining (NHEJ) repairs DSBs by joining the broken extremities with or without editing, such as addition or loss of nucleotides. Microhomology-Mediated End-Joining (MMEJ) is a similar mechanism but it relies on a 5’-3’ resection to expose nucleotides of microhomology between the broken ends to ease their rejoining. **To the left, at the bottom, replication-based mechanisms.** Fork Stalling and Template Switching (FoSTeS) happens after a replication fork stalls and the lagging DNA strand invades other active replication forks due to microhomology and continues DNA replication. Microhomology-Mediated Break-Induced Replication (MMBIR) occurs when a replication fork breaks and a 3′ single-stranded overhang invades other genomic regions also due to microhomology. **To the right, the chromoanagenesis mechanisms.** Chromothripsis involves one chromosome that is encapsulated in a micronucleus, where it breaks into multiple fragments that rejoin through NHEJ/MMEJ, forming a new chromosome that can present deletions. Chromoanasynthesis also involves a single chromosome that is subjected to DNA resynthesis through FoSTeS/MMBIR, forming a new rearranged chromosome with microhomology at the junctions as well as deletions, inversions, duplications, and triplications. Chromoplexy involves more than two chromosomes that undergo a chain of breaking and joining their DNA segments through NHEJ/MMEJ, forming usually balanced complex translocations.

Patient 1 presented a complex chromosomal rearrangement between chromosomes 12, 14, and 21. The junction points from der(14) and der(21) suggest that the broken ends were simply joined with no editing, indicating that they were joined through the NHEJ mechanism. Since fragments of three different chromosomes joined together to form complex translocations, we can infer the mechanism of chromoplexy for the formation of these two derivative chromosomes (Pellestor, 2019; Burssed *et al*., 2022). The der(12), meanwhile, appears to be formed due to backward replication slippage, in which, during DNA replication, the leading strand dissociates from the template, slips backward, and replicates a region more than it should, leading to the insertion of a repetitive sequence at the junction point (Ohye *et al*., 2014; Hansson, 2018). In this case, the final 12 nucleotides of the upstream breakpoint were replicated repetitively before joining with the downstream breakpoint. All breakpoints are located in regions that show the presence of repetitive sequences of the genome (Supplementary File 1 – Supplementary Table 4). Notably, the downstream breakpoints of chromosome 12 and the breakpoints of chromosome 14 happen within LINE elements, being from families L2 and L1, respectively. Repetitive sequences have been associated with rearrangements formed through NHEJ therefore their presence at and around the breakpoints may have predisposed these regions to rearrange (Shaw and Lupski, 2005; Gu, Zhang and Lupski, 2008; Burssed *et al*., 2022).

Patients 2 and 3 present CCRs involving four (3, 6, 8, and 14) and six (1, 3, 9, 10, 11, and 15) chromosomes, respectively. As with Patient 1, chromoplexy immediately appeared as a plausible mechanism for the rearrangements’ formation, given that they involve multiple chromosomes. However, unlike the previous patient, these patients present CNVs and a much larger number of breakpoints when compared to the number of chromosomes involved. Patient 2 harbors 31 breakpoints and Patient 3 present 42 breakpoints. Both show junctions of broken ends with no editing, indicating the NHEJ mechanism, but also present a considerable number of junctions exhibiting microhomology. Given these characteristics, we proposed a different mechanism for their formation that relies on certain points of each chromoanagenesis mechanism.

Since these rearrangements involve multiple chromosomes, numerous breakpoints, and are constant throughout the cells, a micronucleus could have been formed early in development and encapsulated all of the involved chromosomes due to lagging chromosomes or after mitotic errors or DNA damage (Kwon, Leibowitz and Lee, 2020). Inside the micronucleus, additional DNA breaks probably happened across all chromosomes and, afterwards, the breakpoints joined amongst themselves, driven by the presence of REs and microhomology, mainly through NHEJ and MMEJ, but also possibly through replication-bases mechanisms. This led to the formation of the complex derivative chromosomes presenting several intra-fusions and inter-fusions amongst only the micronucleus-encapsulated chromosomes as well as deletions, duplications, and insertions. The micronucleus formation would explain why only these chromosomes are involved in the rearrangement and how the fragments formed due to the same breaks are exchanged between all of them. This proposed mechanism combines the involvement of multiple chromosomes from chromoplexy, the micronucleus from chromothripsis, the NHEJ and MMEJ mechanisms from both, and the CNVs and insertions from chromoanasynthesis.

Patients 2 and 3 exhibited REs surrounding all breakpoints, with the majority being Alu (SINE) and L1 (LINE) elements (Supplementary File 2 – Supplementary Table 4). Patient 2 had seven junctions with REs of the same class and family, eight with REs of only the same class, and three with no such similarities, but with breakpoints in regions full of REs. For Patient 3, 13 junctions showed REs of the same class and family, eight had REs of only the same class, five had no such similarities, but presented breakpoints in regions full of REs, and two involved chromosome 15p (Supplementary File 3 – Supplementary Table 4).

Patient 4 presents a duplication-inverted/triplication-duplication rearrangement, which was first described by Carvalho et al. (2011). Contrary to what may be perceived at first glance, this rearrangement produces only two junction points (d/c and b/a in Figures 5B,C), where the inverted triplication joined with each duplication in a two-step process in which a replication-based mechanism that involves template switches associates with NHEJ (Carvalho *et al*., 2011, 2015; Grochowski *et al*., 2024). In this patient, we infer that, first, a single-ended double-strand break was formed either after a double-strand break or a replication fork collapse. A 5’-3’ resection created a 3’ single-stranded overhang with exposed microhomology that folded back and intrastrand paired with itself at the site of the microhomologies (1 nucleotide in this case), forming junction point 1 through the Fold-back mechanism (d/c). DNA synthesis was restarted in an inverted orientation until another replication fork collapse happened and created a new 3’ single-stranded overhang that then switched to the sister chromatid, thus forming junction point 2 through NHEJ with an insertion of 29 nucleotides (b/a) and resumed DNA synthesis in the direct orientation until the end of the chromosome (Figure 4C). The Fold-back mechanism has been linked with the formation of inverted duplications associated with terminal deletions (inv-dup-del) that contain a disomic spacer between the two copies of the duplicated segment (Hermetz *et al*., 2014; Burssed *et al*., 2023) however, here, it fits well to explain the first extra copy of the triplicated region. Instead of the DNA replication continuing and then forming a dicentric chromosome, a different path is followed to duplicate/triplicate the regions and continue until the end of the chromosome, therefore not forming a deletion. The DUP-TRP/INV-DUP has been widely reported in chromosome X as a mechanism for diseases such as Pelizaeus-Merzbacher disease (OMIM #312080), Duchenne Muscular Dystrophy (OMIM #310200), and in 20% of the *MECP2* copy number gains in the *MECP2* duplication syndrome (OMIM #300260) (Carvalho *et al*., 2011, 2015; Dittwald *et al*., 2013; Beck *et al*., 2019; Grochowski *et al*., 2024). These rearrangements in the X chromosome are mediated by inverted repeats, low copy repeats, and repetitive elements located at the breakpoints and junctions, thus forming almost recurrent rearrangements (Carvalho *et al*., 2011), whereas for DUP-TRP/INV-DUP rearrangements in autosomes (which are sometimes associated with absence of heterozygosity), these structures are not always present (Carvalho *et al*., 2015, 2019). The genomic architecture of chromosome X is hypothesized to play a role in this difference (Carvalho *et al*., 2015). One breakpoint of patient 4 (b) overlaps with an Alu element but no other repetitive sequences seem to be present to mediate the rearrangement. Grochowski et al. (2024) found four different haplotype sub-structures of DUP-TRP/INV-DUP rearrangements formed due to inverted SDs in the *MECP2* locus. These structures differ amongst themselves with regard to which segments are actually inverted in the final rearrangement. Even though our patient’s rearrangement formation does not involve SDs, we were able to identify that he presents the haplotype structure 4 since only the triplication is inverted.

Patient 5 presents a complex genomic rearrangement in chromosome 16 with one deletion in 16p, one in 16q, and three large duplications, with all of which having the extra copy in tandem of the original sequence in direct orientation. In a detailed analysis of duplicated genomic sequences in this chromosome, Martin et al. (2004) (Martin *et al*., 2004) identified that 9.89% of chromosome 16 is made up of segmental duplications, which is higher than the genomic average (5.3%), therefore it is among the most SD-enriched chromosomes. Given that its intrachromosomal SDs are longer and have higher sequence identity than the interchromosomal SDs (Martin *et al*., 2004), we can infer that these segmental duplications and their higher complexity may play a role in the formation of the patient’s rearranged chromosome 16, which presents five junction points and 14 breakpoints. REs overlapping and surrounding all but one breakpoint may have also favored the rearrangement. *Alu* elements may have contributed to the 16p deletion formation; the Human Satellite II, a complex region formed by extremely repetitive pericentromeric DNA, features in the proximal 16q duplication breakpoint; SINE elements are reported for the middle 16q duplication; and JP4 and JP5 presented more complexity with the insertion of 277 and 96 nucleotides, respectively, mapping to downstream 16q regions, with three subjunctions exhibiting microhomology. Mobile elements have been associated with the formation of non-recurrent rearrangements through NAHR (Burwinkel and Kilimann, 1998; Shaw and Lupski, 2005; Luo *et al*., 2011; Burssed *et al*., 2022). Given that most of the breakpoints are in regions containing SINE elements, we can infer that these may play a role in the formation of patient 5’s complex chromosome 16 likely through chromoanasynthesis, since it involves a single chromosome, duplications, and microhomology.

Patient 6 presents the most complex genomic rearrangement with 28 breakpoints, nine junctions, and eight subjunctions. He also presents a large duplication and a large deletion in 18q, but also 12 microduplications across the chromosome. Interestingly, his father presents 11 micro duplications in the same chromosome. Given this striking similarity, it is acceptable to infer that the father’s genetic alterations may have led to his son’s alterations. Since the father is unfortunately not available for study, we are not able to resolve his rearranged chromosome 18 to completely understand how it may have led to his son’s CGR. However, according the available CMA data, we observe that four microduplications of Patient 6 overlap with his father’s, one encompasses two of his father’s, and four of the father’s microduplications are located in all large CNVs breakpoint regions. In light of this, we infer that the son may have inherited some of his father’s microduplications while others may have created a condition that favored the rearrangement. The latter could have led to the formation of the large CNVs with the microduplications acting as REs or SDs for NAHR to occur between them. Chromoanasynthesis may have happened in the father or at some point in his family, since reports indicate other affected relatives, and in Patient 6, since both of them present various microduplications, insertions, and inversions, which would be consistent with the mechanism. The presence of actual REs reported in and near the breakpoints cannot be discarded as irrelevant for the CGR formation, since they could be bringing together different regions to rearrange.

### Karyotype-phenotype correlation

Patient 1 presents five deleted genes in the likely pathogenic 2.4 Mb chromosome 4 deletion. Among them, the *CLCN3* (*Chloride Voltage-Gated Channel 3*, OMIM *600580) gene is dosage sensitive (pLI = 1.00, LOEUF = 0.34) and has been associated with neurodevelopmental disorder with hypotonia and brain abnormalities (OMIM #619512). The gene encodes the ClC-3, which is a chloride (Cl^-^) channel/transporter located in the vesicles of the endosomal/lysosomal system that mediates 2Cl^-^/H^+^ exchange (Duncan *et al*., 2021). Duncan et al. (2021) reported 11 patients with variants in the *CLCN3* gene that present deleterious impact. These patients present similar phenotypes to patient 1, such as neuropsychomotor developmental delay, microcephaly, plagiocephaly, hypoplasia of the corpus callosum, hypotonia, strabismus, and prominent forehead. The authors highlight that both loss and gain of ClC-3 function affect the nervous system with overlapping possible phenotypes, therefore our patient’s deletion of the gene likely decreases the channel’s function and affects her phenotype. The break in chromosome 12 to form the CCR ruptured the *MGAT4C* (*MGAT4 Family Member C*, OMIM *607385) gene. This gene has been associated with neurocognitive disorders in Mexican individuals (Bliskunova *et al*., 2021) and it has been suggested that genes located in the same chromosome band may play an important role in cognition development (Akilapa, Smith and Balasubramanian, 2015; Bliskunova *et al*., 2021), therefore disruption of this gene may have contributed to the patient’s intellectual disability. The remaining breaks that form the CCR do not rupture any genes. However, the break in chromosome 14 ruptures the interaction between enhancers and the *PRKD1* (*Protein Kinase D1*, OMIM *605435) gene, which is associated with congenital heart defects and ectodermal dysplasia (OMIM #617364), which includes phenotypic features that overlap with our patient’s, such as microcephaly, prominent forehead, widely spaced teeth, hypotonia, global developmental delay, and dysmorphic features. It is therefore intriguing to speculate that the patient presents a composite phenotype due to the disrupted function of at least two genes: *CLCN3* and *MGAT4C*. In order to prove the regulatory defect, a follow-up is planned to look into the impact on methylation and gene function with RNA studies as well as possibly other omics analyses since a multi-omics approach has been proven to resolve complex phenotypes (Vollger et al., 2023).

Patient 2 presents four OMIM genes deleted in the 595 kb chromosome 14 deletion and two in the 1.8 kb chromosome 3 deletion. Two of the chromosome 14 deleted genes are retinol dehydrogenases expressed in the retina which share 73% sequence identity in humans: *RDH11* (*Retinol Dehydrogenase 11*, OMIM *607849) and *RDH12* (*Retinol Dehydrogenase 12*, OMIM *608830) (Kanan *et al*., 2008). Patient 2 presents retinal alterations and these genes might be somewhat responsible for them. *RDH11* is associated with an autosomal recessive disorder (OMIM #616108), though Patient 2 presents only heterozygous alterations in the gene. *RDH12* causes Leber congenital amaurosis 13 (OMIM #612712), which can be recessive or dominant. *RDH12* variants have been linked to retinal degeneration, which Patient 2 presents. Even though they have also been reported in a few patients presenting peripapillary sparing (Garg *et al*., 2017), which is the opposite of Patient 2’s phenotype, the role of *RDH12* in retinal development might be associated with the present patient’s alterations. Both the *PIGH* (*Phosphatidylinositol Glycan Anchor Biosynthesis Class H Protein*, OMIM *600154) gene and the *ZFYVE26* (*Zinc Finger FYVE Domain-Containing Protein 26*, OMIM *612012) gene cause autosomal recessive disorders and are deleted in Patient 2, however, their remaining copy does not present variants that could link them to the patient’s phenotypes (Tremblay-Laganière *et al*., 2021). Lower expression of the chromosome 3-deleted *SYN2* (*Synapsin II*, OMIM *600755) gene have been associated to susceptibility to schizophrenia (OMIM #181500) and the patient also presents variants linked with this association in Chinese individuals; therefore, it is a phenotype that could still develop in the patient (Chen *et al*., 2004). The patient’s Dandy-Walker malformation could possibly be associated with the rupture of the *GMDS* (*GDP-mannose 4,6-dehydratase*, OMIM *602884) gene, since CNVs involving this and the *FOXC1* gene have been reported with more severe phenotypes of this common human cerebellar malformation (Aldinger *et al*., 2009). The disruption of regulatory elements of the *RUNX2* (*Runt-Related Transcription Factor 2*, OMIM *600211) due to a break in 6p21.1 may be causing its loss of function and leading to Cleidocranial dysplasia (OMIM #119600), which presents many overlaps with the patient’s phenotypes.

Patient 3 presents ten OMIM morbid genes deleted in his ∼2.3 Mb deletion in chromosome 11q23.3. Since atopic dermatitis was the main reason why his parents required medical assistance, the deleted *SIK3* (*SIK Family Kinase 3*, OMIM *614776) gene appeared as relevant given that eczema is one of the phenotypes that alterations in the gene may cause. However, the disease associated with this gene (OMIM #618162) is autosomal recessive and causes a myriad of other phenotypes that the patient does not present. The *CEP164* and *FXYD2* genes are both associated with renal conditions that the patient does not present, which is the same seen for other deleted genes that lead to phenotypes not shown by the patients. Among the genes ruptured by breakpoints, alterations in *DRD3* (*Dopamine Receptor D3*, OMIM *126541) have been associated with susceptibility to schizophrenia, and in *CBL* (*CBL protooncogene*; OMIM *165360) leads to Noonan syndrome-like disorder with or without juvenile myelomonocytic leukemia (OMIM #613563), which causes phenotypes that overlap with Patient 3, including mild delayed psychomotor development, language delay, low-set ears, and triangular face, while also causing various others that the patient lacks. Two important genes had their regulatory context disrupted due to breaks. The *DDX6* (*DEAD-box Helicase 6*, OMIM *600326) gene is associated with intellectual developmental disorder with impaired language and dysmorphic facies (OMIM #618653) with phenotypes similar to Patient 3’s. The *KMT2A* (*Lysine-Specific Methyltransferase 2A*, OMIM *159555) gene causes Wiedemann-Steiner syndrome (OMIM #605130), an autosomal dominant condition with a highly variable presentation that can lead to global developmental delay, language delay, prenatal growth deficiency, intellectual deficiency, and behavioral problems, including aggression and autistic features (Silveira *et al*., 2024), all of which Patient 3 presents; therefore, the regulatory disruption of this gene could be associated with the patients phenotypic presentation.

Patient 4 presents copy number gains in the 2q37.2q37.3 region. The *AGAP1* (*ArfGAP with GTPase Domain, Ankyrin Repeat and PH Domain 1*, OMIM *608651) gene is ruptured by two breakpoints. This gene plays a role in brain development and has been associated with neurodevelopmental disorders and autism, mainly due to loss of function (Arnold *et al*., 2016; Pacault *et al*., 2019; Lewis *et al*., 2023). Part of the gene overlaps with the patient’s first duplication and part with his triplication. Given the way that the patient’s derivative chromosome 2 is arranged, he still presents two normal copies of the gene despite being ruptured by two breakpoints, therefore we cannot entirely correlate the breaks with the patient’s autistic phenotype. Interestingly, though, several genes in the 2q37.3 region are expressed in the brain and could be associated with autism (Vera-Carbonell *et al*., 2010), such as the *GBX2* (*Gastrulation Brain Homeobox 2*, OMIM *601135) gene, which presents the highest probability of triplosensitivity (pTriplo = 1.00) among the genes in the triplication. In mice, the gene *Gbx2* plays a key role in the development of the forebrain and the hindbrain, and its gain of function led to hypoplasia of the cerebellum (Sunmonu, Chen and Li, 2009), which is a phenotype presented by our patient. Phenotypic characterization of patients with trisomy 2q is considered difficult since they present different breakpoints and affected regions, which leads to phenotypic variability (Elbracht *et al*., 2009; Ponnala *et al*., 2012). Also, these patients are usually the offspring of parents carrying balanced reciprocal translocations and therefore also present deletions, which hinders the correlation of the 2q duplication with their phenotype (Fritz *et al*., 1999; Elbracht *et al*., 2009). Our patient does not present the main characteristics found in patients with 2q duplications, such as prominent forehead, epicanthal folds, microretrognatia, abnormal ears, hypotonia, and long and slender fingers (Vera-Carbonell *et al*., 2010; Ponnala *et al*., 2012). Cardiac anomalies are also seen in 2q duplication patients (Bird and Mascarello, 2001) but not in ours. Male infants can present genital alterations such as cryptorchidism (Elbracht *et al*., 2009), which our patient also presented. It has been noted that duplications proximal to 2q33 seem to be more severe when compared to distal duplications, whose patients usually present normal stature, no major malformations, and moderate intellectual disability (Elbracht *et al*., 2009), which could be also said about patient 2. A distal 2q deletion in a patient with no associated clinical phenotype has been reported (Balikova *et al*., 2007) and has prompted the hypothesis that subtelomeric 2q duplications could be benign given that deletions are more relevant to the phenotype than duplications (Elbracht *et al*., 2009).

Patient 5 presents five CNVs in chromosome 16, three being duplications and two being deletions. Trisomy 16 is the most common autosomal trisomy recorded in miscarriages, being lethal in early pregnancy (Brisset *et al*., 2002; Lonardo *et al*., 2011). When compared to trisomy 13, 18, and 21, trisomy 16 seems to present less tolerance for higher gene dosage (Manor *et al*., 2021). While the molecular basis responsible for the developmental failure seen in spontaneous abortions is still unknown, it is hypothesized that duplications in the long arm of chromosome 16 seem to be mainly responsible for the lethality associated with trisomy 16 and consistently lead to early postnatal death when compared to 16p trisomy (Lonardo *et al*., 2011; Manor *et al*., 2021). Despite the indication that 16q trisomy has a poor general prognosis (Manor *et al*., 2021), patient 5, who presents, in total, ∼23.4 Mb of 16q duplicated regions, reached 7 years old, which seems to be same as the oldest patient described with 16q duplication, without reports of requiring constant multidisciplinary follow up, assisted respiratory support, feeding devices, intensive rehabilitation, or invasive procedures. Previously reported patients with 16q trisomy present a wide variety of congenital anomalies, with the most common being intellectual disability, speech delay, behavioral problems, generalized hypotonia, failure to thrive, short stature, central nervous system malformations, microcephaly, congenital heart defects, limb contractures, overlying fingers, clinodactyly, genitourinary anomalies, and characteristic facial dysmorphism, which includes high prominent forehead, downslanting and small palpebral fissures, epicanthal folds, a broad nasal bridge, dysplastic/low set ears, a high-arched palate, thin upper lip, and micrognathia (Brisset *et al*., 2002; Lonardo *et al*., 2011; Manor *et al*., 2021). Patient 5 does not present duplication of the terminal band 16q24, which appears to be critical for most of these phenotypes (Brisset *et al*., 2002). From the common features found in duplications involving most of 16q, patient 5 presents only a few, namely low set ears, clinodactyly, downslanted palpebral fissures, thin upper lip, overlying toes, and central nervous system malformations. Patients with duplications involving the *RPGRIP1L* (*RPGRIP1-Like*, OMIM *610937) and the *FTO* (*FTO Alpha-Ketoglutarate Dependent Dioxygenase*, OMIM *610966) genes usually present obesity (Lonardo *et al*., 2011), which is not the case for patient 5. The karyotype-phenotype correlation of duplications is difficult due to the large number of overlapped genes, reduced number of studies, and the heterogeneity of previously reported patients (Lonardo *et al*., 2011). The fact that most cases of partial 16 trisomy are associated with CNVs in other chromosomes, likely due to a parental balanced translocation, also hampers phenotype correlations. Each of the patient’s deletions overlap a small part of some of the largest human genes. The 16p deletion encompasses part of the *RBFOX1* (*RNA Binding Fox-1 Homolog 1*, OMIM *605104) gene, which is not associated with any described conditions. It has been reported that the gene is mainly expressed in muscle and the nervous system (Bhalla *et al*., 2004; Martin *et al*., 2007). Previously described patients with deletions involving the gene present intellectual and developmental delay, seizures, autism, and mild cerebellar atrophy (Bhalla *et al*., 2004; Martin *et al*., 2007). Patient 5 presents seizures and cerebellar atrophy; therefore, these phenotypes could be associated with the loss of this gene. It is important to note, though, that these patients’ deletions do not overlap with patient 5’s deletion. The 16q deletion covers part of the *WWOX* (*WW Domain Containing Oxidoreductase*, OMIM *605131) gene, which contains the second most common fragile site in the human genome (FRA16D), leading the gene to be susceptible to breaks and rearrangements (Marcelo Aldaz and Hussain, 2020). *WWOX* has a high expression in the brain and the cerebellum, being associated with Early Infantile Epileptic Encephalopathy (OMIM #616211), an autosomal recessive condition that mainly affects the nervous system. Several genes related to epilepsy were found differentially expressed in *Wwox*-KO neurospheres (Marcelo Aldaz and Hussain, 2020). Patient 5 presents phenotypes, such as cerebellar atrophy and seizures, that could be explained by the gene rupture; however, the other copy of the gene does not present any pathogenic variants and, since the gene is not haploinsufficient and heterozygous mice with a wildtype gene copy did not display any central nervous system alterations (Marcelo Aldaz and Hussain, 2020), we cannot make this correlation.

Patient 6 presents an 8.7 Mb 18q duplication, a 6.7 Mb 18q deletion, and 12 microduplications along chromosome 18 involving multiple genes. However, his clinical description was limited to mild intellectual disability, autism, and epilepsy, which hampers phenotype correlation. Among the duplicated genes, the ones with highest triplosensitivity score include *SMAD4* (*SMAD Family Member 4*, OMIM *600993), *TCF4* (*transcription factor 4*, OMIM *602272), and *ATP8B1* (*ATPase, Class I, Type 8b, Member 1*, OMIM *602397). *SMAD4* gain of function is associated with Myhre syndrome (OMIM #139210) (Lindsay *et al*., 2025), leading to multiple phenotypes that Patient 6 does not have, and there is a lack of studies investigating *TCF4* and *ATP8B1* gain of function. Among the deleted genes, three present high haploinsufficiency and lead to phenotypes not reported in Patient 6 – *CYB5A* (*Cytochrome b5, type A*, OMIM *613218) associated with male infertility, *TSHZ1* (*Teashirt Zinc Finger Homeobox 1*, OMIM *614427) linked to aural atresia, and *ZNF407* (*Zinc Finger Protein 407*, OMIM *615894) causes SIMHA syndrome (OMIM #619557) (Cody *et al*., 2015). Since the patient’s phenotypes are present in multiple disorders associated with other phenotypes, it is not feasible to pinpoint a gene or location that could be causing only them; however, it is possible that the patient may present more unreported phenotypes that could lead to a better correlation.

The karyotype-phenotype correlation for patients that present many affected chromosomes or chromosome regions is challenging, making it difficult to point out which gene defect(s) contribute to the phenotype. The smaller number of studies of the consequences of duplications when compared to deletions hampers even more the execution of a karyotype-phenotype correlation for these patients. Furthermore, different from classical microdeletion and microduplication syndromes, also known as "genomic disorders", which are observed recurrently, the CCRs studied here are likely completely unique.

### Different methodologies in the study of complex rearrangements

In this study, different methodologies were used to completely characterize the patients’ rearrangements since each of them presents its advantages and limitations. When comparing with classical cytogenomics techniques – karyotyping, CMA, and FISH – all rearrangements were better defined after OGM analysis, with lrGS completely characterizing the alterations.

Patient 2’s analysis provides a more complete view of the depth of findings by using each approach. He was studied as a newborn with the methodologies available at the time (Guilherme *et al*., 2013), which revealed nine breakpoints, eight junctions and a 14q deletion. The present study shows that, besides the deletion, he actually presents 31 breakpoints and 18 junctions, which prompted us to compare both analyses to pinpoint the information we gained by using the high-resolution techniques (Supplementary File 2 – Supplementary Figure 4). While cytogenetic analysis indicated that the four chromosomes were broken into 12 fragments (excluding the 14q deletion), high-resolution analysis found that the derivative chromosomes were formed by 23 fragments. Despite the “overall” rearrangement structure being found through FISH, classical cytogenetics techniques missed nine fragments smaller than 4 Mb, as is expected. Chromosome 3 presented the highest increase in breaks with eight breakpoints located within a 5.3 Mb region, which were missed and considered as only one break in the karyotyping and FISH analysis, likely due to seven of them being in the same chromosome band (3p25). The der(14) is the most similar derivative chromosome between methodologies while der(6) is the most different one because FISH missed the involved of small fragments from chromosomes 3 and 14. This comparison of findings illustrates the drawbacks of karyotyping and FISH with regards to resolution and of CMA in not detecting balanced alterations, which could be overcome with OGM and lrGS.

Even though, classical cytogenomics techniques’ downsides mainly concern resolution (karyotyping, CMA, and FISH), lack of balanced alterations detection (CMA), and definition of a region of investigation interest (FISH), OGM and lrGS can also present challenges.

OGM presents poor coverage on highly repetitive regions of the genome, such as centromeres, acrocentric short arms, and telomeres (Dremsek *et al*., 2021; Shim *et al*., 2024). Most importantly for this study, OGM’s resolution depends on the distance between the fluorescent labels in the DNA molecules, since, on average, it is expected to find 15 labels in every 100 kb across the genome (Mantere *et al*., 2021; Zhang *et al*., 2023). In this way, breakpoint uncertainty can be 3.2 kb on average (Mantere *et al*., 2021). Therefore, we can see that OGM does not locate the breakpoint at the nucleotide level nor does it provide a good enough distance for primer design and Sanger sequencing, which is usually limited to fragments of 1 kb (Bjørnstad *et al*., 2024; Sund *et al*., 2024).

On the other hand, lrGS allows for a better alignment to the reference genome and detection of structural variations, including the ones located in regions with repetitive elements, segmental duplications, or high GC content (Pauper *et al*., 2021; Marwaha, Knowles and Ashley, 2022). Since it does not rely on PCR amplification, it is able to analyze the DNA in its native state, which makes breakpoint detection more accurate (Mantere, Kersten and Hoischen, 2019). It still presents challenges, though, in characterizing some regions harboring SDs (van der Sanden *et al*., 2026) and the acrocentric chromosomes’ short arms, since it can be an issue to align a sequence to a single acrocentric short arm even when using longer molecules and the T2T-CHM13 reference, which maps them better (Burssed *et al*., 2026). A critical lrGS disadvantage is the high cost which, associated with variable read accuracy and throughput, hampers the use of lrGS especially in settings with fewer resources (Warburton and Sebra, 2026).

Still, every technique might have had an important role in the characterization of the patients of this study. The involvement of chromosomes 14 and 21 in Patient 1’a CCR was missed by the karyotype and CMA, with the final rearrangement only being uncovered with the OGM analysis and leading to a correction made in the karyogram with the repositioning of the der(14) and der(21) to their correct spot (Supplementary File 1 – Supplementary Figure 1). Patient 3’ CCR had an additional sixth chromosome uncovered and was partly elucidated with OGM, though with the misinterpretation of intra-fusions as CNVs and without calling some inter-fusions, requiring visual inspection. Even though his karyotype was laborious to analyze, it was vital for complete rearrangement characterization, since it showed the necessary evidence that the acrocentric chromosome short arm involved in the rearrangement was chromosome 15p. For Patient 4, karyotype was normal, CMA revealed a triplication, OGM showed that the three copies were in tandem with the middle one being inverted, given that it identified the triplication through the CNV pipeline while the SV pipeline called two inverted duplications since it only detected the fusions in the maps overlapping the extremities of the triplication (SUP), and lrGS revealed the flanking duplications, thus resolving the DUP-TRP/INV-DUP rearrangement. A return to the karyotype with this knowledge facilitated the identification of a lighter chromosome band at the 2qter corresponding to the alteration (blue arrow in Figure 5A), which had been missed in the initial analysis due to its subtlety. For Patient 5, each technique also revealed a new detail: karyotype showed the added material to chromosome 16, CMA informed the duplicated regions and its sizes, OGM elucidated part of the structure and uncovered another deletion, and lrGS complemented OGM to completely assemble the rearranged chromosome 16. Patient 6’s CGR was extremely difficult to analyze and could only be resolved with the concomitant analysis and visual inspection of OGM and lrGS, as with Patient 3, while his father’s rearranged chromosome 18 remains unresolved with only microduplication presence attested by CMA but with no information about the extra copies’ locations. In total, the complex rearrangements presented 127 breakpoints, 66 junction points and involved 14 of the 24 chromosomes (Table 1).

**Table 1.** Involved chromosomes, number (#) of breakpoints and junction points found by OGM and lrGS and average distance between these breakpoints for each patient and in total.

| P | Involved chromosomes | OGM |  | lrGS |  | Average distance between OGM BP and lrGS BP (bp) |
| --- | --- | --- | --- | --- | --- | --- |
|  |  | #BPs | #JPs | #BPs | #JPs |  |
| 1 | 12, 14, 21 | 8 | 4 | 8 | 4 | 2,641 ($\pm 1,256$) |
| 2 | 3, 6, 8, 14 | 31 | 18 | 31 | 18 | 5,989 ($\pm 6,920$) |
| 3 | 1, 3, 9, 10, 11, 15 | 30 | 15 | 42 | 28 | 6,451 ($\pm 6,325$) |
| 4 | 2 | 2 | 0 | 4 | 2 | 3,962 and 23,281<sup>a</sup> |
| 5 | 16 | 10 | 5 | 14 | 5 | 7,832 ($\pm 7,002$) |
| 6 | 18 | 16 | 9 | 28 | 9<sup>b</sup> | 5,290 ($\pm 2,317$) |
| **Total** | 16 (14 different) | 97 | 51 | 127 | 66 | 6,143 ($\pm 6,174$) |

P: patient; <sup>a</sup> Differences recorded for each of the breakpoints; <sup>b</sup> Patient 6 presented nine junctions and eight subjunctions.

For all patients in the study, the previous analyses by OGM allowed us to perform lrGS at a lower coverage (∼10× instead of the usual 30×) as the approximate locations of the patients’ rearrangements and breakpoints were already known. This targeted analysis of lrGS data based on a known chromosomal rearrangement is highly effective since lrGS detects on average 25.000 SVs per genome (Kronenberg *et al*., 2024), which sometimes cannot be properly filtered mainly due to limitations in the available population databases to determine population variation and benign alterations (Sund *et al*., 2024), especially when studying less represented groups, as is the case of this study.

To evaluate OGM’s accuracy in breakpoint detection, we compared the location of its detected breakpoints with the position of the breakpoints identified by lrGS. Considering all 127 of the patient’s breakpoints, the average distance between OGM and lrGS was 6,143 (± 6,174) bp (Table 1). Based on an inter-label distance of ∼6.6 kb, the proposed breakpoint uncertainty was supposed to be 3.2 kb (Mantere *et al*., 2021). In this study, 38% of the computed distances were below 3.2 kb, whereas 30% exceeded 6.6 kb; of these, 11 (9% of the total) were greater than 15 kb. Patient 1 presented the smallest breakpoint difference while Patient 5 showcased the largest. For patient 2, the breakpoints were not only farther away between the techniques, but also the rearrangement itself could not be entirely characterized by OGM. This happened because the labels around the breakpoint regions in the 2q37.2q37.3 region are farther apart than expected. With an average label distance of 8,745 (± 5,226) bp in the reference map and 8,546 (± 1,260) bp in the patient’s map in the upstream breakpoints’ regions, the OGM breakpoint was 3,962 bp away from the lrGS breakpoint and the 10 kb duplication could not be identified. Even more significant was the label distance in the downstream breakpoints’ region: an average label distance of 24,613 (± 8,953) bp in the reference map and 27,118 (± 13,640) bp in the patient’s map meant that the OGM and lrGS breakpoints were 23,281 bp apart and that OGM had no chance of detecting the 1 kb duplication. Grochowski et al. (2024) were able to find the DUP-TRP/INV-DUP rearrangements in at least 18 of their 24 patients using OGM. However, all 18 of them presented larger duplications than patient 2, with only one of their patients presenting a larger rearrangement size (from the beginning of the first duplication to the end of the second duplication) than him. Therefore, the individual size of each duplication seems to dictate OGM’s ability to characterize this complex rearrangement, which explains why the ones in their patients were identified and the one in ours was not. Patient 3 presented six differences above 15 kb. Upon an investigation of the OGM maps, we observed that the breakpoints fell within large unlabeled regions, with 15-30 kb without labels. Despite these differences, lrGS breakpoint location was still facilitated by their OGM locations.

OGM had been proposed as an alternative for classical cytogenomics techniques (Dremsek *et al*., 2025) and our results suggest that it could easily replace CMA and FISH, since it can provide more information than both of them combined; however, we believe it could not replace karyotyping, since the technique remains important for whole chromosome visualization. Our findings indicate that an integrated approach of karyotyping, OGM, and lower coverage lrGS is capable of completely resolving SVs and complex rearrangements. Following karyotype analysis, OGM investigation would provide a whole-genome visual structure overview of rearrangements of various types and sizes, and guide the lrGS analysis, which would precisely locate the breakpoints and characterize junction points (Dutta *et al*., 2026). Both techniques provide automatic variant calling, which Dutta et al. (2026) recommended avoiding in OGM’s case. We observed that automatic calling assists in breakpoint and SV location for both techniques by guiding the analysis directly to the regions of interest; however, the investigation must not be restricted to these calls and a visual inspection of both OGM and lrGS should be performed, especially in complex cases since they require a detailed interpretation for complete resolution. Sund et al. (2024) performed a similar experiment and hypothesized that the issue of variant callers missing information could potentially be solved with a higher lrGS coverage or a more optimized variant caller software. By fully resolving the rearrangements, the inference of a mechanism of formation and a correlation with the phenotype could be performed.

## CONCLUSION

In this study, the complex rearrangements of six patients were fully characterized, their breakpoints precisely mapped, and their junction points sequenced using a combination of long-read techniques, including OGM and lrGS. Three patients presented CCRs involving three, four, and six chromosomes while the remaining three exhibited CGRs, each involving a different chromosome. Thus, a major strength of this study is the inclusion of a diverse spectrum of complex structural variants, highlighting the complementary contributions of each technique for comprehensive rearrangement resolution. The higher resolution techniques uncovered additional complexity in all cases. Karyotype remained essential for complete rearrangement resolution even after OGM and lrGS analyses. A novel mechanism combining features of all chromoanagenesis mechanisms was proposed for two patients, evidence of an inherited alteration were identified, and the comprehensive characterization of the rearrangements allowed a correlation with the patients’ phenotypes. Overall, this study demonstrated the strengths and limitations of technologies employed, highlighting that long-reads-based approached provided the most comprehensive information owing to their higher resolution. Although no single currently available technique appears sufficient to fully resolve complex SVs, we propose that the combination of karyotyping, optical genome mapping and low-coverage long-read sequencing should be employed for rearrangement resolution, regardless of its complexity.

## ACKNOWLEDGEMENTS

We thank the patients and their families for their participation in the study. We thank the nurses of the Pediatrics Department of the São Paulo Hospital and Thainá Vilella, BSc, for their assistance in sample collection. The authors thank São Paulo Research Foundation (FAPESP) and Coordenação de Aperfeiçoamento de Pessoal de Nível Superior (CAPES) for supporting this study. We thank the Radboud Genome Technology Center for technical and support.

## LIST OF ABBREVIATIONS

ACMG: American College of Medical Genetics and Genomics
bp: base pairs
CCRs: Complex Chromosomal Rearrangements
CGRs: Complex Genomic Rearrangements
CMA: Chromosomal Microarray Analysis
CNVs: Copy Number Variants
DLE-1: Direct Label Enzyme
FISH: Fluorescence In Situ Hybridization
FoSTeS: Fork Stalling and Template Switching
gDNA: genomic DNA
GRCh38/hg38: Genome Reference Consortium Human Build 38
IGV: Integrative Genomics Viewer
JP: Junction Point
LrGS: Long-Read Genome Sequencing
MMBIR: Microhomology-Mediated Break-Induced Replication
NAHR: Non-Allelic Homologous Recombination
NHEJ: Non-Homologous End-Joining
OGM: Optical Genome Mapping
OMIM: Online Mendelian Inheritance in Man
REs: Repetitive Elements
SDs: Segmental Duplications
SNVs: Single Nucleotide Variants
SVs: Structural Variants
T2T-CHM13: Telomere-To-Telomere Complete Hydatidiform Mole 13
UHMW: Ultra-High Molecular Weight
WGS: Whole Genome Sequencing

## STATEMENTS AND DECLARATIONS

### Funding

This work was supported by São Paulo Research Foundation (FAPESP), Brazil (grants #2019/21644-0, #2022/03989-2, #2022/02202-9, and #2023/14043-5) and Coordenação de Aperfeiçoamento de Pessoal de Nível Superior (CAPES). The aims of this study contribute to the ERDERA project (to BvdS and AH), which has received funding from the European Union’s Horizon Europe research and innovation program under grant agreement No 101156595. AH was supported by the Netherlands Organization for Scientific Research (VICI grant 09150182310053).

### Ethics approval

This study’s protocol was reviewed and approved by the Ethics Committee of the Universidade Federal de São Paulo (CAAE 40846114.2.0000.5505, CEP 0028/2015; CAAE 78269424.4.0000.5505, CEP 0215/2024).

### Competing Interests

The authors have no relevant financial or non-financial interests to disclose.

### Data Availability Statement

The data that support the findings of this study are available from the corresponding author upon reasonable request.

### Consent to participate

Written informed consent was obtained from the parents.

### Consent to publish

The parents signed informed consent for publication.

### Author Contributions

**Bruna Burssed:** Conceptualization, Formal Analysis, Investigation, Data curation, Writing – Original Draft, Visualization, Project Administration. **Bart van der Sanden:** Formal Analysis, Investigation, Writing – Original Draft, Supervision. **Wolfram Höps:** Methodology, Software, Formal Analysis. **Kornelia Neveling, Eveline Kamping, and Ronald van Beek:** Investigation (OGM). **Amber den Ouden, Ronny Derks, and Raoul Timmermans:** Investigation (lrGS). **Eduardo Perrone and Marco Antonio Ramos:** Investigation (clinical evaluation of patients). **Fernanda Teixeira Bellucco:** Conceptualization, Supervision. **Alexander Hoischen:** Conceptualization, Validation, Investigation, Resources, Writing – Original Draft, Supervision, Project Administration, Funding acquisition. **Maria Isabel Melaragno:** Conceptualization, Validation, Investigation, Resources, Writing – Original Draft, Supervision, Project Administration, Funding acquisition.

## Notes

### Competing Interest Statement

The authors have declared no competing interest.

## REFERENCES

Akilapa, R.S., Smith, K. and Balasubramanian, M. (2015) ‘Clinical report: Inherited deletion of chromosome 12q21.31q21.32 associated with a distinct phenotype and intellectual disability’, Clinical Dysmorphology, 24(4), pp. 151–155. Available at: 10.1097/MCD.0000000000000096.

Aldinger, K.A. et al. (2009) ‘FOXC1 is required for normal cerebellar development and is a major contributor to chromosome 6p25.3 Dandy-Walker malformation’, Nature Genetics, 41(9), pp. 1037–1042. Available at: 10.1038/ng.422.

Arnold, M. et al. (2016) ‘The endosome localized Arf-GAP AGAP1 modulates dendritic spine morphology downstream of the neurodevelopmental disorder factor dysbindin’, Frontiers in Cellular Neuroscience, 10(SEP2016). Available at: 10.3389/fncel.2016.00218.

Aynaci, S. et al. (2025) ‘A Novel de novo Exceptional Complex Chromosomal Rearrangement Involving 5 Chromosomes Resulting in Neurodevelopmental Delay and Dysmorphism’, Molecular Syndromology, 16(6), pp. 631–639. Available at: 10.1159/000545465.

Balikova, I. et al. (2007) ‘Subtelomeric imbalances in phenotypically normal individuals’, Human Mutation, 28(10), pp. 958–967. Available at: 10.1002/humu.20537.

Beck, C.R. et al. (2019) ‘Megabase Length Hypermutation Accompanies Human Structural Variation at 17p11.2’, Cell, 176(6), pp. 1310–1324.e10. Available at: 10.1016/j.cell.2019.01.045.

Bhalla, K. et al. (2004) ‘The de novo chromosome 16 translocations of two patients with abnormal phenotypes (mental retardation and epilepsy) disrupt the A2BP1 gene’, Journal of Human Genetics, 49(6), pp. 308–311. Available at: 10.1007/s10038-004-0145-4.

Bionano (2024) ‘Bionano System Application Specifications’, Bionano Genomics, CG-00008.

Bird, L.T. and Mascarello, J.T. (2001) ‘Chromosome 2q duplications: Case report of a De Novo interstitial duplication and review of the literature’, American Journal of Medical Genetics, 100(1), pp. 13–24. Available at: 10.1002/1096-8628(20010415)100:1<13::AID-AJMG1185>3.0.CO;2-5.

Bjørnstad, P.M. et al. (2024) ‘A 39 kb structural variant causing Lynch Syndrome detected by optical genome mapping and nanopore sequencing’, European Journal of Human Genetics, 32(5), pp. 513–520. Available at: 10.1038/s41431-023-01494-7.

Bliskunova, T. et al. (2021) ‘Association of MGAT4C with major neurocognitive disorder in the Mexican population’, Gene, 778. Available at: 10.1016/j.gene.2021.145484.

Brisset, S. et al. (2002) ‘Molecular Characterization of Partial Trisomy 16q24.1-qter: Clinical Report and Review of the Literature’, American Journal of Medical Genetics, 113, pp. 339–345. Available at: 10.1002/ajmg.10740.

Burssed, B. et al. (2022) ‘Mechanisms of structural chromosomal rearrangement formation’, Molecular Cytogenetics, 15(1), pp. 1–15. Available at: 10.1186/s13039-022-00600-6.

Burssed, B. et al. (2023) ‘Fold-back mechanism originating inv-dup-del rearrangements in chromosomes 13 and 15’, Chromosome Research, 31(1), pp. 1–11. Available at: 10.1007/s10577-023-09720-0.

Burssed, B. et al. (2026) ‘Ring chromosomes uncovered by optical genome mapping: impact of telomeric-associated regions and reference genome selection on structural variant interpretation’, Chromosome Research, 34(1). Available at: 10.1007/s10577-026-09806-5.

Burwinkel, B. and Kilimann, M.W. (1998) ‘Unequal homologous recombination between LINE-1 elements as a mutational mechanism in human genetic disease’, Journal of Molecular Biology, 277(3), pp. 513–517. Available at: 10.1006/jmbi.1998.1641.

Carvalho, C.M.B. et al. (2011) ‘Inverted genomic segments and complex triplication rearrangements are mediated by inverted repeats in the human genome’, Nature Genetics, 43(11), pp. 1074–1081. Available at: 10.1038/ng.944.

Carvalho, C.M.B. et al. (2015) ‘Absence of heterozygosity due to template switching during replicative rearrangements’, American Journal of Human Genetics, 96(4), pp. 555–564. Available at: 10.1016/j.ajhg.2015.01.021.

Carvalho, C.M.B. et al. (2019) ‘Interchromosomal template-switching as a novel molecular mechanism for imprinting perturbations associated with Temple syndrome’, Genome Medicine, 11(1). Available at: 10.1186/s13073-019-0633-y.

Carvalho, C.M.B. and Lupski, J.R. (2016) ‘Mechanisms underlying structural variant formation in genomic disorders’, Nature Reviews Genetics, 17(4), pp. 224–238. Available at: 10.1038/nrg.2015.25.

Casper, J. et al. (2026) ‘The UCSC Genome Browser database: 2026 update’, Nucleic Acids Research, 54(D1), pp. D1331–D1335. Available at: 10.1093/nar/gkaf1250.

Chen, Q. et al. (2004) Family-Based Association Study of Synapsin II and Schizophrenia, Am. J. Hum. Genet.

Ciuladaite, Z. et al. (2014) ‘Relatives with opposite chromosome constitutions, rec(10)dup(10p)inv(10)(p15.1q26.12) and rec(10)dup(10q)inv(10)(p15.1q26.12), due to a familial pericentric inversion’, Cytogenetic and Genome Research, 144(2), pp. 109–113. Available at: 10.1159/000368863.

Cody, J.D. et al. (2015) ‘Consequences of chromsome 18q deletions’, *American Journal of Medical Genetics*, Part C: Seminars in Medical Genetics, 169(3), pp. 265–280. Available at: 10.1002/ajmg.c.31446.

Czakó, M. et al. (2025) ‘Uncovering Rare Structural Chromosomal Rearrangements: Insights from Molecular Cytogenetics’, International Journal of Molecular Sciences, 26(18). Available at: 10.3390/ijms26188886.

Dittwald, P. et al. (2013) ‘Inverted Low-Copy Repeats and Genome Instability-A Genome-Wide Analysis’, Human Mutation, 34(1), pp. 210–220. Available at: 10.1002/humu.22217.

Dremsek, P. et al. (2021) ‘Optical genome mapping in routine human genetic diagnostics—its advantages and limitations’, Genes, 12(12). Available at: 10.3390/genes12121958.

Dremsek, P. et al. (2025) ‘Retrospective study on the utility of optical genome mapping as a follow-up method in genetic diagnostics’, Journal of Medical Genetics, 62(2), pp. 89–96. Available at: 10.1136/jmg-2024-110265.

Duncan, A.R. et al. (2021) ‘Unique variants in CLCN3, encoding an endosomal anion/proton exchanger, underlie a spectrum of neurodevelopmental disorders’, American Journal of Human Genetics, 108(8), pp. 1450–1465. Available at: 10.1016/j.ajhg.2021.06.003.

Dutta, U.R. et al. (2026) ‘Integrative approach for delineating structural variants using optical genome mapping and long-read genome sequencing’, Molecular Biology Reports, 53(1). Available at: 10.1007/s11033-026-11906-8.

Eisfeldt, J. et al. (2023) ‘Towards routine long-read sequencing for rare disease: a national pilot study on chromosomal rearrangements’, medRxiv [Preprint]. Available at: 10.1101/2023.12.15.23299892.

Elbracht, M. et al. (2009) ‘Pure distal trisomy 2q: A rare chromosomal abnormality with recognizable phenotype’, *American Journal of Medical Genetics*, Part A, 149(11), pp. 2547–2550. Available at: 10.1002/ajmg.a.33086.

Fritz, B. et al. (1999) ‘Trisomy 2q35-q37 due to insertion of 2q material into 17q25: Clinical, cytogenetic, and molecular cytogenetic characterization’, American Journal of Medical Genetics, 87(4), pp. 297–301. Available at: 10.1002/(SICI)1096-8628(19991203)87:4<297::AID-AJMG3>3.0.CO;2-M.

Garg, A. et al. (2017) ‘Peripapillary sparing in RDH12-associated Leber congenital amaurosis’, Ophthalmic Genetics, 38(6), pp. 575–579. Available at: 10.1080/13816810.2017.1323339.

Geoffroy, V. et al. (2018) ‘AnnotSV: An integrated tool for structural variations annotation’, Bioinformatics, 34(20), pp. 3572–3574. Available at: 10.1093/bioinformatics/bty304.

Grochowski, C.M. et al. (2024) ‘Inverted triplications formed by iterative template switches generate structural variant diversity at genomic disorder loci’, Cell Genomics, 4(7). Available at: 10.1016/j.xgen.2024.100590.

Gu, W., Zhang, F. and Lupski, J.R. (2008) ‘Mechanisms for human genomic rearrangements’, PathoGenetics, 1(1), p. 4. Available at: 10.1186/1755-8417-1-4.

Guilherme, R.S. et al. (2013) ‘A complex chromosome rearrangement involving four chromosomes, nine breakpoints and a cryptic 0.6-Mb deletion in a boy with cerebellar hypoplasia and defects in skull ossification.’, Cytogenetic and genome research, 141(4), pp. 317–323. Available at: 10.1159/000353302.

Hansson, O. (2018) ‘Development of computer software to characterise and simulate molecular biology processes used in forensic DNA profiling assays’, Series of dissertations submitted to the Faculty of Medicine, University of Oslo [Preprint].

Hermetz, K.E. et al. (2014) ‘Large Inverted Duplications in the Human Genome Form via a Fold-Back Mechanism’, PLoS Genetics, 10(1). Available at: 10.1371/journal.pgen.1004139.

Kanan, Y. et al. (2008) ‘Retinol dehydrogenases RDH11 and RDH12 in the mouse retina: Expression levels during development and regulation by oxidative stress’, Investigative Ophthalmology and Visual Science, 49(3), pp. 1071–1078. Available at: 10.1167/iovs.07-1207.

Kocagil, S. et al. (2026) ‘Resolving Complex Chromosomal Rearrangements and Rare Structural Variants: An Integrated Cytogenomic Analysis of Four Cases’, Bratislava Medical Journal [Preprint]. Available at: 10.1007/s44411-026-00684-1.

Kronenberg, Z. et al. (2024) ‘The Platinum Pedigree: A long-read benchmark for genetic variants’, bioRxiv [Preprint]. Available at: 10.1101/2024.10.02.616333.

Kwon, M., Leibowitz, M.L. and Lee, J.H. (2020) ‘Small but mighty: the causes and consequences of micronucleus rupture’, Experimental and Molecular Medicine. Springer Nature, pp. 1777–1786. Available at: 10.1038/s12276-020-00529-z.

Lewis, S.A. et al. (2023) ‘AGAP1-associated endolysosomal trafficking abnormalities link gene-environment interactions in neurodevelopmental disorders’, DMM Disease Models and Mechanisms, 16(9). Available at: 10.1242/dmm.049838.

Lindsay, M.E. et al. (2025) ‘Gain-of-function variants in SMAD4 compromise respiratory epithelial function’, Journal of Allergy and Clinical Immunology, 155(1), pp. 107–119.e2. Available at: 10.1016/j.jaci.2024.08.024.

Lonardo, F. et al. (2011) ‘Clinical, cytogenetic and molecular-cytogenetic characterization of a patient with a de novo tandem proximal-intermediate duplication of 16q and review of the literature’, American Journal of Medical Genetics, Part A, 155(4), pp. 769–777. Available at: 10.1002/ajmg.a.33852.

Luo, Y. et al. (2011) ‘Diverse mutational mechanisms cause pathogenic subtelomeric rearrangements’, Human Molecular Genetics, 20(19), pp. 3769–3778. Available at: 10.1093/hmg/ddr293.

Lupski, J.R. (1998) ‘Genomic disorders: Structural features of the genome can lead to DNA rearrangements and human disease traits’, Trends in Genetics, 14(10), pp. 417–422. Available at: 10.1016/S0168-9525(98)01555-8.

Lupski, J.R. and Stankiewicz, P. (2005) ‘Genomic disorders: Molecular mechanisms for rearrangements and conveyed phenotypes’, PLoS Genetics, 1(6), pp. 0627–0633. Available at: 10.1371/journal.pgen.0010049.

Madan, K. (2012) ‘Balanced complex chromosome rearrangements: Reproductive aspects. A review’, *American Journal of Medical Genetics*, Part A, pp. 947–963. Available at: 10.1002/ajmg.a.35220.

Manor, J. et al. (2021) ‘A rare description of pure partial trisomy of 16q12.2q24.3 and review of the literature’, *American Journal of Medical Genetics*, Part A, 185(10), pp. 2903–2912. Available at: 10.1002/ajmg.a.62368.

Mantere, T. et al. (2021) ‘Optical genome mapping enables constitutional chromosomal aberration detection’, American Journal of Human Genetics, 108(8), pp. 1409–1422. Available at: 10.1016/j.ajhg.2021.05.012.

Mantere, T., Kersten, S. and Hoischen, A. (2019) ‘Long-read sequencing emerging in medical genetics’, Frontiers in Genetics, 10(MAY), pp. 1–14. Available at: 10.3389/fgene.2019.00426.

Marcelo Aldaz, C. and Hussain, T. (2020) ‘Wwox loss of function in neurodevelopmental and neurodegenerative disorders’, International Journal of Molecular Sciences, 21(23), pp. 1–23. Available at: 10.3390/ijms21238922.

Martin, C.L. et al. (2007) ‘Cytogenetic and molecular characterization of A2BP1/FOX1 as a candidate gene for autism’, American Journal of Medical Genetics, Part B: Neuropsychiatric Genetics, 144(7), pp. 869–876. Available at: 10.1002/ajmg.b.30530.

Martin, J. et al. (2004) The sequence and analysis of duplication-rich human chromosome 16. Available at: http://www.jgi.doe.gov/.

Marwaha, S., Knowles, J.W. and Ashley, E.A. (2022) ‘A guide for the diagnosis of rare and undiagnosed disease: beyond the exome’, Genome Medicine, 14(1), pp. 1–22. Available at: 10.1186/s13073-022-01026-w.

Miller, D.T. et al. (2010) ‘Consensus Statement: Chromosomal Microarray Is a First-Tier Clinical Diagnostic Test for Individuals with Developmental Disabilities or Congenital Anomalies’, American Journal of Human Genetics, 86(5), pp. 749–764. Available at: 10.1016/j.ajhg.2010.04.006.

Neveling, K. et al. (2021) ‘Next-generation cytogenetics: Comprehensive assessment of 52 hematological malignancy genomes by optical genome mapping’, American Journal of Human Genetics, 108(8), pp. 1423–1435. Available at: 10.1016/j.ajhg.2021.06.001.

Ohye, T. et al. (2014) ‘Signature of backward replication slippage at the copy number variation junction’, Journal of Human Genetics, 59(5), pp. 247–250. Available at: 10.1038/jhg.2014.20.

Pacault, M. et al. (2019) ‘A de novo 2q37.2 deletion encompassing AGAP1 and SH3BP4 in a patient with autism and intellectual disability’, European Journal of Medical Genetics, 62(12). Available at: 10.1016/j.ejmg.2018.11.020.

Pauper, M. et al. (2021) ‘Long-read trio sequencing of individuals with unsolved intellectual disability’, European Journal of Human Genetics, 29(4), pp. 637–648. Available at: 10.1038/s41431-020-00770-0.

Pellestor, F. (2019) ‘Chromoanagenesis: Cataclysms behind complex chromosomal rearrangements’, Molecular Cytogenetics, 12(1), pp. 1–12. Available at: 10.1186/s13039-019-0415-7.

Pellestor, F. and Gatinois, V. (2018) ‘Chromoanasynthesis: Another way for the formation of complex chromosomal abnormalities in human reproduction’, Human Reproduction, 33(8), pp. 1381–1387. Available at: 10.1093/humrep/dey231.

Ponnala, R. et al. (2012) ‘Phenotypic and molecular characterization of partial trisomy 2q resulting from insertion-duplication in chromosome 18q: A case report and review of literature’, Cytogenetic and Genome Research, 136(3), pp. 229–234. Available at: 10.1159/000336974.

Priya, P.K. et al. (2018) ‘Characterization of a complex chromosomal rearrangement involving chromosomes 1, 3, and 4 in a slightly affected male with bad obstetrics history’, Journal of Assisted Reproduction and Genetics, 35(4), pp. 721–725. Available at: 10.1007/s10815-018-1117-5.

Riggs, E.R. et al. (2020) ‘Technical standards for the interpretation and reporting of constitutional copy-number variants: a joint consensus recommendation of the American College of Medical Genetics and Genomics (ACMG) and the Clinical Genome Resource (ClinGen)’, Genetics in Medicine, 22(2), pp. 245– 257. Available at: 10.1038/s41436-019-0686-8.

van der Sanden, B. et al. (2026) ‘HiFi sequencing accurately identifies clinically relevant variants in paralogous genes’, The American Journal of Human Genetics, 113(6), pp. 1357–1363. Available at: 10.1016/j.ajhg.2026.04.014.

Schuy, J. et al. (2022) ‘Complex genomic rearrangements: an underestimated cause of rare diseases’, Trends in Genetics. Elsevier Ltd, pp. 1134–1146. Available at: 10.1016/j.tig.2022.06.003.

Shaffer, L.G. and Lupski, J.R. (2000) ‘C Hromosomal R Earrangements in H Umans’, Annu. Rev. Genet., 34, pp. 297–329. Available at: 10.1016/j.ajhg.2009.03.010.

Shaw, C.J. and Lupski, J.R. (2004) ‘Implications of human genome architecture for rearrangement-based disorders: The genomic basis of disease’, Human Molecular Genetics, 13(REV. ISS. 1), pp. 57–64. Available at: 10.1093/hmg/ddh073.

Shaw, C.J. and Lupski, J.R. (2005) ‘Non-recurrent 17p11.2 deletions are generated by homologous and non-homologous mechanisms’, Human Genetics, 116(1–2), pp. 1–7. Available at: 10.1007/s00439-004-1204-9.

Shim, Y. et al. (2024) ‘Comparison of Optical Genome Mapping With Conventional Diagnostic Methods for Structural Variant Detection in Hematologic Malignancies’, Annals of Laboratory Medicine, 44(4), pp. 324–334. Available at: 10.3343/alm.2023.0339.

Silveira, H.G. et al. (2024) ‘Variants in KMT2A in Three Individuals with Previous Suspicion of 22q11.2 Deletion Syndrome’, Genes, 15(2). Available at: 10.3390/genes15020211.

Stankiewicz, P., Pursley, A.N. and Cheung, S.W. (2010) ‘Challenges in clinical interpretation of microduplications detected by array CGH analysis’, *American Journal of Medical Genetics*, Part A, pp. 1089–1100. Available at: 10.1002/ajmg.a.33216.

Sund, K.L. et al. (2024) ‘Long-read sequencing and optical genome mapping identify causative gene disruptions in noncoding sequence in two patients with neurologic disease and known chromosome abnormalities’, *American Journal of Medical Genetics*, Part A [Preprint]. Available at: 10.1002/ajmg.a.63818.

Sunmonu, N.A., Chen, L. and Li, J.Y.H. (2009) ‘Misexpression of Gbx2 throughout the mesencephalon by a conditional gain-of-function transgene leads to deletion of the midbrain and cerebellum in mice’, Genesis, 47(10), pp. 667–673. Available at: 10.1002/dvg.20546.

Tremblay-Laganière, C. et al. (2021) ‘PIGH deficiency can be associated with severe neurodevelopmental and skeletal manifestations’, Clinical Genetics, 99(2), pp. 313–317. Available at: 10.1111/cge.13877.

Vera-Carbonell, A. et al. (2010) ‘Molecular characterization of a new patient with a non-recurrent inv dup del 2q and review of the mechanisms for this rearrangement’, *American Journal of Medical Genetics*, Part A, 152 A(10), pp. 2670–2680. Available at: 10.1002/ajmg.a.33613.

Vollger, M.R. et al. (2023) ‘Synchronized long-read genome, methylome, epigenome, and transcriptome for resolving a Mendelian condition.’, bioRxiv : the preprint server for biology [Preprint]. Available at: 10.1101/2023.09.26.559521.

Warburton, P.E. and Sebra, R.P. (2026) ‘Long-Read DNA Sequencing: Recent Advances and Remaining Challenges’, Annu. Rev. Genom. Hum. Genet. 2023, 24, p. 2023. Available at: 10.1146/annurev-genom-101722.

Weckselblatt, B. and Rudd, M.K. (2015) ‘Human structural variation: Mechanisms of chromosome rearrangements’, Trends in genetics : TIG, 31(10), pp. 587–599. Available at: 10.1016/j.tig.2015.05.010.

Zepeda-Mendoza, C.J. and Morton, C.C. (2019) ‘The Iceberg under Water: Unexplored Complexity of Chromoanagenesis in Congenital Disorders’, American Journal of Human Genetics, 104(4), pp. 565–577. Available at: 10.1016/j.ajhg.2019.02.024.

Zhang, S. et al. (2023) ‘Detection of cryptic balanced chromosomal rearrangements using high-resolution optical genome mapping’, Journal of Medical Genetics, 60(3), pp. 274–284. Available at: 10.1136/jmedgenet-2022-108553.

