## Supplementary material for "Integrative optical genome mapping and long-read sequencing resolve constitutional complex rearrangements at nucleotide resolution": Supplementary File 1 - Patient 1.docx

Bruna Burssed, Bart van der Sanden, Wolfram Höps, Kornelia Neveling, Eveline Kamping, Ronald van Beek, Amber den Ouden, Ronny Derks, Raoul Timmermans, Eduardo Perrone, Marco Antonio Ramos, Fernanda Teixeira Bellucco, Alexander Hoischen, Maria Isabel Melaragno

**Supplementary File 1 – Additional information in Tables and Figures from Patient 1**

**TABLES.**

**FIGURES.**

**Supplementary Table 1. Patient 1’s phenotypes with Human Phenotype Ontology (HPO) terms**

| **Phenotype** | **HPO** |
| --- | --- |
| **General** |  |
| Neurodevelopmental delay | HP:0012758 |
| Global developmental delay | HP:0001263 |
| Delayed speech and language development | HP:0000750 |
| Intellectual disability | HP:0001249 |
| Generalized hypotonia | HP:0001290 |
| **CNS** |  |
| Microcephaly | HP:0000252 |
| Plagiocephaly | HP:0001357 |
| Hypoplasia of the corpus callosum | HP:0002079 |
| **Eyes** |  |
| Strabismus | HP:0000486 |
| **Ears** |  |
| Protruding ear | HP:0000411 |
| **Thorax** |  |
| Widely spaced nipples | HP:0006610 |
| **Limbs** |  |
| Lower limb asymmetry | HP:0100559 |
| Pes cavus | HP:0001761 |
| Increased laxity of ankles | HP:0006460 |
| **Facial Dysmorphisms** |  |
| Prominent forehead | HP:0011220 |
| Bitemporal narrowness | HP:0000341 |
| Synophrys | HP:0000664 |
| Depressed nasal bridge | HP:0005280 |
| Smooth philtrum | HP:0000319 |
| Thin upper lip vermilion | HP:0000219 |
| Thin lower lip vermilion | HP:0010282 |
| Downturned corners of mouth | HP:0002714 |
| Widely spaced teeth | HP:0000687 |

**Supplementary Table 2. Results of the techniques performed for the patient and final rearrangement.**

| **Technique** | **Result** |
| --- | --- |
| **Karyotype** | 46,XX,del(12)(q?22qq?24.1) |
| **Chromosomal Microarray Analysis** | arr[GRCh38] 4q33q34.1(169400580_171774496)×1 |
| **Optical Genome Mapping** | ogm[GRCh38] 4q33q34.1(169388085_171786819)×1 |
| **Long-read Sequencing** | seq[GRCh38] 4q33q34.1(169440001_171792000)×1 |
| **Final Rearrangement** | 46,XX,t(12;14;21)  (12pter→12q21.32(chr12:86,717,790)::12q23.3(chr12:105,474,617)→12qter;  14pter→14q12(chr14:29,367,001):: 21q21.1(chr21:22,080,569)→21qter;  21pter→21q21.1(chr21:22,080,571)::12q23.3(chr12:105,474,616)→12q21.32(chr12:86,717,792)::14q12(chr14:29,367,002)→14qter)dn |
|  | 46,XX,t(12;14;21)(12pter→12q21.32::12q23.3→12qter;14pter→14q12:: 21q21.1→21qter;21pter→21q21.1::12q23.3→12q21.32::14q12→14qter)dn |

**Supplementary Table 3. Breakpoint (BP) coordinates found through long-read genome sequencing and their number (#) for identification.**

| **#BP** | **BP coordinate** |
| --- | --- |
| 1 | chr12:86,717,790 |
| 2 | chr12:86,717,792 |
| 3 | chr12:105,474,616 |
| 4 | chr12:105,474,617 |
| 5 | chr14:29,367,001 |
| 6 | chr14:29,367,002 |
| 7 | chr21:22,080,569 |
| 8 | chr21:22,080,571 |

**Supplementary Table 4. Junction points and breakpoints details found on the long-read genome sequencing analysis and annotation of Repetitive Elements in regions surrounding the breakpoints.**

| **Junction Point** | | | | | **Breakpoint** | | **Repetitive Elements** | | | **Obs.** | |
| --- | --- | --- | --- | --- | --- | --- | --- | --- | --- | --- | --- |
| **der** | **#JP** | **JP ID** | **Ins** | **MH** | **#BP** | **BP coord.** | **BP - 1 kb** | **BP** | **BP + 1 kb** | |  |
| **12** | 1 | fus(12) | 88  (Repeating sequence: TATATATGTATG) | - | 1 | chr12:86,717,790 | SINE (tRNA-RTE) – MamSINE1  chr12:86716891-86716986 (-) | - | DNA (hAT-Charlie) – MER20  chr12:86717971-86718180 (+) | | 1bp del: chr12:86,717,791 |
|  |  |  |  |  |  |  |  |  | LTR (ERVL-MaLR) – MLT1G  chr12:86718279-86718577 (-) | |  |
|  |  |  |  |  |  |  |  |  | DNA (hAT-Charlie) – MER1B  chr12:86718578-86718920 (+) | |  |
|  |  |  |  |  | 4 | chr12:105,474,617 | LTR (Gypsy) – MamGypsy2-LTR  chr12:105473693-105473796 (-) | LINE (L2) – L2a  chr12:105474305-105474977 (-) | SINE (Alu) – AluJr  chr12:105475111-105475383 (+) | |  |
|  |  |  |  |  |  |  |  |  | DNA (hAT-Tip100) – Zaphod  chr12:105475392-105475485 (-) | |  |
| **14** | 2 | t(14;21) | - | - | 5 | chr14:29,367,001 | SINE (Alu) – AluJr  chr14:29366159-29366458 (+) | LINE (L1) – L1ME3A  chr14:29366459-29367412 (+) | LINE (L1) – L1ME3A  chr14:29367408-29369444 (+) | | 3bp dup: chr21:22,080,569-22,080,571 |
|  |  |  |  |  |  |  | LINE (L1) – L1ME3A  chr14:29366029-29366158 (+) |  |  |  |  |
|  |  |  |  |  | 7 | chr21:22,080,569 | LTR (ERVL-MaLR) – MLT1G  chr21:22080083-22080307 (+) | - | SINE (MIR) – MIR3  chr21:22081050-22081127 (+) | |  |
|  |  |  |  |  |  |  | SINE (Alu) – AluSc  chr21:22079372-22079676 (+) |  | LTR (ERV1) – LTR8A  chr21:22081359-22082080 (-) | |  |
| **21** | 3 | t(12;21) | - | - | 8 | chr21:22,080,571 | LTR (ERVL-MaLR) – MLT1G  chr21:22080083-22080307 (+) | - | SINE (MIR) – MIR3  chr21:22081050-22081127 (+) | |  |
|  |  |  |  |  |  |  | SINE (Alu) – AluSc  chr21:22079372-22079676 (+) |  | LTR (ERV1) – LTR8A  chr21:22081359-22082080 (-) | |  |
|  |  |  |  |  | 3 | chr12:105,474,616 | LTR (Gypsy) – MamGypsy2-LTR  chr12:105473693-105473796 (-) | LINE (L2) – L2a  chr12:105474305-105474977 (-) | SINE (Alu) – AluJr  chr12:105475111-105475383 (+) | |  |
|  |  |  |  |  |  |  |  |  | DNA (hAT-Tip100) – Zaphod  chr12:105475392-105475485 (-) | |  |
| **21** | 4 | t(12;14) | - | - | 2 | chr12:86,717,792 | SINE (tRNA-RTE) – MamSINE1  chr12:86716891-86716986 (-) | - | DNA (hAT-Charlie) – MER20  chr12:86717971-86718180 (+) | |  |
|  |  |  |  |  |  |  |  |  | LTR (ERVL-MaLR) – MLT1G  chr12:86718279-86718577 (-) | |  |
|  |  |  |  |  |  |  |  |  | DNA (hAT-Charlie) – MER1B  chr12:86718578-86718920 (+) | |  |
|  |  |  |  |  | 6 | chr14:29,367,002 | SINE (Alu) – AluJr  chr14:29366159-29366458 (+) | LINE (L1) – L1ME3A  chr14:29366459-29367412 (+) | LINE (L1) – L1ME3A  chr14:29367408-29369444 (+) | |  |
|  |  |  |  |  |  |  | LINE (L1) – L1ME3A  chr14:29366029-29366158 (+) |  |  |  |  |
| **der** | **#JP** | **JP ID** | **Ins** | **MH** | **#BP** | **BP coord.** | **BP - 1 kb** | **BP** | **BP + 1 kb** | | **Obs.** |
| **Junction Point** | | | | | **Breakpoint** | | **Repetitive Elements** | | |  | |

der: derivative chromosome where the junction is located; JP: junction point; JP ID: junction point identification; Ins: insertion; SJP: sub-junction point; MH: microhomology; Del: deletion; BP: breakpoint; coord.: coordinate; SNVs: single nucleotide variants; Obs.: observation. All genomic coordinates are according to reference genome GRCh38/hg38.

**Supplementary Table 5. Difference between breakpoints found by Optical Genome Mapping (OGM) and Long-read Genome Sequencing (lrGS).**

| **#JP** | **JP ID** | **BP** | **BP OGM** | | **BP lrGS** | **BPs difference (bp)** | **Sum of BPs difference (bp)** | **Size of Uncertain OGM region (bp)** |
| --- | --- | --- | --- | --- | --- | --- | --- | --- |
| 1 | fus(12) | 1 | chr12:86,714,964 | | chr12:86,717,790 | 2,826 | 5,834 | 0 |
|  |  | 4 | chr12:105,471,609 | | chr12:105,474,617 | 3,008 |  |  |
| 2 | t(14;21) | 5 | chr14:29,365,337 | | chr14:29,367,001 | 1,664 | 1,876 | 1.895 |
|  |  | 7 | chr21:22,080,781 | | chr21:22,080,569 | 212 |  |  |
| 3 | t(12;21) | 8 | chr21:22,076,521 | | chr21:22,080,571 | 4,050 | 7,057 | 7.131 |
|  |  | 3 | chr12:105,471,609 | | chr12:105,474,616 | 3,007 |  |  |
| 4 | t(12;14) | 2 | chr12:86,720,162 | | chr12:86,717,792 | 2,370 | 6,361 | 6.997 |
|  |  | 6 | chr14:29,370,993 | | chr14:29,367,002 | 3,991 |  |  |
|  | | | | **Average** | | 2,641 |  |  |
|  | | | | **Standard Deviation** | | 1,256 |  |  |

JP: junction point; JP ID: junction point identification; BP(s): breakpoint(s); NF: not found. All genomic coordinates are according to reference genome GRCh38/hg38.

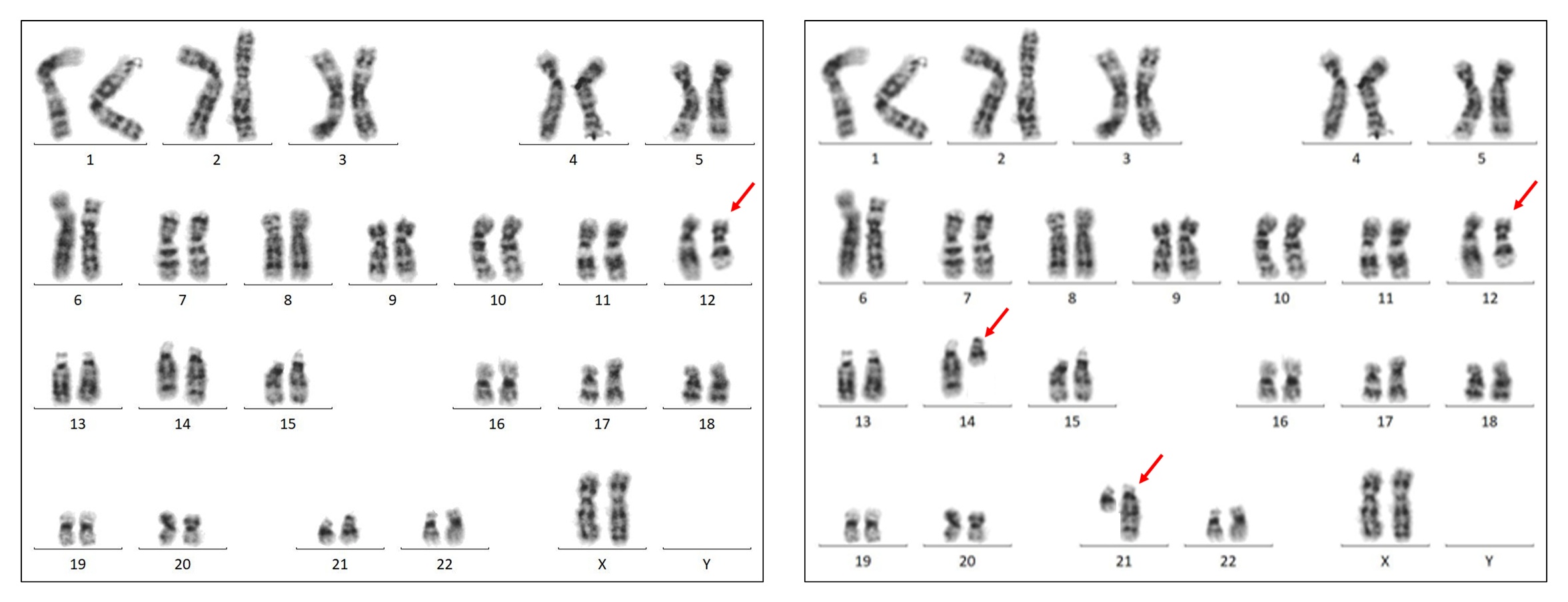

**Supplementary Figure 1. Karyotype of Patient 1.**

On the left, the initial assembled karyotype showing an altered chromosome 12 (red arrow). On the right, the revised version of the karyotype with the repositioning of the derivative chromosomes after the complete characterization of the rearrangement.

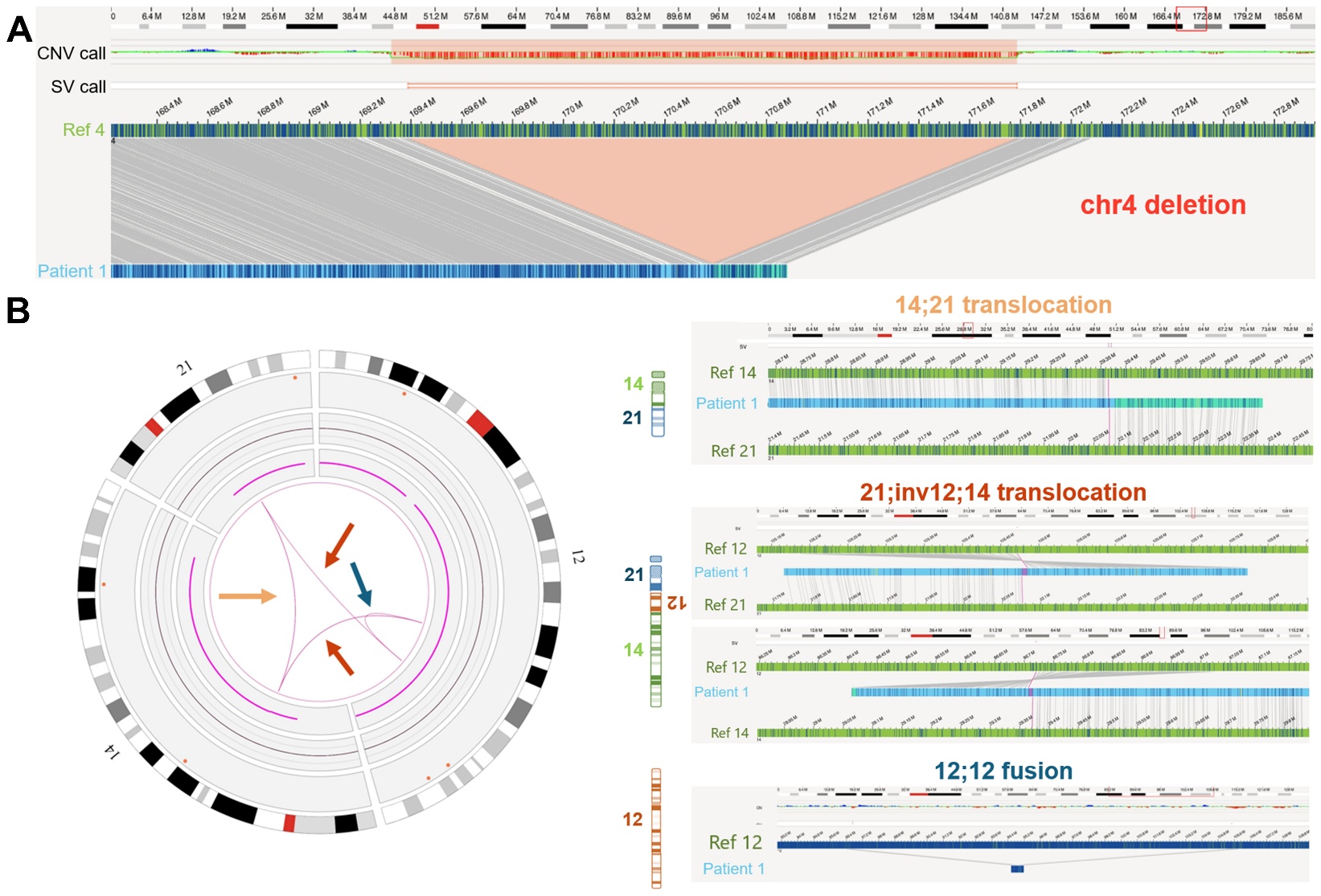

**Supplementary Figure 2. OGM results for Patient 1.**

**(A) Chromosome 4 deletion.** From top to bottom, chromosome 4 idiogram, CNV call track, SV call track, reference map from chromosome 4 (hg38) in green with labels in blue, patient map in light blue with labels in dark blue. The CNV call track shows a ~2.4 Mb deletion (red square), which is also called in the SV call (small red horizontal lines). Below, the region in red shows the labels present in the reference map that are missing in the patient map.

**(B) CCR.** To the left, circos plot showing chromosomes 12, 14, and 21. The chromosomes’ idiograms can be seen in the periphery while the pink lines in the middle represent the translocations. To the right, OGM maps from each translocation displaying chromosome idiogram, reference map (hg38) in green with labels in blue, patient map in light blue with labels in dark blue. The orange arrow highlights the translocation between chromosomes 14 and 21 with the patient map (at the top of the right panel) aligning to the reference of chromosome 14 then to the reference of chromosome 21. The red arrows arrow highlights the translocations between chromosomes 12 and 21 and between chromosomes 12 and 14 with the patient maps (in the middle of the right panel) aligning to the reference of chromosome 12 and to the reference of chromosomes 21 and 14. The blue arrow highlights the chromosome 12 fusion with the patient map (at the bottom of the right panel) aligning to different regions of the reference of chromosome 12.

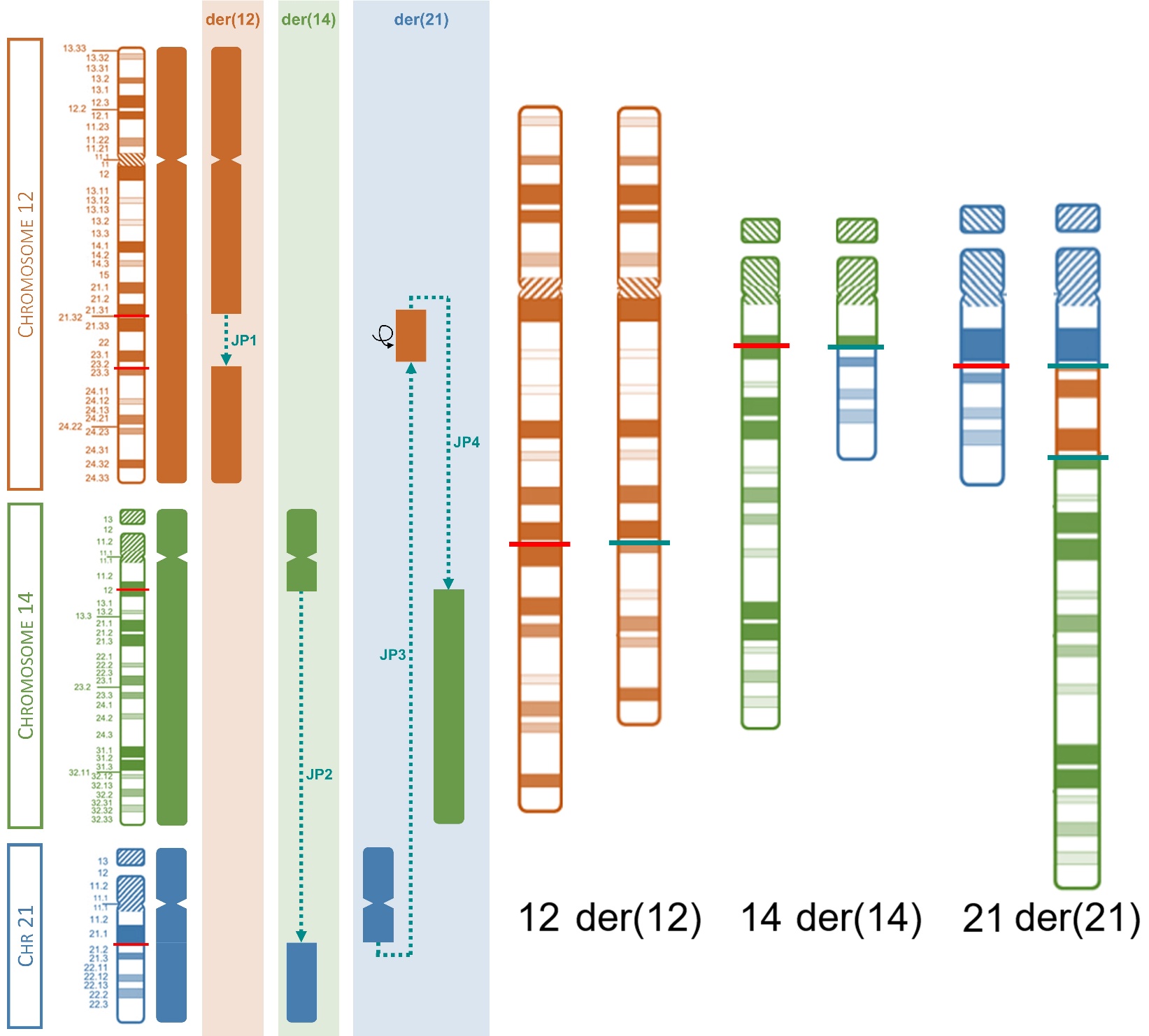

**Supplementary Figure 3. Patient 1’s resolved complex chromosomal rearrangement**

To the left, a schematic representation of the junctions between all chromosome regions involved in the CCR. To the right, idiogram of the chromosomes involved in the complex chromosomal rearrangement. Red lines show the breakpoints in the normal chromosomes and cyan lines show the junction points in the derivative chromosomes.
