## Supplementary material for "Integrative optical genome mapping and long-read sequencing resolve constitutional complex rearrangements at nucleotide resolution": Supplementary File 2 - Patient 2.docx

Bruna Burssed, Bart van der Sanden, Wolfram Höps, Kornelia Neveling, Eveline Kamping, Ronald van Beek, Amber den Ouden, Ronny Derks, Raoul Timmermans, Eduardo Perrone, Marco Antonio Ramos, Fernanda Teixeira Bellucco, Alexander Hoischen, Maria Isabel Melaragno

**Supplementary File 2 – Additional information in Tables and Figures from Patient 2**

**TABLES.**

**FIGURES.**

**Supplementary Table 1. Patient 2’s phenotypes with Human Phenotype Ontology (HPO) terms**

| **Phenotype** | **HPO** |
| --- | --- |
| **General** |  |
| Neurodevelopmental delay | HP:0012758 |
| Speech delay | HP:0000750 |
| Failure to thrive | HP:0001508 |
| **Brain** |  |
| Dandy-Walker malformation | HP:0001305 |
| Hypoplasia of the corpus callosum | HP:0002079 |
| Enlargement of the cisterna magna | HP:0002280 |
| Cerebellar vermis hypoplasia | HP:0001320 |
| **Bones** |  |
| Bell-shaped thorax | HP:0001591 |
| Hypoplastic right clavicle | HP:0000894 |
| Marked reduction in ossification of the cranial vault (Decreased calvarial ossification) | HP:0005474 |
| Large fontanelles | HP:0000239 |
| Wormian bones | HP:0002645 |
| Sclerosis of skull base | HP:0002694 |
| **Cardiovascular** |  |
| Abnormal vena cava physiology | HP:0030970 |
| Persistent left superior vena cava | HP:0005301 |
| **Renal** |  |
| Pieloectasia renal bilateral (Dilatation of the renal pelvis) | HP:0010946 |
| **Eyes** |  |
| Peripapillary atrophy | HP:0500087 |
| Diffuse thinning of the retinal pigment epithelium (Retinal thinning) | HP:0030329 |
| **Ear** |  |
| Bilateral conductive hearing loss | HP:0008513 |
| External auditory canal stenosis | HP:0000402 |
| **Facial dysmorphisms** |  |
| High forehead | HP:0000348 |
| Prominent forehead | HP:0011220 |
| Large fontanelles | HP:0000239 |
| Protruding eyes (Proptosis) | HP:0000520 |
| Hypertelorism | HP:0000316 |
| Epicanthal folds | HP:0000286 |
| Low-set ears | HP:0000369 |
| Exaggerated cupid's bow | HP:0002263 |
| Downturned corners of mouth | HP:0002714 |
| Short neck | HP:0000470 |
| Wide neck | HP:0000475 |
| Cervical hypotonia (Neck muscle hypoplasia) | HP:0008984 |
| Swallowing difficulty (dysphagia) | HP:0002015 |

**Supplementary Table 2. Results of the techniques performed for the patient and final rearrangement.**

| **Technique** | **Result** |
| --- | --- |
| **Karyotype** | 46,XY,add(3)(p26),?inv(6)(p21q15),del(8)(q24),del(14)(q24) |
| **Chromosomal Microarray Analysis** | arr[GRCh38] 14q24.1(67,474,797_68,060,395)×1 |
| **Optical Genome Mapping** | ogm[GRCh38] 14q24.1(67,458,617_68,215,643)×1 |
| **Long-read Sequencing** | seq[GRCh38] 3p25.2(12,157,018_12,158,900)×1, 14q24.1(67,473,334_68,065,355)×1 |
| **Final Rearrangement**  (colors represent similar breakpoints) | **der(3):** 14qter→14q24.1(chr14:68,065,355)::6p22.3(chr6:22,216,040)→6p21.1(chr6:45,041,636)::6p12.3(chr6:46,293,571)→  6p21.1(chr6:45,617,626)::3p25.2(chr3:12,158,900)→3p25.1(chr3:16,137,682)::3p25.3(chr3:11,062,692)→3p25.3(chr3:11,536,045)::  3p24.3(chr3:16,433,321)**→**3q26.1zchr3:162,794,346)::3q26.1(chr3:162,908,547)→3qter  **der(6):** 8qter→8q23.3(chr8:111,765,700)::6p12.3(chr6:46,293,571)**→**6q13(chr6:74,206,502)::3p25.2(chr3:12,157,018)→  3p25.3(chr3:11,536,053)::6q13(chr6:74,206,522)→6q16.1(chr6:96,991,559)::14q23.3(chr14:65,594,522)→14q24.1(chr14:67,473,334)::  6q16.1(chr6:96,991,559)→6q16.1(chr6:98,626,637)::6q24.1(chr6:12,214,527)→6p25.3(chr6:2,029,107)::6p22.3(chr6:21,627,912)→  6p22.3(chr6:22,216,040)::6q16.1(chr6:98,626,637)→6qter  **der(8):** 8pter**→**8q23.3(chr8:111,765,702)::6p21.1(chr6:45,617,625)→6p21.1(chr6:45,041,636)::3p24.3(chr3:16,433,321)→  3p25.1(chr3:16,137,682)::3p25.3(chr3:11,062,692)→3pter  **der(14):** 14pter**→**14q23.3(chr14:65,594,523)::6q24.1(chr6:12,214,527)→6p22.3(chr6:21,627,913):: 6p25.3(chr6:2,029,106)→6pter |
|  | 46,XY,t(3;6;8;14)(14qter→14q24.1::6p22.3→6p21.1::6p12.3→6p21.1::3p25.2→3p25.1::3p25.3→3p25.3::3p24.3→3qter;8qter→8q23.3::  6p12.3→6q13::3p25.2→3p25.3::6q13→6q16.1::14q23.3→14q24.1::6q16.1→6q16.1::6q24.1→6p25.3::6p22.3→6p22.3::6q16.1→6qter;  8pter→8q23.3::6p21.1→6p21.1::3p24.3→3p25.1::3p25.3→3pter;14pter→14q23.3::6q24.1→6p22.3::6p25.3→6pter) |

**Supplementary Table 3. Breakpoint coordinates found through long-read sequencing and their number (#) for identification.**

| **#BP** | **BP coordinate** |
| --- | --- |
| **1** | chr3:11,062,692 |
| **2** | chr3:11,536,045 |
| **3** | chr3:11,536,053 |
| **4** | chr3:12,157,018 |
| **5** | chr3:12,158,900 |
| **6** | chr3:16,137,682 |
| **7** | chr3:16,433,321 |
| **8** | chr3:16,433,327 |
| **9** | chr6:2,029,106 |
| **10** | chr6:2,029,107 |
| **11** | chr6:12,214,526 |
| **12** | chr6:12,214,527 |
| **13** | chr6:21,627,912 |
| **14** | chr6:21,627,913 |
| **15** | chr6:22,216,039 |
| **16** | chr6:22,216,040 |
| **17** | chr6:45,041,636 |
| **18** | chr6:45,617,625 |
| **19** | chr6:45,617,626 |
| **20** | chr6:46,293,565 |
| **21** | chr6:46,293,571 |
| **22** | chr6:74,206,502 |
| **23** | chr6:74,206,520 |
| **24** | chr6:96,991,559 |
| **25** | chr6:98,626,637 |
| **26** | chr8:111,765,700 |
| **27** | chr8:111,765,702 |
| **28** | chr14:65,594,522 |
| **29** | chr14:65,594,523 |
| **30** | chr14:67,473,334 |
| **31** | chr14:68,065,355 |

**Supplementary Table 4. Junction points and breakpoints details found on the long-read sequencing analysis and annotation of Repetitive Elements in regions surrounding the breakpoints.**

| **Junction Point** | | | | | | **Breakpoint** | | **Repetitive Elements** | | | **Obs.** |
| --- | --- | --- | --- | --- | --- | --- | --- | --- | --- | --- | --- |
| **der** | **#JP** | **JP ID** | **Ins** | **MH** | **#BP** | | **BP coord.** | **BP - 1 kb** | **BP** | **BP + 1 kb** |  |
| 3 | **1** | t(6;14).1 | - | 1 (G) | **31** | | chr14:68,065,355 | LINE (L1) – L1M2  chr14:68063776-68065287 (-) | - | LINE (L1) – L1ME4a  chr14:68065860-68066235 (+) | - |
|  |  |  |  |  |  |  |  |  |  | Simple Rep – (TTCTGTT)n  chr14:68066331-68066364 (+) |  |
|  |  |  |  |  | **16** | | chr6:22,216,040 | DNA (TcMar-Tigger) – Tigger3b  chr6:22214822-22216016 (+) | - | LTR (ERVL-MaLR) – MLT1H  chr6:22216118-22216256 (-) |  |
|  |  |  |  |  |  |  |  |  |  | SINE (MIR) – MIR  chr6:22216569-22216725 (-) |  |
|  |  |  |  |  |  |  |  |  |  | LINE (CR1) – L3  chr6:22216793-22217126 (-) |  |
| 3 | **2** | fus(6).1 | 1 (A) | - | **17** | | chr6:45,041,636 | LINE (L2) – L2a  chr6:45041041-45041539 (-) | - | SINE (MIR) – MIRb  chr6:45041792-45041949 (+) | 1 bp dup: chr6:45,041,636 |
|  |  |  |  |  |  |  |  |  |  | SINE (Alu) – AluSq2  chr6:45042141-45042432 (+) |  |
|  |  |  |  |  |  |  |  |  |  | LINE (L1) – L1P2  chr6:45042540-45042557 (+) |  |
|  |  |  |  |  |  |  |  |  |  | LINE (L1) – L1PA4  chr6:45042558-45042608 (-) |  |
|  |  |  |  |  | **20** | | chr6:46,293,565 | LINE (L1) – L1PA7  chr6:46292725-46293245 (-) | - | LINE (L2) – L2c  chr6:46293627-46293974 (-) | 5 bp del: chr6:46,293,566-46,293,570 |
|  |  |  |  |  |  |  |  |  |  | LINE (L1) – L1ME4a  chr6:46294138-46294720 (+) |  |
| 3 | **3** | t(3;6).1 | - | - | **19** | | chr6:45,617,626 | LINE (CR1) – CR1-3_Croc  chr6:45617484-45617535 (+) | - | LINE (CR1) – L3b  chr6:45617689-45617777 (+) | - |
|  |  |  |  |  |  |  |  | SINE (MIR) – MIR  chr6:45617358-45617427 (-) |  |  |  |
|  |  |  |  |  |  |  |  | LINE (L2) – L2a  chr6:45617179-45617327 (-) |  |  |  |
|  |  |  |  |  |  |  |  | LINE (CR1) – L3  chr6:45616884-45617098 (-) |  |  |  |
|  |  |  |  |  | **5** | | chr3:12,158,900 | LINE (L2) – L2b  chr3:12157988-12158278 (-) | - | Simple Rep – (ACAG)n  chr3:12159252-12159328 (+) |  |
|  |  |  |  |  |  |  |  | Simple Rep – (GCA)n  chr3:12158774-12158806 (+) |  | SINE (Alu) – AluSx  chr3:12159761-12160072 (+) |  |
| **Junction Point** | | | | | **Breakpoint** | | | **Repetitive Elements** | | | **Obs.** |
| **der** | **#JP** | **JP ID** | **Ins** | **MH** | **#BP** | | **BP coord.** | **BP – 1 kb** | **BP** | **BP + 1 kb** |  |
| 3 | **4** | fus(3).1 | - | 1 (G) | **6** | | chr3:16,137,682 | DNA (hAT-Blackjack) – MER81  chr3:16137577-16137675 (-) | - | LTR (ERVL-MaLR) – MLT1I  chr3:11063055-11063352 (-) | - |
|  |  |  |  |  |  |  |  | ERVL-MaLR (LTR) – MLT1A  chr3:16137364-16137576 (+) |  |  |  |
|  |  |  |  |  |  |  |  | Simple Rep – (TC)n  chr3:16137338-16137363 (+) |  |  |  |
|  |  |  |  |  |  |  |  | DNA (hAT-Blackjack) – MLT1A  chr3:16137172-16137337 (+) |  | SINE (Alu) – AluY  chr3:11063424-11063728 (-) |  |
|  |  |  |  |  |  |  |  | DNA (hAT-Blackjack) – MER81  chr3:16137159-16137171 (-) |  |  |  |
|  |  |  |  |  | **1** | | chr3:11,062,692 | LTR (ERV1) – LTR12C  chr3:11061209-11062580 (+) | LINE (L2) – L2a  chr3:11062583-11063054 (-) | LTR (ERVL-MaLR) – MLT1I  chr3:11063055-11063352 (-) |  |
|  |  |  |  |  |  |  |  |  |  | SINE (Alu) – AluY  chr3:11063424-11063728 (-) |  |
| 3 | **5** | fus(3).2 | - | 1 (A) | **2** | | chr3:11,536,045 | LINE (L2) – L2c  chr3:11535178-11535381 (+) | - | SINE (MIR) – MIR3  chr3:11536719-11536895 (+) | 7 bp del: chr3:11,536,046-11,536,052 |
|  |  |  |  |  |  |  |  |  |  | LINE (L2) – L2b  chr3:11536912-11537012 (+) |  |
|  |  |  |  |  | **8** | | chr3:16,433,327 | SINE (Alu) – AluJr  chr3:16432556-16432684 (+) | - | SINE (Alu) – AluJb  chr3:16434242-16434361 (+) | - |
|  |  |  |  |  |  |  |  | SINE (Alu) – AluJb  chr3:16432236-16432545 (+) |  |  |  |
| 6 | **6** | t(6;8).1 | - | - | **26** | | chr8:111,765,700 | LINE (L1) – L1MC3  chr8:111764958-111765641 (+) | LTR (ERV1) – LTR1A1  chr8:111765642-111766427 (+) | LINE (L1) – L1MC3  chr8:111766428-111766465 (+) | 3 bp dup: chr8:111,765,700-chr8:111,765,702 |
|  |  |  |  |  |  |  |  | LINE (L1) – L1PA7  chr8:111764568-111764957 (+) |  |  |  |
|  |  |  |  |  | **21** | | chr6:46,293,571 | LINE (L1) – L1PA7  chr6:46292725-46293245 (-) | - | LINE (L2) – L2c  chr6:46293627-46293974 (-) | - |
|  |  |  |  |  |  |  |  |  |  | LINE (L1) – L1ME4a  chr6:46294138-46294720 (+) |  |
| 6 | **7** | t(3;6).2 | 3 (TCT) | 9 (GAGAGGTTTT) | **22** | | chr6:74,206,502 | - | LINE (L1) – L1MA5  chr6:74204925-74207294 (-) | Simple Rep – (TA)n  chr6:74207295-74207324 (+) | 7 bp del: chr6:74,206,513-74,206,519 |
|  |  |  |  |  |  |  |  |  |  | LINE (L1) – L1MA5  chr6:74207325-74208081 (-) |  |
|  |  |  |  |  | **4** | | chr3:12,157,018 | LINE (CR1) – L3  chr3:12156450-12156677 (+) | - | Simple Rep – (AAAAC)n  chr3:12157093-12157119 (+) | - |
|  |  |  |  |  |  |  |  |  |  | LINE (L2) – L2b  chr3:12157988-12158278 (-) |  |
| **Junction Point** | | | | | **Breakpoint** | | | **Repetitive Elements** | | | **Obs.** |
| **der** | **#JP** | **JP ID** | **Ins** | **MH** | **#BP** | | **BP coord.** | **BP – 1 kb** | **BP** | **BP + 1 kb** |  |
| 6 | **8** | t(3;6).3 | - | 2 (GG) | **3** | | chr3:11,536,053 | LINE (L2) – L2c  chr3:11535178-11535381 (+) | - | SINE (MIR) – MIR3  chr3:11536719-11536895 (+) | - |
|  |  |  |  |  |  |  |  |  |  | LINE (L2) – L2b  chr3:11536912-11537012 (+) |  |
|  |  |  |  |  | **23** | | chr6:74,206,520 | - | LINE (L1) – L1MA5  chr6:74204925-74207294 (-) | Simple Rep – (TA)n  chr6:74207295-74207324 (+) |  |
|  |  |  |  |  |  |  |  |  |  | LINE (L1) – L1MA5  chr6:74207325-74208081 (-) |  |
| 6 | **9** | t(6;14).2 | - | - | **24** | | chr6:96,991,559 | - | LINE (L1) – L1PA15  chr6:96989111-96992512 (-) | - | 1 bp dup: chr6:96,991,559 |
|  |  |  |  |  | **28** | | chr14:65,594,522 | SINE (Alu) – AluSx  chr14:65593878-65594009 (-) | LTR (ERV1) – LTR28B  chr14:65594101-65595106 (-) | - | - |
|  |  |  |  |  |  |  |  | SINE (Alu) – AluSp  chr14:65593565-65593874 (-) |  |  |  |
| 6 | **10** | t(6;14).3 | - | - | **30** | | chr14:67,473,334 | - | - | SINE (Alu) – AluSz  chr14:67473971-67474275 (+) | - |
|  |  |  |  |  | **24** | | chr6:96,991,559 | - | LINE (L1) – L1PA15  chr6:96989111-96992512 (-) | - |  |
| 6 | **11** | fus(6).2 | - | 3 (ACC) | **25** | | chr6:98,626,637 | Simple Rep – (TG)n  chr6:98626163-98626211 (+) | - | tRNA (tRNA) – tRNA-Ile-ATA  chr6:98626784-98626822 (-) | 1 bp dup: chr6:98,626,637 |
|  |  |  |  |  |  |  |  | DNA (hAT-Charlie) – MER33  chr6:98625857-98626117 (-) |  | Simple Rep – (AC)n  chr6:98627045-98627075 (+) |  |
|  |  |  |  |  |  |  |  |  |  | LINE (RTE-BovB) – MamRTE1  chr6:98627447-98627566 (-) |  |
|  |  |  |  |  | **12** | | chr6:12,214,527 | SINE (MIR) – MIR3  chr6:12214200-12214263 (-) | - | LTR (ERVL-MaLR) – THE1B  chr6:12214567-12214922 (-) | - |
|  |  |  |  |  |  |  |  | SINE (Alu) – AluJr  chr6:12213563-12213849 (+) |  | SINE (Alu) – AluJb  chr6:12215009-12215308 (-) |  |
|  |  |  |  |  |  |  |  | LINE (L1) – HAL1b  chr6:12213285-12213538 (-) |  | LINE (Penelope) – Penelope1_Vert chr6:12215317-12215393 (+) |  |
| 6 | **12** | fus(6).3 | - | - | **10** | | chr6:2,029,107 | Simple Rep – (T)n  chr6:2028983-2029009 (+) | - | DNA (hAT-Tip100) – MamTip1  chr6:2029869-2029957 (+) | - |
|  |  |  |  |  | **13** | | chr6:21,627,912 | Simple Rep – (AAAT)n  chr6:21627895-21627908 (+) | SINE (Alu) – AluSq  chr6:21627909-21628165 (+) | Simple Rep – (AAAT)n  chr6:21628166-21628177 (+) | 2 bp dup: chr6:21,627,912-21,627,913 |
|  |  |  |  |  |  |  |  | SINE (Alu) – AluY  chr6:21627286-21627593 (+) |  |  |  |
|  |  |  |  |  |  |  |  | Simple Rep – (ATTTA)n  chr6:21627168-21627207 (+) |  |  |  |
| **Junction Point** | | | | | **Breakpoint** | | | **Repetitive Elements** | | | **Obs.** |
| **der** | **#JP** | **JP ID** | **Ins** | **MH** | **#BP** | | **BP coord.** | **BP – 1 kb** | **BP** | **BP + 1 kb** |  |
| 6 | **13** | fus(6).4 | - | 3 (CCT) | **15** | | chr6:22,216,039 | DNA (TcMar-Tigger) – Tigger3b  chr6:22214822-22216016 (+) | - | LTR (ERVL-MaLR) – MLT1H  chr6:22216118-22216256 (-) | - |
|  |  |  |  |  |  |  |  |  |  | SINE (MIR) – MIR  chr6:22216569-22216725 (-) |  |
|  |  |  |  |  |  |  |  |  |  | LINE (CR1) – L3  chr6:22216793-22217126 (-) |  |
|  |  |  |  |  | **25** | | chr6:98,626,637 | Simple Rep – (TG)n  chr6:98626163-98626211 (+) | - | tRNA (tRNA) – tRNA-Ile-ATA  chr6:98626784-98626822 (-) | 1 bp dup: chr6:98,626,637 |
|  |  |  |  |  |  |  |  | DNA (hAT-Charlie) – MER33  chr6:98625857-98626117 (-) |  | Simple Rep – (AC)n  chr6:98627045-98627075 (+) |  |
|  |  |  |  |  |  |  |  |  |  | LINE (RTE-BovB) – MamRTE1  chr6:98627447-98627566 (-) |  |
| 8 | **14** | t(6;8).2 | - | - | **27** | | chr8:111,765,702 | LINE (L1) – L1MC3  chr8:111764958-111765641 (+) | LTR (ERV1) – LTR1A1  chr8:111765642-111766427 (+) | LINE (L1) – L1MC3  chr8:111766428-111766465 (+) | - |
|  |  |  |  |  |  |  |  | LINE (L1) – L1PA7  chr8:111764568-111764957 (+) |  |  |  |
|  |  |  |  |  | **18** | | chr6:45,617,625 | LINE (CR1) – CR1-3_Croc  chr6:45617484-45617535 (+) | - | LINE (CR1) – L3b  chr6:45617689-45617777 (+) |  |
|  |  |  |  |  |  |  |  | SINE (MIR) – MIR  chr6:45617358-45617427 (-) |  |  |  |
|  |  |  |  |  |  |  |  | LINE (L2) – L2a  chr6:45617179-45617327 (-) |  |  |  |
|  |  |  |  |  |  |  |  | LINE (CR1) – L3  chr6:45616884-45617098 (-) |  |  |  |
| 8 | **15** | t(3;6).4 | - | - | **17** | | chr6:45,041,636 | LINE (L2) – L2a  chr6:45041041-45041539 (-) | - | SINE (MIR) – MIRb  chr6:45041792-45041949 (+) | - |
|  |  |  |  |  |  |  |  |  |  | SINE (Alu) – AluSq2  chr6:45042141-45042432 (+) |  |
|  |  |  |  |  |  |  |  |  |  | LINE (L1) – L1P2  chr6:45042540-45042557 (+) |  |
|  |  |  |  |  |  |  |  |  |  | LINE (L1) – L1PA4  chr6:45042558-45042608 (-) |  |
|  |  |  |  |  | **7** | | chr3:16,433,321 | SINE (Alu) – AluJr  chr3:16432556-16432684 (+) | - | SINE (Alu) – AluJb  chr3:16434242-16434361 (+) | 5 bp del: chr3:16,433,322-16,433,326 |
|  |  |  |  |  |  |  |  | SINE (Alu) – AluJb  chr3:16432236-16432545 (+) |  |  |  |
| **Junction Point** | | | | | **Breakpoint** | | | **Repetitive Elements** | | | **Obs.** |
| **der** | **#JP** | **JP ID** | **Ins** | **MH** | **#BP** | | **BP coord.** | **BP – 1 kb** | **BP** | **BP + 1 kb** |  |
| 8 | **16** | fus(3).3 | - | 1 (G) | **6** | | chr3:16,137,682 | DNA (hAT-Blackjack) – MER81  chr3:16137577-16137675 (-) | - | LTR (ERVL-MaLR) – MLT1I  chr3:11063055-11063352 (-) | - |
|  |  |  |  |  |  |  |  | ERVL-MaLR (LTR) – MLT1A  chr3:16137364-16137576 (+) |  |  |  |
|  |  |  |  |  |  |  |  | Simple Rep – (TC)n  chr3:16137338-16137363 (+) |  |  |  |
|  |  |  |  |  |  |  |  | DNA (hAT-Blackjack) – MLT1A  chr3:16137172-16137337 (+) |  | SINE (Alu) – AluY  chr3:11063424-11063728 (-) |  |
|  |  |  |  |  |  |  |  | DNA (hAT-Blackjack) – MER81  chr3:16137159-16137171 (-) |  |  |  |
|  |  |  |  |  | **1** | | chr3:11,062,692 | LTR (ERV1) – LTR12C  chr3:11061209-11062580 (+) | LINE (L2) – L2a  chr3:11062583-11063054 (-) | LTR (ERVL-MaLR) – MLT1I  chr3:11063055-11063352 (-) |  |
|  |  |  |  |  |  |  |  |  |  | SINE (Alu) – AluY  chr3:11063424-11063728 (-) |  |
| 14 | **17** | t(6;14).4 | - | 1 (G) | **29** | | chr14:65,594,523 | SINE (Alu) – AluSx  chr14:65593878-65594009 (-) | LTR (ERV1) – LTR28B  chr14:65594101-65595106 (-) | - | - |
|  |  |  |  |  |  |  |  | SINE (Alu) – AluSp  chr14:65593565-65593874 (-) |  |  |  |
|  |  |  |  |  | **11** | | chr6:12,214,526 | SINE (MIR) – MIR3  chr6:12214200-12214263 (-) | - | LTR (ERVL-MaLR) – THE1B  chr6:12214567-12214922 (-) |  |
|  |  |  |  |  |  |  |  | SINE (Alu) – AluJr  chr6:12213563-12213849 (+) |  | SINE (Alu) – AluJb  chr6:12215009-12215308 (-) |  |
|  |  |  |  |  |  |  |  | LINE (L1) – HAL1b  chr6:12213285-12213538 (-) |  | LINE (Penelope) – Penelope1_Vert chr6:12215317-12215393 (+) |  |
| 14 | **18** | fus(6).5 | 1 (T) | - | **14** | | chr6:21,627,913 | Simple Rep – (AAAT)n  chr6:21627895-21627908 (+) | SINE (Alu) – AluSq  chr6:21627909-21628165 (+) | Simple Rep – (AAAT)n  chr6:21628166-21628177 (+) | - |
|  |  |  |  |  |  |  |  | SINE (Alu) – AluY  chr6:21627286-21627593 (+) |  |  |  |
|  |  |  |  |  |  |  |  | Simple Rep – (ATTTA)n  chr6:21627168-21627207 (+) |  |  |  |
|  |  |  |  |  | **9** | | chr6:2,029,106 | Simple Rep – (T)n  chr6:2028983-2029009 (+) | - | DNA (hAT-Tip100) – MamTip1  chr6:2029869-2029957 (+) |  |
| **der** | **#JP** | **JP ID** | **Ins** | **MH** | **#BP** | | **BP coord.** | **BP – 1 kb** | **BP** | **BP + 1 kb** | **Obs.** |
| **Junction Point** | | | | | | **Breakpoint** | | **Repetitive Elements** | | |  |

JP: junction point; JP ID: junction point identification; Ins: insertion; SJP: sub-junction point; MH: microhomology; Del: deletion; BP: breakpoint; coord.: coordinate; SNVs: single nucleotide variants; Obs.: observation. All genomic coordinates are according to reference genome GRCh38/hg38.

**Supplementary Table 5. Difference between breakpoints found by Optical Genome Mapping (OGM) and Long-read Sequencing (lrGS).**

| **#JP** | **JP ID** | **#BP** | **BP OGM** | | **BP lrGS** | **BPs difference (bp)** | | **Sum of BPs difference (bp)** | | **Size of Uncertain OGM region (bp)** |
| --- | --- | --- | --- | --- | --- | --- | --- | --- | --- | --- |
| **1** | t(6;14).1 | **31** | chr14:68,069,448 | | chr14:68,065,355 | 4,093 | | 16,555 | | 16,218 |
|  |  | **16** | chr6:22,228,502 | | chr6:22,216,040 | 12,462 | |  |  |  |
| **2** | fus(6).1 | **17** | chr6:45,046,039 | | chr6:45,041,636 | 4,403 | | 8,622 | | 0 |
|  |  | **20** | chr6:46,289,346 | | chr6:46,293,565 | 4,219 | |  |  |  |
| **3** | t(3;6).1 | **19** | chr6:45,619,872 | | chr6:45,617,626 | 2,246 | | 5,595 | | 5,634 |
|  |  | **5** | chr3:12,162,249 | | chr3:12,158,900 | 3,349 | |  |  |  |
| **4**^a^ | fus(3).1 | **6** | chr3:16,128,547 | | chr3:16,137,682 | 9,135 | | 12,559 | | 12,606 |
|  |  | **1** | chr3:11,066,116 | | chr3:11,062,692 | 3,424 | |  |  |  |
| **5**^b^ | fus(3).2 | **2** | chr3:11,533,381 | | chr3:11,536,045 | 2,664 | | 4,128 | | 4,488 |
|  |  | **8** | chr3:16,434,791 | | chr3:16,433,327 | 1,464 | |  |  |  |
| **6** | t(6;8).1 | **26** | chr8:111,771,184 | | chr8:111,765,700 | 5,484 | | 6,752 | | 6,769 |
|  |  | **21** | chr6:46,294,839 | | chr6:46,293,571 | 1,268 | |  |  |  |
| **7** | t(3;6).2 | **22** | chr6:74,205,211 | | chr6:74,206,502 | 1,291 | | 3,531 | | 3,805 |
|  |  | **4** | chr3:12,154,778 | | chr3:12,157,018 | 2,240 | |  |  |  |
| **8** | t(3;6).3 | **3** | chr3:11,538,484 | | chr3:11,536,053 | 2,431 | | 9,217 | | 9,235 |
|  |  | **23** | chr6:74,213,306 | | chr6:74,206,520 | 6,786 | |  |  |  |
| **9** | t(6;14).2 | **24** | chr6:96,982,764 | | chr6:96,991,559 | 8,795 | | 18,162 | | 19,635 |
|  |  | **28** | chr14:65,603,889 | | chr14:65,594,522 | 9,367 | |  |  |  |
| **10** | t(6;14).3 | **30** | chr14:67,474,613 | | chr14:67,473,334 | 1,279 | | 2,633 | | 0 |
|  |  | **24** | chr6:96,992,913 | | chr6:96,991,559 | 1,354 | |  |  |  |
| **11** | fus(6).2 | **25** | chr6:98,619,024 | | chr6:98,626,637 | 7,613 | | 8,986 | | 8,973 |
|  |  | **12** | chr6:12,213,154 | | chr6:12,214,527 | 1,373 | |  |  |  |
| **12** | fus(6).3 | **10** | chr6:2,033,635 | | chr6:2,029,107 | 4,528 | | 8,400 | | 8,104 |
|  |  | **13** | chr6:21,631,784 | | chr6:21,627,912 | 3,872 | |  |  |  |
| **#JP** | **JP ID** | **#BP** | **BP OGM** | | **BP lrGS** | **BPs difference (bp)** | | **Sum of BPs difference (bp)** | | **Size of Uncertain OGM region (bp)** |
| **13** | fus(6).4 | **15** | chr6:22,214,492 | | chr6:22,216,039 | 1,547 | | 15,245 | | 15,505 |
|  |  | **25** | chr6:98,640,335 | | chr6:98,626,637 | 13,698 | |  |  |  |
| **14** | t(6;8).2 | **27** | chr8:111,760,447 | | chr8:111,765,702 | 5,255 | | 12,364 | | 12,584 |
|  |  | **18** | chr6:45,610,516 | | chr6:45,617,625 | 7,109 | |  |  |  |
| **15** | t(3;6).4 | **17** | chr6:45,047,282 | | chr6:45,041,636 | 5,646 | | 6,533 | | 6,851 |
|  |  | **7** | chr3:16,432,434 | | chr3:16,433,321 | 887 | |  |  |  |
| **16** | fus(3).3 | **6** | chr3:16,128,547 | | chr3:16,137,682 | 9,135 | | 12,559 | | 12,606 |
|  |  | **1** | chr3:11,066,116 | | chr3:11,062,692 | 3,424 | |  |  |  |
| **17** | t(6;14).4 | **29** | chr14:65,568,702 | | chr14:65,594,523 | 25,821 | | 60,676 | | 60,413 |
|  |  | **11** | chr6:12,249,381 | | chr6:12,214,526 | 34,855 | |  |  |  |
| **18** | fus(6).5 | **14** | chr6:21,625,655 | | chr6:21,627,913 | 2,258 | | 3,084 | | 3,015 |
|  |  | **9** | chr6:2,028,280 | | chr6:2,029,106 | 826 | |  |  |  |
|  | | | | **Average** | | | 5,989 | |  | |
|  | | | | **Standard Deviation** | | | 6,920 | |  | |

JP: junction point; JP ID: junction point identification; BP(s): breakpoint(s); ^a^Detected by OGM SV pipeline as a duplication; ^b^Detected by OGM SV pipeline as a deletion.

All genomic coordinates are according to reference genome GRCh38/hg38.

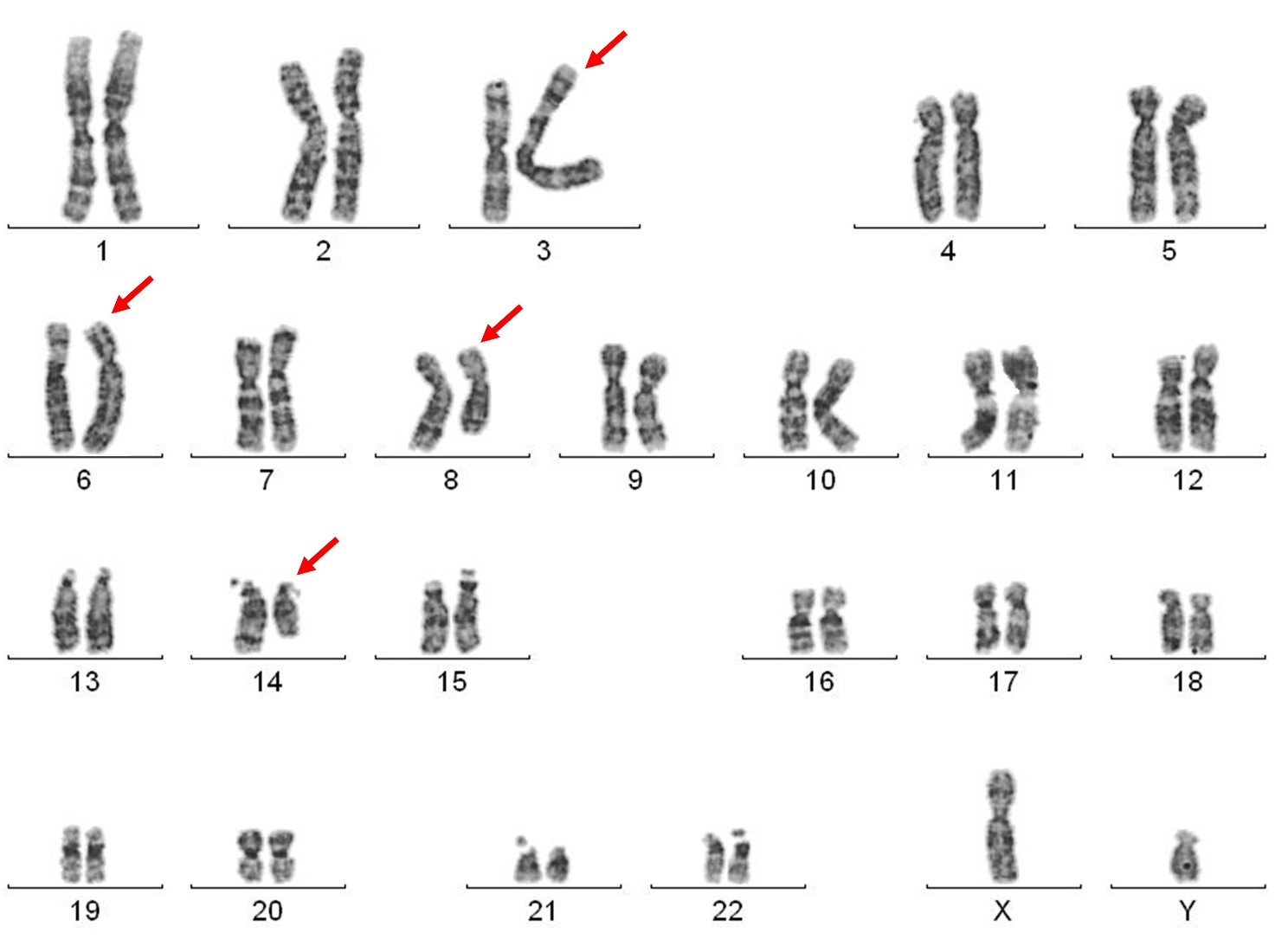

**Supplementary Figure 1. Karyotype of Patient 2.**

The red arrows highlight the four chromosomes involved in the patient’s rearrangement.

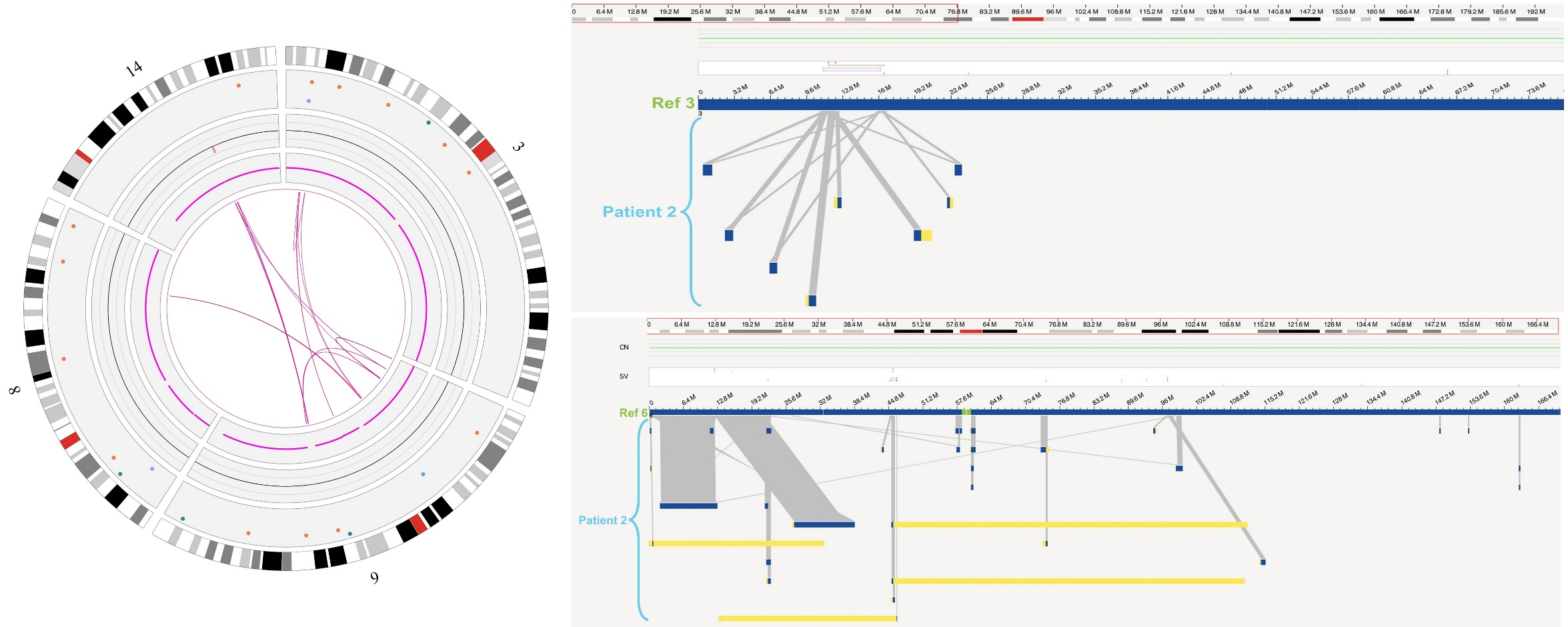

**Supplementary Figure 2. Optical Genome Mapping results of Patient 2.**

To the left, circos plot with OGM results of Patient 2 with the four involved chromosomes’ idiograms in the periphery and the pink lines in the middle representing intra- and inter-fusions. To the right, genome browser view of chromosomes 3 (top) and 6 (bottom). It is possible to see various patient maps mapping to different regions of the same chromosome as well as chrimeric maps, whose yellow parts map to a different chromosome.

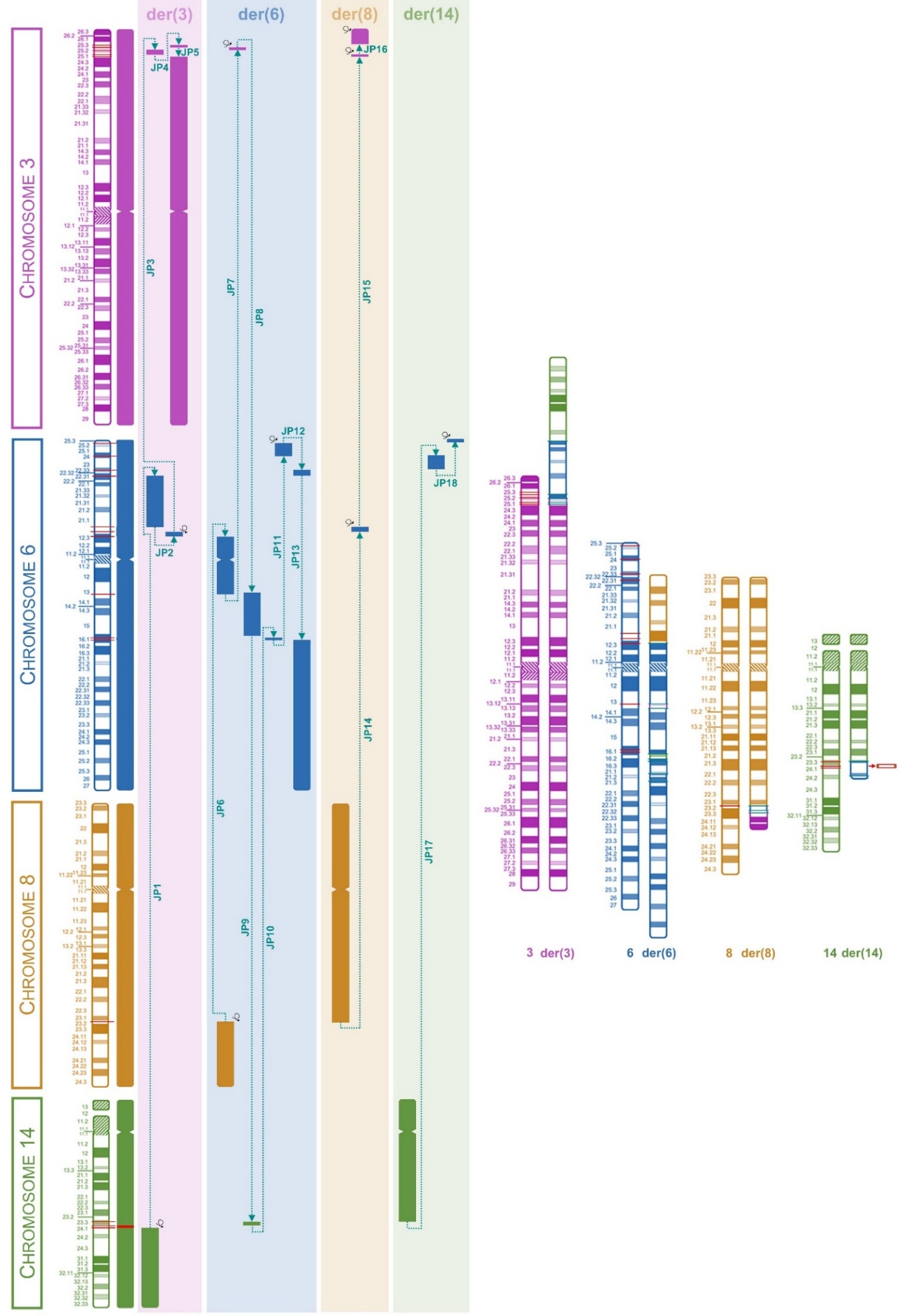

**Supplementary Figure 3. Patient 2’s resolved complex chromosomal rearrangement.**

To the left, a schematic representation of the junctions between all chromosome regions involved in the CCR. To the right, idiogram of the chromosomes involved in the complex chromosomal rearrangement. Red lines show the breakpoints in the normal chromosomes and cyan lines show the junction points in the derivative chromosomes.

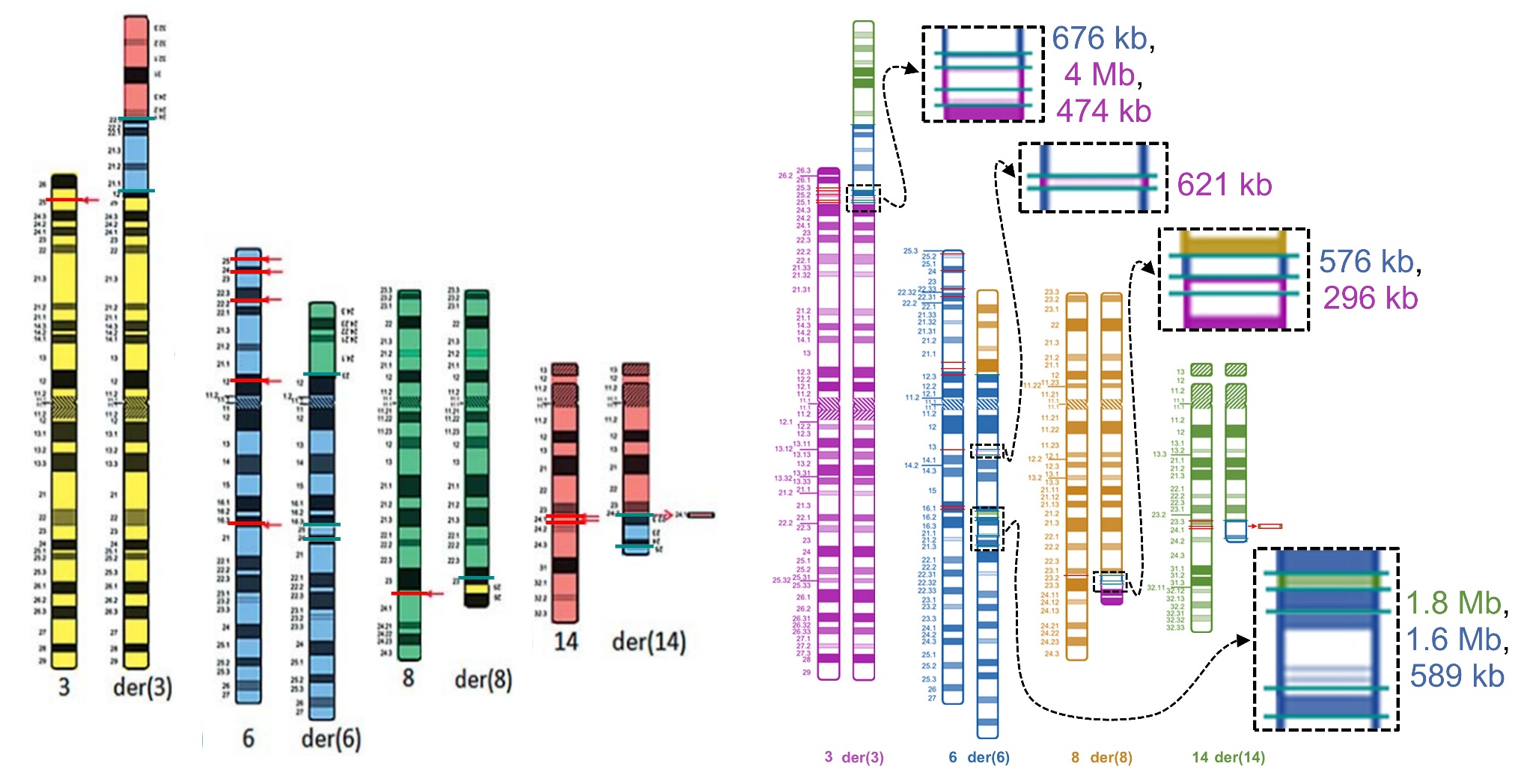

**Supplementary Figure 4. Comparison between the Patient 2's final rearrangement between Guilherme et al. 2013 and the present study.**

To the left, chromosome idiograms of the CGR from Guilherme et al. 2013. To the right, CGR chromosome idiograms revealed by this study, highlighting the regions that were only uncovered through OGM and lrGS.
