## Supplementary material for "Integrative optical genome mapping and long-read sequencing resolve constitutional complex rearrangements at nucleotide resolution": Supplementary File 3 - Patient 3.docx

Bruna Burssed, Bart van der Sanden, Kornelia Neveling, Wolfram Höps, Eveline Kamping, Ronald van Beek, Amber den Ouden, Ronny Derks, Raoul Timmermans, Eduardo Perrone, Marco Antonio Ramos, Fernanda Teixeira Bellucco, Alexander Hoischen, Maria Isabel Melaragno

**Supplementary File 3 – Additional information in Tables and Figures from Patient 3**

**TABLES.**

**FIGURES.**

**Supplementary Table 1. Patient 3’s phenotypes with Human Phenotype Ontology (HPO) terms.**

| **Phenotype** | **HPO** |
| --- | --- |
| **General** |  |
| Neurodevelopmental delay | HP:0012758 |
| speech delay | HP:0000750 |
| Intrauterine growth retardation | HP:0001511 |
| failure to thrive | HP:0001508 |
| **Skin** |  |
| Atopic dermatitis | HP:0001047 |
| **Behavior** |  |
| Irritability | HP:0000737 |
| Difficulty falling asleep | HP:0031354 |
| **Bones/Limbs** |  |
| Genu valgum | HP:0002857 |
| Pes planus | HP:0001763 |
| **Respiratory** |  |
| Laryngomalacia | HP:0001600 |
| **Eyes** |  |
| Poor eye contact | HP:0000817 |
| **Facial dysmorphisms** |  |
| Brachycephaly | HP:0000248 |
| Plagiocephaly | HP:0001357 |
| Triangular face | HP:0000325 |
| Prominent forehead | HP:0011220 |
| Facial edema | HP:0000282 |
| Thick eyebrow | HP:0000574 |
| Long eyelashes | HP:0000527 |
| Epicanthal folds | HP:0000286 |
| Upslanted palpebral fissure | HP:0000582 |
| Depressed nasal bridge | HP:0005280 |
| Anteverted nostrils | HP:0000463 |
| Small external ear | HP:0008772 |
| Low-set ears | HP:0000369 |

**Supplementary Table 2. Results of the techniques performed for the patient and final rearrangement.**

| **Technique** | **Result** |
| --- | --- |
| **Karyotype** | 46, XY, der(1)t(1;?)(q12;?), der(3)t(3;9)(p14;p23)t(3;11)(q13;q24),t(3;9)(q13;p23),  der(11)t(11;15)(q24;q?11.2), der(15)t(1;15)(q21;p13) hsr(1;15)(q21;p13)t(15;?)(q?11.2;?) |
| **Chromosomal Microarray Analysis** | arr[GRCh38] 11q23.3(115746405_118001992)×1 |
| **Optical Genome Mapping** | ogm[GRCh38] 3p24.1[29,909,121-30,428,590]×1, 3q13.11[103,861,058-109,242,307]×3, inv(3)(q13.11)(105,571,481_105,792,873), inv(3)(q25.32)(157,835,535_158,109,424), inv(9)(p21.3p13.3)(20,877,178_33,852,709), 11q23.3[115,756,201_118,962,20X]×1 |
| **Long-read Sequencing** | seq[GRCh38] 3p12.3[78,523,989_78,525,096]×1, 3q13.13[108,399,585_108,413,406]×1, 9p13.3[33,831,476_33,836,084]×1, 11q23.3[115,767,430_118,006,937]×1, 11q23.3[chr11:118,930,314_118,932,549]×1, 15q13.2[30,748,256_30,749,458]×1, inv(3)(q13.11)(105,573,426_105,790,639), inv(3)(q13.31)(114,157,417_114,289,729), inv(3)(q25.32)(157,835,921_158,124,238), inv(9)(p21.p13.3)(20,873,864_33,836,084) |
| **Final Rearrangement**  (colors represent similar breakpoints) | **der(1):** 1pter→1q25.1(chr1:175,239,230)::11q23.3(chr11:119,279,434)→11qter  **der(3):** 9pter→9p21.3(chr9:20,873,864)::3p12.3(chr3:78,525,097)→3q13.11(chr3:103,859,837)::3q13.13(chr3:108,399,584)→  3q13.11(chr3:105,790,639)::3q13.11(chr3:105,573,426)→3q13.11(chr3:105,790,639)::3q13.11(chr3:105,573,426)→  3q13.11(chr3:103,859,837)::3q13.13(chr3:109,246,317)→3q13.13(chr3:108,413,407):: 15p→15pter  **der(9):** 3qter→3q25.32(chr3:158,124,238)::3q25.32(chr3:157,835,920)→3q25.32(chr3:158,124,238)::3q25.32(chr3:157,835,920)→  3q25.31(chr3:156,977,787)::15q13.3(chr15:31,648,953)→15q13.3(chr15:30,749,459)::3q25.2(chr3:154,356,278)→  3q25.31(chr3:156,977,787)::11q23.3(chr11:118,932,550)→11q23.3(chr11:118,932,353)::3p22.3(chr3:33,892,569)→  3p12.3(chr3:78,523,988)::9p13.3(chr9:33,831,475)→9p21.3(chr9:20,873,864)::9p13.3(chr9:33,836,084)→9qter  **der(10):** 10pter→10q26.3(chr10:129,525,155)::3p24.1(chr3:30,427,217)→3p24.1(chr3:29,910,078)::10q26.3(chr10:129,525,155)→10qter  **der(11):** 11pter→11q23.3(chr11:115,767,429)::11q23.3(chr11:118,957,417)→11q23.3(chr11:119,278,574)::11q23.3(chr11:118,400,940)→  11q23.3(chr11:118,006,938)::15q21.3(chr15:53,606,243)→15qter  **der(15):** 1qter→1q25.1(chr1:175,239,230)::11q23.3(chr11:118,400,938)→11q23.3(chr11:118,930,313)::11q23.3(chr11:118,957,417)→  11q23.3(chr11:118,932,550)::15q21.3(chr15:53,606,243)→15q13.3(chr15:31,648,953)::3q25.2(chr3:154,356,278)→  3q13.31(chr3:114,289,729)::3q13.31(chr3:114,157,417)→3q13.31(chr3:114,289,729)::3q13.31(chr3:114,157,417)→  3q13.13(chr3:109,246,317)::15p→15q13.3(chr15:30,748,255)::3p22.3(chr3:33,892,542)→3p24.1(chr3:30,427,217)::  3p24.1(chr3:29,910,078)→3pter |
|  | 46,XY,t(1;3;9;10;11;15)(1pter→1q25.1::11q23.3→11qter;9pter→9p21.3::3p12.3→3q13.11::3q13.13→3q13.11::3q13.11→3q13.11::3q13.11→3q13.11::3q13.13→3q13.13::15p→15pter;3qter→3q25.32::3q25.32→3q25.32::3q25.32→3q25.31::15q13.3→15q13.3::3q25.2→3q25.31::11q23.3→11q23.3::3p22.3→3p12.3::9p13.3→9p21.3::9p13.3→9qter;10pter→10q26.3::3p24.1→3p24.1::10q26.3→10qter;11pter→11q23.3::11q23.3→11q23.3::11q23.3→11q23.3::15q21.3→15qter;1qter→1q25.1::11q23.3→11q23.3::11q23.3→11q23.3::15q21.3→15q13.3::3q25.2→3q13.31::3q13.31→3q13.31::3q13.31→3q13.13::15p→15q13.3::3p22.3→3p24.1::3p24.1→3pter) |

**Supplementary Table 3. Breakpoint coordinates found through long-read sequencing and their number (#) for identification.**

| **#BP** | **BP coordinate** |
| --- | --- |
| 23 | chr9:20,873,864 |
| 24 | chr9:20,873,865 |
| 25 | chr9:33,831,475 |
| 26 | chr9:33,836,084 |
| 27 | chr10:129,525,155 |
| 28 | chr11:115,767,429 |
| 29 | chr11:118,006,938 |
| 30 | chr11:118,400,940 |
| 31 | chr11:118,930,313 |
| 32 | chr11:118,932,353 |
| 33 | chr11:118,932,550 |
| 34 | chr11:118,957,417 |
| 35 | chr11:119,278,574 |
| 36 | chr11:119,279,434 |
| 37 | chr15p |
| 38 | chr15:30,748,255 |
| 39 | chr15:30,749,459 |
| 40 | chr15:31,648,952 |
| 41 | chr15:31,648,953 |
| 42 | chr15:53,606,243 |

| **#BP** | **BP coordinate** |
| --- | --- |
| 1 | chr1:175,239,230 |
| 2 | chr3:29,910,078 |
| 3 | chr3:30,427,217 |
| 4 | chr3:33,892,542 |
| 5 | chr3:33,892,569 |
| 6 | chr3:78,523,988 |
| 7 | chr3:78,525,097 |
| 8 | chr3:103,859,836 |
| 9 | chr3:103,859,837 |
| 10 | chr3:105,573,426 |
| 11 | chr3:105,790,639 |
| 12 | chr3:108,399,584 |
| 13 | chr3:108,413,407 |
| 14 | chr3:109,246,317 |
| 15 | chr3:114,157,417 |
| 16 | chr3:114,289,729 |
| 17 | chr3:154,356,278 |
| 18 | chr3:156,977,787 |
| 19 | chr3:157,835,920 |
| 20 | chr3:157,835,921 |
| 21 | chr3:158,124,237 |
| 22 | chr3:158,124,238 |

**Supplementary Table 4. Junction points and breakpoints details found on the long-read sequencing analysis and annotation of Repetitive Elements in regions surrounding the breakpoints.**

| **Junction Point** | | | | | | | **Breakpoint** | | **Repetitive Elements** | | | | **Obs.** |
| --- | --- | --- | --- | --- | --- | --- | --- | --- | --- | --- | --- | --- | --- |
| **der** | **#JP** | **JP ID** | **Ins** | **MH** | **#SJP** | **Ins/MH** | **#BP** | **BP coord.** | | **BP - 1 kb** | **BP** | **BP + 1 kb** |  |
| 1 | **1** | t(1;11).1 | - | - | - | - | **1** | chr1:175,239,230 | | - | LINE (L1) - L1PBa1  chr1:175234071-175240273 (+) | - | - |
|  |  |  |  |  |  |  | **36** | chr11:119,279,434 | | SINE (Alu) - FLAM_C  chr11:119279218-119279361 (+) | SINE (Alu) - AluJo  chr11:119279365-119279660 (+) | - |  |
|  |  |  |  |  |  |  |  |  |  | SINE (MIR) - MIR  chr11:119278813-119279056 (-) |  |  |  |
|  |  |  |  |  |  |  |  |  |  | Simple Rep - (ATG)n  chr11:119278643-119278677 (+) |  |  |  |
| 3 | **2** | t(3;9).1 | - | 3 (CAC) | - | - | **24** | chr9:20,873,865 | | LINE (L1) - L1MB8  chr9:20872649-20873224 (-) | - | - | 2bp dup: chr9:20,873,864-20,873,865 |
|  |  |  |  |  |  |  | **7** | chr3:78,525,097 | | LINE (L1) - L1M4a1  chr3:78523687-78525037 (+) | - | Simple Rep - (TTTTA)n  chr3:78525532-78525595 (+) |  |
|  |  |  |  |  |  |  |  |  |  |  |  | SINE (Alu) - AluJo  chr3:78525598-78525886 (-) |  |
|  |  |  |  |  |  |  |  |  |  |  |  | LINE (L1) - L1MA6  chr3:78525899-78526736 (-) |  |
| 3 | **3** | fus(3).1 | - | - | - | - | **8** | chr3:103,859,836 | | Simple Rep - (TG)n  chr3:103859554-103859626 (+) | - | LINE (L1) - L1ME4a  chr3:103860608-103860910 (+) | - |
|  |  |  |  |  |  |  |  |  |  | LINE (L1) - L1ME4a  chr3:103859435-103859538 (+) |  |  |  |
|  |  |  |  |  |  |  |  |  |  | LTR (ERV1) - LTR16B2  chr3:103858901-103859334 (+) |  |  |  |
|  |  |  |  |  |  |  |  |  |  | Simple Rep - (TTA)n  chr3:103858840-103858882 (+) |  |  |  |
|  |  |  |  |  |  |  | **12** | chr3:108,399,584 | | - | - | LTR (ERVL-MaLR) - MLT1J2  chr3:108399996-108400316 (+) |  |
|  |  |  |  |  |  |  |  |  |  |  |  | SINE (Alu) - AluSg  chr3:108400545-108400827 (+) |  |
| 3 | **4** | fus(3).2 | - | 1 (C) | - | - | **10** | chr3:105,573,426 | | SINE (Alu) - AluSz  chr3:105573039-105573353 (+) | - | Simple Rep - (TA)n  chr3:105573450-105573564 (+) | - |
|  |  |  |  |  |  |  |  |  |  | LINE (L1) - L1PA7  chr3:105572088-105572736 (-) |  |  |  |
|  |  |  |  |  |  |  | **11** | chr3:105,790,639 | | - | LINE (L2) - L2c  chr3:105790488-105790640 (-) | - |  |
| 3 | **5** | fus(3).3 | - | 1 (C) | - | - | **10** | chr3:105,573,426 | | SINE (Alu) - AluSz  chr3:105573039-105573353 (+) | - | Simple Rep - (TA)n  chr3:105573450-105573564 (+) | - |
|  |  |  |  |  |  |  |  |  |  | LINE (L1) - L1PA7  chr3:105572088-105572736 (-) |  |  |  |
|  |  |  |  |  |  |  | **11** | chr3:105,790,639 | |  | LINE (L2) - L2c  chr3:105790488-105790640 (-) | - |  |
| **Junction Point** | | | | | | | **Breakpoint** | | | **Repetitive Elements** | | | **Obs.** |
| **der** | **#JP** | **JP ID** | **Ins** | **MH** | **#SJP** | **Ins/MH** | **#BP** | **BP coord.** | | **BP - 1 kb** | **BP** | **BP + 1 kb** |  |
| 3 | **6** | fus(3).4 | - | - | - | - | **9** | chr3:103,859,837 | | Simple Rep - (TG)n  chr3:103859554-103859626 (+) | - | LINE (L1) - L1ME4a  chr3:103860608-103860910 (+) | - |
|  |  |  |  |  |  |  |  |  |  | LINE (L1) - L1ME4a  chr3:103859435-103859538 (+) |  |  |  |
|  |  |  |  |  |  |  |  |  |  | LTR (ERV1) - LTR16B2  chr3:103858901-103859334 (+) |  |  |  |
|  |  |  |  |  |  |  |  |  |  | Simple Rep - (TTA)n  chr3:103858840-103858882 (+) |  |  |  |
|  |  |  |  |  |  |  | **14** | chr3:109,246,317 | | LTR (ERVL-MaLR) - THE1A  chr3:109245490-109245847 (-) | LTR (ERVL-MaLR) - THE1Aint  chr3:109245848-109247409 (-) | - |  |
| 3 | **7** | t(3;15).1 | - | - | - | - | **13** | chr3:108,413,407 | | DNA (hAT-Blackjack) - MER81  chr3:108412511-108412578 (-) | SINE (5S-Deu-L2) - AmnSINE1  chr3:108413282-108413447 (-) | LINE (L2) - L2b  chr3:108413924-108414135 (+) | - |
|  |  |  |  |  |  |  |  |  |  | LTR (ERVL-MaLR) - THE1C  chr3:108412141-108412510 (+) |  |  |  |
|  |  |  |  |  |  |  | **37** | chr15p | | - | - | - |  |
| 9 | **8** | fus(3).5 | - | - | - | - | **22** | chr3:158,124,238 | | SINE (Alu) - AluJr  chr3:158123318-158123597 (-) | LTR (ERVL-MaLR) - MLT1H  chr3:158124000-158124295 (-) | LINE (L1) - L1M4  chr3:158124318-158124806 (-) | - |
|  |  |  |  |  |  |  |  |  |  | SINE (Alu) - AluSq2  chr3:158123032-158123306 (-) |  | SINE (Alu) - AluJr  chr3:158124807-158125118 (-) |  |
|  |  |  |  |  |  |  |  |  |  |  |  | LINE (L1) - L1M4  chr3:158125119-158126874 (-) |  |
|  |  |  |  |  |  |  | **20** | chr3:157,835,921 | | LINE (L2) - L2c  chr3:157835443-157835778 (+) | - | - |  |
|  |  |  |  |  |  |  |  |  |  | LTR (ERV1) - MER31B  chr3:157834977-157835442 (+) |  |  |  |
|  |  |  |  |  |  |  |  |  |  | LINE (L2) - L2c  chr3:157834497-157834976 (+) |  |  |  |
| 9 | **9** | fus(3).6 | - | - | - | - | **21** | chr3:158,124,237 | | SINE (Alu) - AluJr  chr3:158123318-158123597 (-) | LTR (ERVL-MaLR) - MLT1H  chr3:158124000-158124295 (-) | LINE (L1) - L1M4  chr3:158124318-158124806 (-) | - |
|  |  |  |  |  |  |  |  |  |  | SINE (Alu) - AluSq2  chr3:158123032-158123306 (-) |  | SINE (Alu) - AluJr  chr3:158124807-158125118 (-) |  |
|  |  |  |  |  |  |  |  |  |  |  |  | LINE (L1) - L1M4  chr3:158125119-158126874 (-) |  |
|  |  |  |  |  |  |  | **19** | chr3:157,835,920 | | LINE (L2) - L2c  chr3:157835443-157835778 (+) | - | - |  |
|  |  |  |  |  |  |  |  |  |  | LTR (ERV1) - MER31B  chr3:157834977-157835442 (+) |  |  |  |
|  |  |  |  |  |  |  |  |  |  | LINE (L2) - L2c  chr3:157834497-157834976 (+) |  |  |  |
| 9 | **10** | t(3;15).2 | - | - | - | - | **18** | chr3:156,977,787 | | SINE (MIR) - MIRb  chr3:156976745-156976855 (+) | - | SINE (MIR) - MIRc  chr3:156978118-156978176 (+) | - |
|  |  |  |  |  |  |  | **40** | chr15:31,648,952 | | LTR (ERV1) - LTR39  chr15:31648149-31648754 (+) | SINE (Alu) - AluSx  chr15:31648755-31649060 (-) | LTR (ERV1) - LTR39  chr15:31649061-31649244 (+) |  |
|  |  |  |  |  |  |  |  |  |  |  |  | LINE (L1) - L1ME1  chr15:31649710-31651184 (+) |  |
| **Junction Point** | | | | | | | **Breakpoint** | | | **Repetitive Elements** | | | **Obs.** |
| **der** | **#JP** | **JP ID** | **Ins** | **MH** | **#SJP** | **Ins/MH** | **#BP** | **BP coord.** | | **BP - 1 kb** | **BP** | **BP + 1 kb** |  |
| 9 | **11** | t(3;15).3 | - | - | - | - | **39** | chr15:30,749,459 | | DNA (hAT-Charlie) - MER58A  chr15:30749107-30749324 (-) | - | LTR (ERVL-MaLR) - MLT1A0  chr15:30749927-30750095 (-) | - |
|  |  |  |  |  |  |  |  |  |  | SINE (MIR) - MIRb  chr15:30748807-30748944 (-) |  | LTR (ERVL-MaLR) - MLT1A0  chr15:30750100-30750168 (-) |  |
|  |  |  |  |  |  |  | **17** | chr3:154,356,278 | | LINE (L1) - L1MB8  chr3:154356002-154356198 (+) | LTR (ERVL-MaLR) - THE1D  chr3:154356203-154356292 (-) | LINE (L1) - L1PA4  chr3:154356315-154358302 (+) |  |
|  |  |  |  |  |  |  |  |  |  | SINE (Alu) - AluSg  chr3:154355030-154355325 (-) |  |  |  |
| 9 | **12** | fus(3).7 | 195 | - | 12-1 | MH:  2 (CA) | **18** | chr3:156,977,787 | | SINE (MIR) - MIRb  chr3:156976745-156976855 (+) | - | SINE (MIR) - MIRc  chr3:156978118-156978176 (+) | - |
|  |  |  |  |  |  |  | **33** | chr11:118,932,550 | | SINE (Alu) - AluJb  chr11:118931915-118932223 (+) | SINE (Alu) - AluSg  chr11:118932293-118932606 (-) | SINE (Alu) - AluY  chr11:118932618-118932934 (-) |  |
|  |  |  |  |  |  |  |  |  |  | LINE (L2) - L2b  chr11:118931592-118931880 (-) |  | SINE (MIR) - MIR3  chr11:118933000-118933088 (+) |  |
|  |  |  |  |  |  |  |  |  |  |  |  | SINE (MIR) - MIRc  chr11:118933430-118933557 (-) |  |
|  |  |  |  |  | 12-2 | MH:  1 (C) | **32** | chr11:118,932,353 | | SINE (Alu) - AluJb  chr11:118931915-118932223 (+) | SINE (Alu) - AluSg  chr11:118932293-118932606 (-) | SINE (Alu) - AluY  chr11:118932618-118932934 (-) |  |
|  |  |  |  |  |  |  |  |  |  | LINE (L2) - L2b  chr11:118931592-118931880 (-) |  |  |  |
|  |  |  |  |  |  |  |  |  |  | Simple Rep - (AAAC)n  chr11:118931529-118931551 (+) |  | SINE (MIR) - MIR3  chr11:118933000-118933088 (+) |  |
|  |  |  |  |  |  |  |  |  |  | srpRNA - 7SLRNA  chr11:118931216-118931442 (+) |  |  |  |
|  |  |  |  |  |  |  | **5** | chr3:33,892,569 | | LINE (L1) - L1PB3  chr3:33891412-33892329 (-) | LTR (ERV1) - MER21A  chr3:33892330-33892767 (-) | LTR (ERV1) - MER21-int  chr3:33892768-33892801 (-) |  |
|  |  |  |  |  |  |  |  |  |  |  |  | Simple Rep - (ATCT)n  chr3:33892802-33892856 (+) |  |
|  |  |  |  |  |  |  |  |  |  |  |  | LTR (ERV1) - MER21-int  chr3:33892857-33893193 (-) |  |
|  |  |  |  |  |  |  |  |  |  |  |  | LINE (L1) - L1PB3  chr3:33893195-33894318 (-) |  |
| 9 | **13** | t(3;9).2 | - | 2 (TG) | - | - | **6** | chr3:78,523,988 | | LINE (L1) - L1MC4  chr3:78523550-78523673 (+) | LINE (L1) - L1M4a1  chr3:78523687-78525037 (+) | - | - |
|  |  |  |  |  |  |  |  |  |  | LINE (L2) - L2a  chr3:78523339-78523538 (-) |  |  |  |
|  |  |  |  |  |  |  |  |  |  | LINE (L1) - L1MB2  chr3:78523192-78523289 (+) |  |  |  |
|  |  |  |  |  |  |  | **25** | chr9:33,831,475 | | SINE (Alu) - FLAM_C  chr9:33831160-33831292 (+) | SINE (Alu) - AluSx  chr9:33831418-33831644 (-) | SINE (Alu) - AluSx1  chr9:33831651-33831960 (-) |  |
|  |  |  |  |  |  |  |  |  |  | Simple Rep - (A)n  chr9:33831075-33831100 (+) |  | SINE (Alu) - AluSp  chr9:33832027-33832333 (+) |  |
|  |  |  |  |  |  |  |  |  |  | scRNA - HY3  chr9:33830978-33831074 (+) |  | SINE (Alu) - AluY  chr9:33832375-33832688 (+) |  |
|  |  |  |  |  |  |  |  |  |  | SINE (Alu) - AluSx  chr9:33830489-33830785 (+) |  |  |  |
| **Junction Point** | | | | | | | **Breakpoint** | | | **Repetitive Elements** | | | **Obs.** |
| **der** | **#JP** | **JP ID** | **Ins** | **MH** | **#SJP** | **Ins/MH** | **#BP** | **BP coord.** | | **BP - 1 kb** | **BP** | **BP + 1 kb** |  |
| 9 | **14** | fus(9) | - | - | - | - | **23** | chr9:20,873,864 | | LINE (L1) - L1MB8  chr9:20872649-20873224 (-) | - | - | - |
|  |  |  |  |  |  |  | **26** | chr9:33,836,084 | | DNA (TcMar-Tigger) - Tigger3b  chr9:33835891-33836017 (+) | SINE (Alu) - AluSx1  chr9:33836021-33836198 (+) | DNA (TcMar-Tigger) - Tigger3c  chr9:33836200-33836412 (+) |  |
|  |  |  |  |  |  |  |  |  |  | SINE (Alu) - AluJo  chr9:33835600-33835889 (+) |  | LINE (L1) - L1MB8  chr9:33836414-33836474 (-) |  |
|  |  |  |  |  |  |  |  |  |  | LINE (L1) - L1MC  chr9:33835493-33835569 (-) |  | SINE (Alu) - AluSz6  chr9:33836475-33836769 (+) |  |
|  |  |  |  |  |  |  |  |  |  | SINE (Alu) - AluSx1  chr9:33835146-33835446 (-) |  | LINE (L1) - HAL1  chr9:33836784-33837216 (-) |  |
| 10 | **15** | t(3;10).1 | - | - | - | - | **27** | chr10:129,525,155 | | SINE (Alu) - AluY  chr10:129524583-129524889 (-) | - | - | - |
|  |  |  |  |  |  |  | **3** | chr3:30,427,217 | | LINE (L1) - L1MB8  chr3:30426308-30426827 (-) | LINE (L1) - L1ME3A  chr3:30426835-30427469 (-) | SINE (Alu) - AluJr  chr3:30427470-30427746 (+) |  |
|  |  |  |  |  |  |  |  |  |  | LINE (L1) - L1MB8  chr3:30425134-30426274 (-) |  | LINE (L1) - L1ME3A  chr3:30427747-30428314 (-) |  |
| 10 | **16** | t(3;10).2 | - | 9 | - | - | **2** | chr3:29,910,078 | | - | - | Simple Rep - (TATATG)n  chr3:29910748-29910782 (+) | (TATGTTAA-T) |
|  |  |  |  |  |  |  |  |  |  |  |  | LINE (L2) - L2c  chr3:29911010-29911239 (-) |  |
|  |  |  |  |  |  |  | **27** | chr10:129,525,155 | | SINE (Alu) - AluY  chr10:129524583-129524889 (-) | - | - |  |
| 11 | **17** | fus(11).1 | - | - | - | - | **28** | chr11:115,767,429 | | - | - | LINE (L2) - L2c  chr11:115767760-115768179 (+) | - |
|  |  |  |  |  |  |  |  |  |  |  |  | LINE (L2) - L2c  chr11:115768182-115768285 (+) |  |
|  |  |  |  |  |  |  | **34** | chr11:118,957,417 | | - | - | LINE (L2) - L2c  chr11:118957831-118957930 (+) |  |
|  |  |  |  |  |  |  |  |  |  |  |  | Simple Rep - (TGGT)n  chr11:118957946-118958023 (+) |  |
| 11 | **18** | fus(11).2 | - | 3 (ACC) | - | - | **35** | chr11:119,278,574 | | - | - | Simple Rep – (ATG)n  chr11:119278643-119278677 (+) | 3bp dup:  chr11:118,400,938-118,400,940 |
|  |  |  |  |  |  |  |  |  |  |  |  | SINE (MIR) – MIR  chr11:119278813-119279056 (-) |  |
|  |  |  |  |  |  |  |  |  |  |  |  | SINE (Alu) – FLAM_C  chr11:119279218-119279361 (+) |  |
|  |  |  |  |  |  |  |  |  |  |  |  | SINE (Alu) – AluJo  chr11:119279365-119279660 (+) |  |
|  |  |  |  |  |  |  | **30** | chr11:118,400,940 | | SINE (MIR) – MIR  chr11:118400583-118400620 (+) | SINE (Alu) – AluSx1  chr11:118400840-118401118 (+) | - |  |
|  |  |  |  |  |  |  |  |  |  | SINE (Alu) – FLAM_A  chr11:118399918-118400044 (+) |  |  |  |
| **Junction Point** | | | | | | | **Breakpoint** | | | **Repetitive Elements** | | | **Obs.** |
| **der** | **#JP** | **JP ID** | **Ins** | **MH** | **#SJP** | **Ins/MH** | **#BP** | **BP coord.** | | **BP - 1 kb** | **BP** | **BP + 1 kb** |  |
| 11 | **19** | t(11;15).1 | - | - | - | - | **29** | chr11:118,006,938 | | LINE (L2) – L2b  chr11:118006464-118006589 (+) | LINE (L2) – L2c  chr11:118006935-118007022 (-) | Simple Rep – (AC)n  chr11:118007043-118007077 (+) | - |
|  |  |  |  |  |  |  |  |  |  |  |  | Simple Rep – (CCCACC)n  chr11:118007090-118007121 (+) |  |
|  |  |  |  |  |  |  |  |  |  | SINE (MIR) – MIRc  chr11:118006345-118006458 (+) |  | SINE (MIR) – MIR3  chr11:118007270-118007387 (-) |  |
|  |  |  |  |  |  |  |  |  |  |  |  | SINE (Alu) – AluSc  chr11:118007401-118007693 (-) |  |
|  |  |  |  |  |  |  | **42** | chr15:53,606,243 | | SINE (MIR) – MIR1_Amn  chr15:53605923-53606059 (-) | - | LINE (L2) – L2a  chr15:53606405-53606912 (-) |  |
|  |  |  |  |  |  |  |  |  |  | SINE (Alu) – AluSz  chr15:53605574-53605839 (+) |  |  |  |
|  |  |  |  |  |  |  |  |  |  | LINE (L1) – L1PA13  chr15:53601066-53605506 (+) |  |  |  |
| 15 | **20** | t(1;11) 2 | - | 1 (T) | - | - | **1** | chr1:175,239,230 | | - | LINE (L1) - L1PBa1  chr1:175234071-175240273 (+) | - | - |
|  |  |  |  |  |  |  | **29** | chr11:118,400,938 | | SINE (MIR) – MIR  chr11:118400583-118400620 (+) | SINE (Alu) – AluSx1  chr11:118400840-118401118 (+) | - |  |
|  |  |  |  |  |  |  |  |  |  | SINE (Alu) – FLAM_A  chr11:118399918-118400044 (+) |  |  |  |
| 15 | **21** | fus(11).2 | - | - | - | - | **31** | chr11:118,930,313 | | Simple Rep – (TCCCCC)n  chr11:118929552-118929596 (+) | - | srpRNA – 7SLRNA  chr11:118931216-118931442 (+) | - |
|  |  |  |  |  |  |  | **34** | chr11:118,957,417 | | - | - | LINE (L2) - L2c  chr11:118957831-118957930 (+) |  |
|  |  |  |  |  |  |  |  |  |  |  |  | Simple Rep - (TGGT)n  chr11:118957946-118958023 (+) |  |
| 15 | **22** | t(11;15).2 | - | - | - | - | **33** | chr11:118,932,550 | | SINE (Alu) – AluJb  chr11:118931915-118932223 (+) | SINE (Alu) – AluSg  chr11:118932293-118932606 (-) | SINE (Alu) – AluY  chr11:118932618-118932934 (-) | - |
|  |  |  |  |  |  |  |  |  |  | LINE (L2) – L2b  chr11:118931592-118931880 (-) |  | SINE (MIR) – MIR3  chr11:118933000-118933088 (+) |  |
|  |  |  |  |  |  |  |  |  |  |  |  | SINE (MIR) – MIRc  chr11:118933430-118933557 (-) |  |
|  |  |  |  |  |  |  | **42** | chr15:53,606,243 | | SINE (MIR) – MIR1_Amn  chr15:53605923-53606059 (-) | - | LINE (L2) – L2a  chr15:53606405-53606912 (-) |  |
|  |  |  |  |  |  |  |  |  |  | SINE (Alu) – AluSz  chr15:53605574-53605839 (+) |  |  |  |
|  |  |  |  |  |  |  |  |  |  | LINE (L1) – L1PA13  chr15:53601066-53605506 (+) |  |  |  |
| 15 | **23** | t(3;15).4 | - | 1 (G) | - | - | **41** | chr15:31,648,953 | | LTR (ERV1) - LTR39  chr15:31648149-31648754 (+) | SINE (Alu) - AluSx  chr15:31648755-31649060 (-) | LTR (ERV1) - LTR39  chr15:31649061-31649244 (+) | - |
|  |  |  |  |  |  |  |  |  |  |  |  | LINE (L1) - L1ME1  chr15:31649710-31651184 (+) |  |
|  |  |  |  |  |  |  | **17** | chr3:154,356,278 | | LINE (L1) - L1MB8  chr3:154356002-154356198 (+) | LTR (ERVL-MaLR) - THE1D  chr3:154356203-154356292 (-) | LINE (L1) - L1PA4  chr3:154356315-154358302 (+) |  |
|  |  |  |  |  |  |  |  |  |  | SINE (Alu) - AluSg  chr3:154355030-154355325 (-) |  |  |  |
| **Junction Point** | | | | | | | **Breakpoint** | | | **Repetitive Elements** | | | **Obs.** |
| **der** | **#JP** | **JP ID** | **Ins** | **MH** | **#SJP** | **Ins/MH** | **#BP** | **BP coord.** | | **BP - 1 kb** | **BP** | **BP + 1 kb** |  |
| 15 | **24** | fus(3).8 | - | - | - | - | **16** | chr3:114,289,729 | | LINE (L1) – L1MD  chr3:114289346-114289683 (-) | - | LTR (ERV1) – MER31A  chr3:114289876-114290342 (+) | - |
|  |  |  |  |  |  |  |  |  |  | SINE (Alu) – AluSp  chr3:114288984-114289256 (+) |  |  |  |
|  |  |  |  |  |  |  |  |  |  | LINE (L1) – L1M4  chr3:114288444-114288859 (-) |  |  |  |
|  |  |  |  |  |  |  | **15** | chr3:114,157,417 | | SINE (Alu) – AluSx1  chr3:114157044-114157353 (-) | - | DNA (TcMar-Tigger) – Tigger8  chr3:114157747-114157820 (+) |  |
|  |  |  |  |  |  |  |  |  |  |  |  | SINE (Alu) – AluSp  chr3:114157821-114158124 (+) |  |
|  |  |  |  |  |  |  |  |  |  | Simple Rep – (CTTT)n  chr3:114156713-114157042 (+) |  | SINE (Alu) – FLAM_A  chr3:114158132-114158264 (+) |  |
|  |  |  |  |  |  |  |  |  |  |  |  | LINE (L1) – L1MB7  chr3:114158274-114158380 (+) |  |
| 15 | **25** | fus(3).9 | - | - | - | - | **16** | chr3:114,289,729 | | LINE (L1) – L1MD  chr3:114289346-114289683 (-) | - | LTR (ERV1) – MER31A  chr3:114289876-114290342 (+) | - |
|  |  |  |  |  |  |  |  |  |  | SINE (Alu) – AluSp  chr3:114288984-114289256 (+) |  |  |  |
|  |  |  |  |  |  |  |  |  |  | LINE (L1) – L1M4  chr3:114288444-114288859 (-) |  |  |  |
|  |  |  |  |  |  |  | **15** | chr3:114,157,417 | | SINE (Alu) – AluSx1  chr3:114157044-114157353 (-) | - | DNA (TcMar-Tigger) – Tigger8  chr3:114157747-114157820 (+) |  |
|  |  |  |  |  |  |  |  |  |  |  |  | SINE (Alu) – AluSp  chr3:114157821-114158124 (+) |  |
|  |  |  |  |  |  |  |  |  |  | Simple Rep – (CTTT)n  chr3:114156713-114157042 (+) |  | SINE (Alu) – FLAM_A  chr3:114158132-114158264 (+) |  |
|  |  |  |  |  |  |  |  |  |  |  |  | LINE (L1) – L1MB7  chr3:114158274-114158380 (+) |  |
| 15 | **26** | t(3;15).5 | - | - | - | - | **14** | chr3:109,246,317 | | LTR (ERVL-MaLR) - THE1A  chr3:109245490-109245847 (-) | LTR (ERVL-MaLR) - THE1Aint  chr3:109245848-109247409 (-) | - | - |
|  |  |  |  |  |  |  | **37** | chr15p | | - | - | - |  |
| 15 | **27** | t(3;15).6 | - | 1 (T) | -- | - | **38** | chr15:30,748,255 | | SINE (MIR) – MIRb  chr15:30747430-30747643 (-) | SINE (Alu) – AluY  chr15:30748008-30748316 (+) | SINE (MIR) – MIRb  chr15:30748807-30748944 (-) | 25bp del:  chr3:33,892,543- 33,892,568 |
|  |  |  |  |  |  |  |  |  |  |  |  | DNA (hAT-Charlie) - MER58A  chr15:30749107-30749324 (-) |  |
|  |  |  |  |  |  |  | **4** | chr3:33,892,542 | | LINE (L1) - L1PB3  chr3:33891412-33892329 (-) | LTR (ERV1) - MER21A  chr3:33892330-33892767 (-) | LTR (ERV1) - MER21-int  chr3:33892768-33892801 (-) |  |
|  |  |  |  |  |  |  |  |  |  |  |  | Simple Rep - (ATCT)n  chr3:33892802-33892856 (+) |  |
|  |  |  |  |  |  |  |  |  |  |  |  | LTR (ERV1) - MER21-int  chr3:33892857-33893193 (-) |  |
|  |  |  |  |  |  |  |  |  |  |  |  | LINE (L1) - L1PB3  chr3:33893195-33894318 (-) |  |
| 15 | **28** | fus(3).10 | - | 1 (T) | - | -  JP: junction point; JP ID: junction point identification; Ins: insertion; SJP: subjunction point; MH: microhomology; BP: breakpoint; coord.: coordinate; SNVs: single nucleotide variants;  Obs.: observation. All genomic coordinates are according to reference genome GRCh38/hg38.  - | **3** | chr3:30,427,217 | | LINE (L1) - L1MB8  chr3:30426308-30426827 (-) | LINE (L1) - L1ME3A  chr3:30426835-30427469 (-) | SINE (Alu) - AluJr  chr3:30427470-30427746 (+) | - |
|  |  |  |  |  |  |  |  |  |  | LINE (L1) - L1MB8  chr3:30425134-30426274 (-) |  | LINE (L1) - L1ME3A  chr3:30427747-30428314 (-) |  |
|  |  |  |  |  |  |  | **2** | chr3:29,910,078 | | - | - | Simple Rep - (TATATG)n  chr3:29910748-29910782 (+) |  |
|  |  |  |  |  |  |  |  |  |  |  |  | LINE (L2) - L2c  chr3:29911010-29911239 (-) |  |

**Supplementary Table 5. Difference between breakpoints found by Optical Genome Mapping (OGM) and Long-read Sequencing (lrGS).**

| **#JP** | **JP ID** | **#BP** | **BP OGM** | **BP lrGS** | **BPs difference (bp)** | | **Sum of BPs difference (bp)** | **Size of Uncertain OGM region (bp)** | **OGM call** |
| --- | --- | --- | --- | --- | --- | --- | --- | --- | --- |
| 1 | t(1;11).1 | **1** | chr1:175,221,828 | chr1:175,239,230 | 17,402 | | 21,270 | 21,310 | Yes |
|  |  | 36 | chr11:119,283,302 | chr11:119,279,434 | 3,868 | |  |  |  |
| 2 | t(3;9).1 | 24 | chr9:20,873,686 | chr9:20,873,865 | 179 | | 2,496 | 2,487 | Yes |
|  |  | 7 | chr3:78,527,414 | chr3:78,525,097 | 2,317 | |  |  |  |
| 3 | fus(3).1 | 8 | chr3:103,857,730 | chr3:103,859,836 | 2,106 | | 4,208 | 0 | No |
|  |  | 12 | chr3:108,401,686 | chr3:108,399,584 | 2,102 | |  |  |  |
| 4 | fus(3).2 | 10 | chr3:105,571,481 | chr3:105,573,426 | 1,945 | | 4,179 | 0 | Inversion |
|  |  | 11 | chr3:105,792,873 | chr3:105,790,639 | 2,234 | |  |  |  |
| 5 | fus(3).3 | 10 | chr3:105,571,481 | chr3:105,573,426 | 1,945 | | 4,179 | 0 | Inversion |
|  |  | 11 | chr3:105,792,873 | chr3:105,790,639 | 2,234 | |  |  |  |
| 6 | fus(3).4 | 9 | chr3:103,861,058 | chr3:103,859,837 | 1,221 | | 5,231 | 5,300 | Duplication |
|  |  | 14 | chr3:109,242,307 | chr3:109,246,317 | 4,010 | |  |  |  |
| 7 | t(3;15).1 | 13 | NF | chr3:108,413,407 | - | | - | - | Acrocentric  short arm |
|  |  | 37 | NF | chr15p | - | |  |  |  |
| 8 | fus(3).5 | 22 | chr3:158,109,424 | chr3:158,124,238 | 14,814 | | 15,200 | 15,220 | Inversion |
|  |  | 20 | chr3:157,835,535 | chr3:157,835,921 | 386 | |  |  |  |
| 9 | fus(3).6 | 21 | chr3:158,130,903 | chr3:158,124,237 | 6,666 | | 14,880 | 14,783 |  |
|  |  | 19 | chr3:157,844,134 | chr3:157,835,920 | 8,214 | |  |  |  |
| 10 | t(3;15).2 | 18 | chr3:156,982,961 | chr3:156,977,787 | 5,174 | | 8,268 | 8,592 | No  (chimeric map) |
|  |  | 40 | chr15:31,645,858 | chr15:31,648,952 | 3,094 | |  |  |  |
| 11 | t(3;15).3 | 39 | chr15:30,749,354 | chr15:30,749,459 | 105 | | 282 | 8,029 | No  (chimeric map) |
|  |  | 17 | chr3:154,356,455 | chr3:154,356,278 | 177 | |  |  |  |
| 12 | fus(3).7 | 18 | chr3:156,976,622 | chr3:156,977,787 | 1,165 | | 4,905 | 3,653 | No  (chimeric map) |
|  |  | 33 | NF | chr11:118,932,550 | - | |  |  |  |
|  |  | 32 | NF | chr11:118,932,353 | - | |  |  |  |
|  |  | 5 | chr3:33,896,309 | chr3:33,892,569 | 3,740 | |  |  |  |
| **#JP** | **JP ID** | **#BP** | **BP OGM** | **BP lrGS** | **BPs difference (bp)** | | **Sum of BPs difference (bp)** | **Size of Uncertain OGM region (bp)** | **OGM call** |
| 13 | t(3;9).2 | 6 | chr3:78,517,154 | chr3:78,523,988 | 6,834 | | 8,905 | 9,143 | Yes |
|  |  | 25 | chr9:33,829,404 | chr9:33,831,475 | 2,071 | |  |  |  |
| 14 | fus(9) | **23** | chr9:20,877,178 | chr9:20,873,864 | 3,314 | | 19,939 | 19,975 | Yes, inversion |
|  |  | **26** | chr9:33,852,709 | chr9:33,836,084 | 16,625 | |  |  |  |
| 15 | t(3;10).1 | **27** | chr10:129,509,966 | chr10:129,525,155 | 15,189 | | 20,155 | 21,355 | Yes |
|  |  | **3** | chr3:30,422,251 | chr3:30,427,217 | 4,966 | |  |  |  |
| 16 | t(3;10).2 | 2 | chr3:29,915,529 | chr3:29,910,078 | 5,451 | | 20,185 | 20,304 | Yes |
|  |  | 27 | chr10:129,539,889 | chr10:129,525,155 | 14,734 | |  |  |  |
| 17 | fus(11).1 | 28 | chr11:115,756,201 | chr11:115,767,429 | 11,228 | | 16,051 | 18,046 | Deletion |
|  |  | 34 | chr11: 118,962,240 | chr11:118,957,417 | 4,823 | |  |  |  |
| 18 | fus(11).2 | 35 | chr11:119,275,141 | chr11:119,278,574 | 3,433 | | 5,928 | 8,121 | No |
|  |  | 30 | chr11:118,398,445 | chr11:118,400,940 | 2,495 | |  |  |  |
| 19 | t(11;15).1 | 29 | chr11:118,019,565 | chr11:118,006,938 | 12,627 | | 25,872 | 25,785 | Yes |
|  |  | 42 | chr15:53,619,488 | chr15:53,606,243 | 13,245 | |  |  |  |
| 20 | t(1;11).2 | 1 | chr1:175,249,885 | chr1:175,239,230 | 10,655 | | 35,882 | 38,115 | Yes |
|  |  | 29 | chr11:118,426,165 | chr11:118,400,938 | 25,227 | |  |  |  |
| 21 | fus(11).2 | 31 | NF | chr11:118,930,313 | - | | - | - | No |
|  |  | 34 | NF | chr11:118,957,417 | - | |  |  |  |
| 22 | t(11;15).2 | 33 | chr11:118,925,056 | chr11:118,932,550 | 7,494 | | 10,127 | 32,.908 | Yes |
|  |  | 42 | chr15:53,603,610 | chr15:53,606,243 | 2,633 | |  |  |  |
| 23 | t(3;15).4 | 41 | chr15:31,656,294 | chr15:31,648,953 | 7,341 | | 8,603 | 8,556 | No  (chimeric map) |
|  |  | 17 | chr3:154,355,016 | chr3:154,356,278 | 1,262 | |  |  |  |
| 24 | fus(3).8 | 16 | chr3:114,313,199 | chr3:114,289,729 | 23,470 | | 30,730 | 30,679 | No |
|  |  | 15 | chr3:114,164,677 | chr3:114,157,417 | 7,260 | |  |  |  |
| 25 | fus(3).9 | 16 | chr3:114,285,606 | chr3:114,289,729 | 4,123 | | 24,048 | 23,970 | No |
|  |  | 15 | chr3:114,137,492 | chr3:114,157,417 | 19,925 | |  |  |  |
| 26 | t(3;15).5 | 14 | NF | chr3:109,246,317 | - | | - | - | Acrocentric  short arm |
|  |  | 37 | NF | chr15p | - | |  |  |  |
| **#JP** | **JP ID** | **#BP** | **BP OGM** | **BP lrGS** | **BPs difference (bp)** | | **Sum of BPs difference (bp)** | **Size of Uncertain OGM region (bp)** | **OGM call** |
| 27 | t(3;15).6 | 38 | chr15:30,747,192 | chr15:30,748,255 | 1,063 | | 8,716 | 8,730 | No |
|  |  | 4 | chr3:33,884,889 | chr3:33,892,542 | 7,653 | |  |  |  |
| 28 | fus(3).10 | 3 | chr3:30,428,590 | chr3:30,427,217 | 1,373 | | 2,330 | 2,611 | Deletion |
|  |  | 2 | chr3:29,909,121 | chr3:29,910,078 | 957 | |  |  |  |
|  | | | | **Average** | | 6,451 |  |  |  |
|  | | | | **Standard Deviation** | | 6,325 |  |  |  |

JP: junction point; JP ID: junction point identification; BP(s): breakpoint(s); NF: not found. All genomic coordinates are according to reference genome GRCh38/hg38.

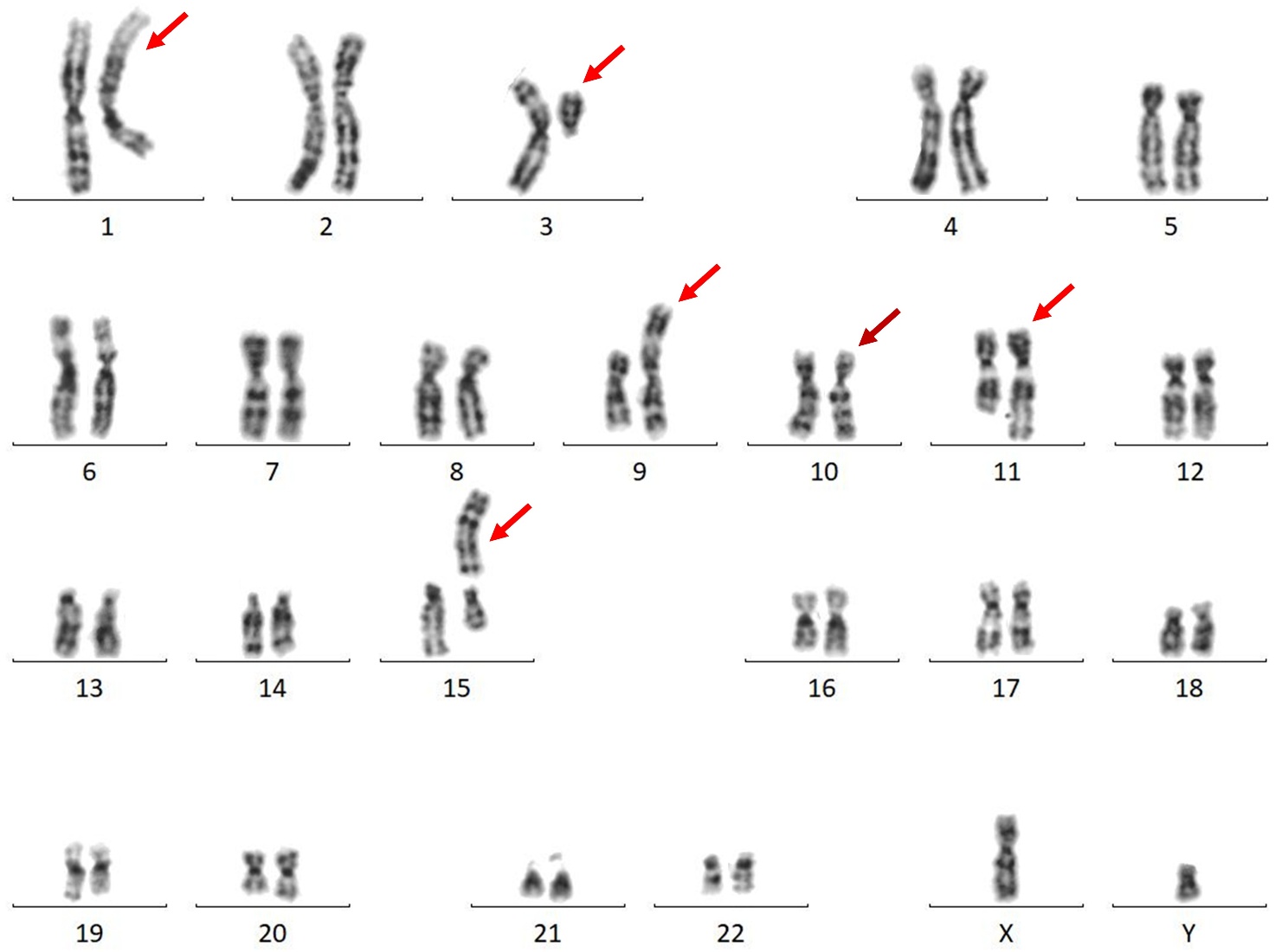

**Supplementary Figure 1. Karyotype of Patient 3.**

The red arrows highlight the five chromosomes involved in the patient’s rearrangement based on karyotyping and the burgundy arrow shows the sixth chromosome involved in the CCR, revealed by OGM and lrGS.

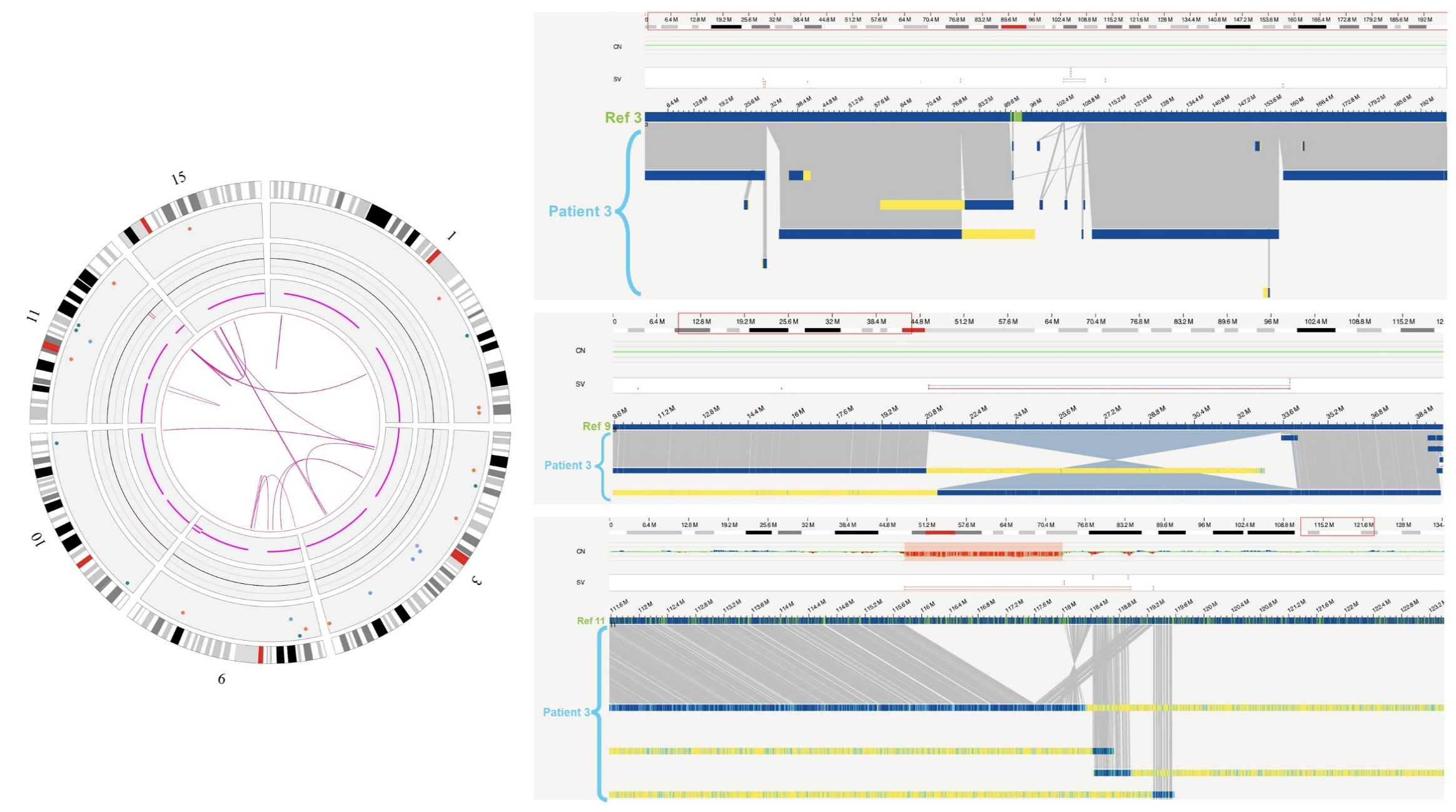

**Supplementary Figure 2. Optical Genome Mapping results of Patient 3.**

To the left, circos plot with OGM results of Patient 3 with the six involved chromosomes’ idiograms in the periphery and the pink lines in the middle representing intra- and inter-fusions. To the right, genome browser view of chromosomes 3 (top), 9 (middle) and 11 (bottom). It is possible to see various patient maps mapping to different regions of the same chromosome as well as chrimeric maps, whose yellow parts map to a different chromosome.

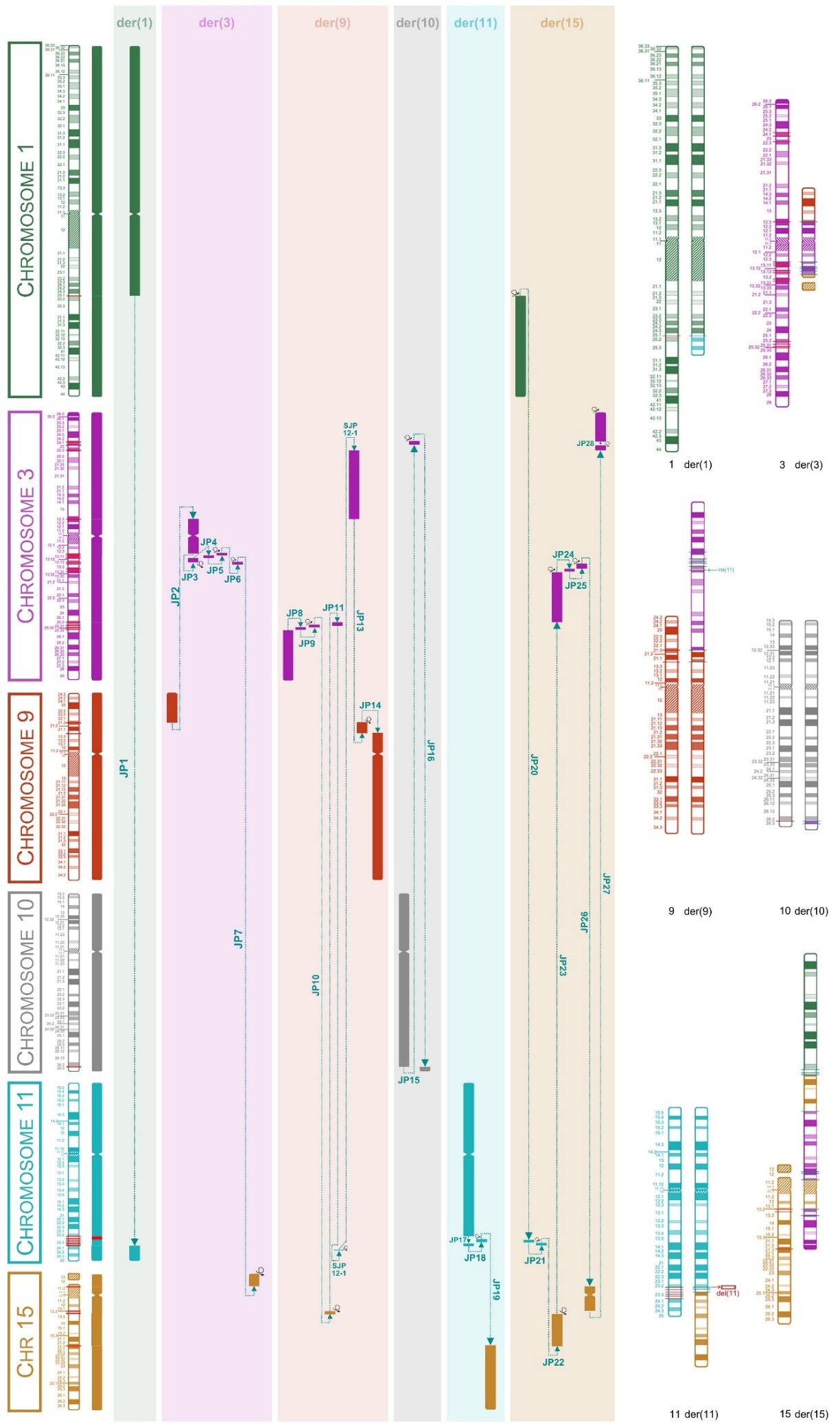

**Supplementary Figure 3. Patient 3’s resolved complex chromosomal rearrangement.**

To the left, a schematic representation of the junctions between all chromosome regions involved in the CCR. To the right, idiogram of the chromosomes involved in the complex chromosomal rearrangement. Red lines show the breakpoints in the normal chromosomes and cyan lines show the junction points in the derivative chromosomes.
