## Supplementary material for "Integrative optical genome mapping and long-read sequencing resolve constitutional complex rearrangements at nucleotide resolution": Supplementary File 4 - Patient 4.docx

Bruna Burssed, Bart van der Sanden, Wolfram Höps, Kornelia Neveling, Eveline Kamping, Ronald van Beek, Amber den Ouden, Ronny Derks, Raoul Timmermans, Eduardo Perrone, Marco Antonio Ramos, Fernanda Teixeira Bellucco, Alexander Hoischen, Maria Isabel Melaragno

**Supplementary File 4 – Additional information in Tables and Figures from Patient 4**

**TABLES.**

**FIGURES.**

**Supplementary Table 1. Patient 4’s phenotypes with Human Phenotype Ontology (HPO) terms.**

| **Phenotype** | **HPO** |
| --- | --- |
| **General** |  |
| Neurodevelopmental delay | HP:0012758 |
| Autism | HP:0000717 |
| Aggressive behavior | HP:0000718 |
| Lack of interest in peers | HP:4000083 |
| Agitation | HP:0000713 |
| **Central Nervous System** |  |
| Microcephaly | HP:0000252 |
| Brachycephaly | HP:0000248 |
| Cerebellar hypoplasia | HP:0001321 |
| Enlargement of the cisterna magna | HP:0002280 |
| **Genitourinary** |  |
| Cryptorchidism | HP:0000028 |
| **Ear** |  |
| Auditory hypersensitivity | HP:5200060 |
| **Facial dysmorphisms** |  |
| Thin eyebrows | HP:0045074 |
| Sparse eyebrows | HP:0045075 |
| Widow's peak | HP:0000349 |
| Slender nose | HP:0000417 |
| Long nasal bridge | HP:0033142 |
| Anteverted nostrils | HP:0000463 |
| Hypoplastic nostrils | HP:0000430 |

**Supplementary Table 2. Results of the techniques performed for the patient and final rearrangement.**

| **Technique** | **Result** |
| --- | --- |
| **Karyotype** | 46,XY |
| **Chromosomal Microarray Analysis** | arr[GRCh38] 2q37.2q37.3(236068385_239769318)×4 |
| **Optical Genome Mapping** | ogm[GRCh38] 2q37.2q37.3(236066554_239754226)amp |
| **Long-read Sequencing** | seq[GRCh38] 2q37.2q37.3(236052250_236062592×3,236062592_239777507×4,239777507_239778548×3) |
| **Final Rearrangement** | 46,XY,der(2)(2pter→2q37.3(chr2:239,778,549)::2q37.3(chr2:239,777,507)→2q37.2(chr2:236,062,592)::2q37.2(chr2:236,052,250)→2qter) |
|  | 46,XY,der(2)(2pter→2q37.3::2q37.3→2q37.2::2q37.2→2qter) |

**Supplementary Table 3. CNVs found in Patient 4.**

| **CNV** | **Start coordinate** | **End coordinate** | **Size (bp)** | **Copy Number** |
| --- | --- | --- | --- | --- |
| **chr2 duplication 1** | chr2:236,052,250 | chr2:236,062,591 | 10,342 | 3 |
| **chr2 triplication** | chr2:236,062,592 | chr2:239,777,507 | 3,714,915 | 4 |
| **chr2 duplication 2** | chr2:239,777,508 | chr2:239,778,549 | 1,041 | 3 |

CNV: copy number variant; chr: chromosome. All genomic coordinates are according to reference genome GRCh38/hg38.

**Supplementary Table 4. Breakpoint (BP) coordinates found through long-read sequencing and their number (#) for identification.**

| **#BP** | **BP coordinate** |
| --- | --- |
| 1 (a) | chr2:236,052,250 |
| 2 (b) | chr2:236,062,592 |
| 3 (c) | chr2:239,777,507 |
| 4 (d) | chr2:239,778,549 |

All genomic coordinates are according to reference genome GRCh38/hg38.

**Supplementary Table 5. Junction points and breakpoints details found on the long-read sequencing analysis and annotation of Repetitive Elements in regions surrounding the breakpoints.**

| **Junction Point** | | | | | **Breakpoint** | | | **Repetitive Elements** | | | | **Obs.** |
| --- | --- | --- | --- | --- | --- | --- | --- | --- | --- | --- | --- | --- |
| **#JP** | **JP ID** | **MH** | **Ins** | **Del** | **#BP** | **BP coord.** | **BP - 1 kb** | | **BP** | **BP + 1 kb** |  | |
| 1 | d/c | 1 (C) | - | - | 4 (d) | chr2:239,778,549 | SINE (MIR) – MIR  chr2:239778207-239778377 (-) | | - | - | - | |
|  |  |  |  |  |  |  | SINE (MIR) – MIR  chr2:239777751-239777846 (+) | |  |  |  |  |
|  |  |  |  |  | 3 (c) | chr2:239,777,507 | LINE (L1) - HAL1ME  chr2:239776684-239777009 (-) | | - | SINE (MIR) – MIR  chr2:239778207-239778377 (-) |  |  |
|  |  |  |  |  |  |  |  |  |  | SINE (MIR) – MIR  chr2:239777751-239777846 (+) |  |  |
| 2 | b/a | - | 29 | - | 2 (b) | chr2:236,062,592 | LINE (L1) - L1MB7  chr2:236061090-236061874 (+) | | SINE (Alu) – AluJr  chr2:236062465-236062642 (-) | SINE (Alu) – AluSg7  chr2:236062643-236062959 (-) | Insertion of 29 nt (TGCAGAAGTAGCTCCCAGCTACTTGGGAG) = 72% homeology with BP3 | |
|  |  |  |  |  |  |  |  |  |  | SINE (Alu) – AluJr  chr2:236062960-236063086 (-) |  |  |
|  |  |  |  |  |  |  |  |  |  | SINE (Alu) – AluSx1  chr2:236063099-236063380 (-) |  |  |
|  |  |  |  |  |  |  |  |  |  | LINE (L2) – L2  chr2:236063405-236063490 (-) |  |  |
|  |  |  |  |  | 1 (a) | chr2:236,052,250 | - | | - | Simple Repeat – (TGAA)n  chr2:236053253-236053288 (+) |  |  |

JP: junction point; JP ID: junction point identification; Ins: insertion; MH: microhomology; Del: deletion; BP: breakpoint; coord.: coordinate; Obs.: observation. All genomic coordinates are according to reference genome GRCh38/hg38.

**Supplementary Table 6. Difference between breakpoints found by Optical Genome Mapping (OGM) and Long-read Sequencing (lrGS).**

| **BP** | **BP OGM** | **BP lrGS** | **BPs difference (bp)** |
| --- | --- | --- | --- |
| 1 (a) | NF | chr2:236,052,250 | - |
| 2 (b) | chr2:236,066,554 | chr2:236,062,592 | 3,962 |
| 3 (c) | chr2:239,754,226 | chr2:239,777,507 | 23,281 |
| 4 (d) | NF | chr2:239,778,549 | - |

BP(s): breakpoint(s); bp: base pair; NF: not found. All genomic coordinates are according to reference genome GRCh38/hg38.

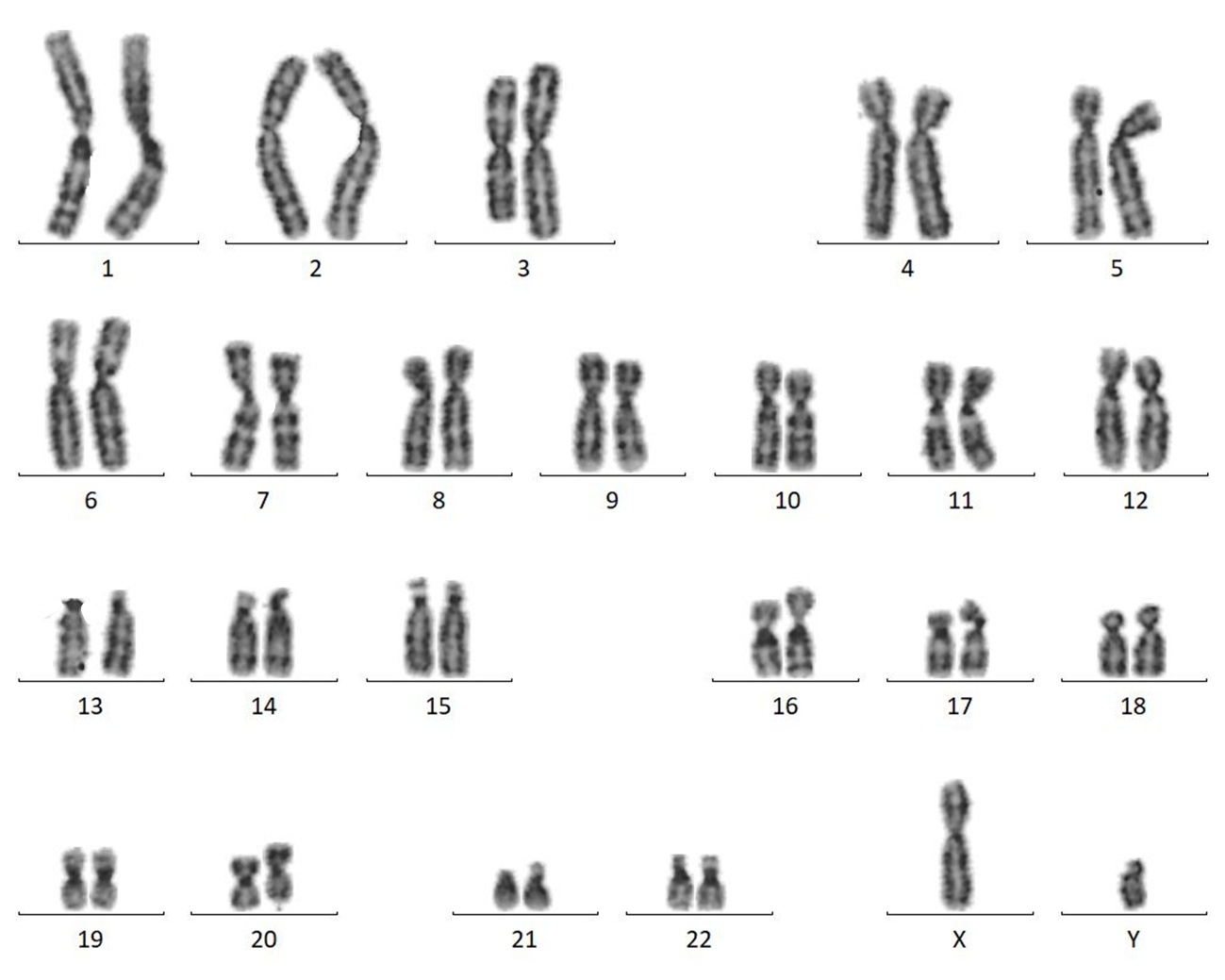

**Supplementary Figure 1. Karyotype of Patient 4.**

Initially, a normal karyotype (46,XY) was reported. After the resolution of the rearrangement with the other methodologies, a closer look at the karyotype allowed for the identification of a lighter chromosome band at the end of chromosome 2 that corresponds to the alteration (blue arrow).

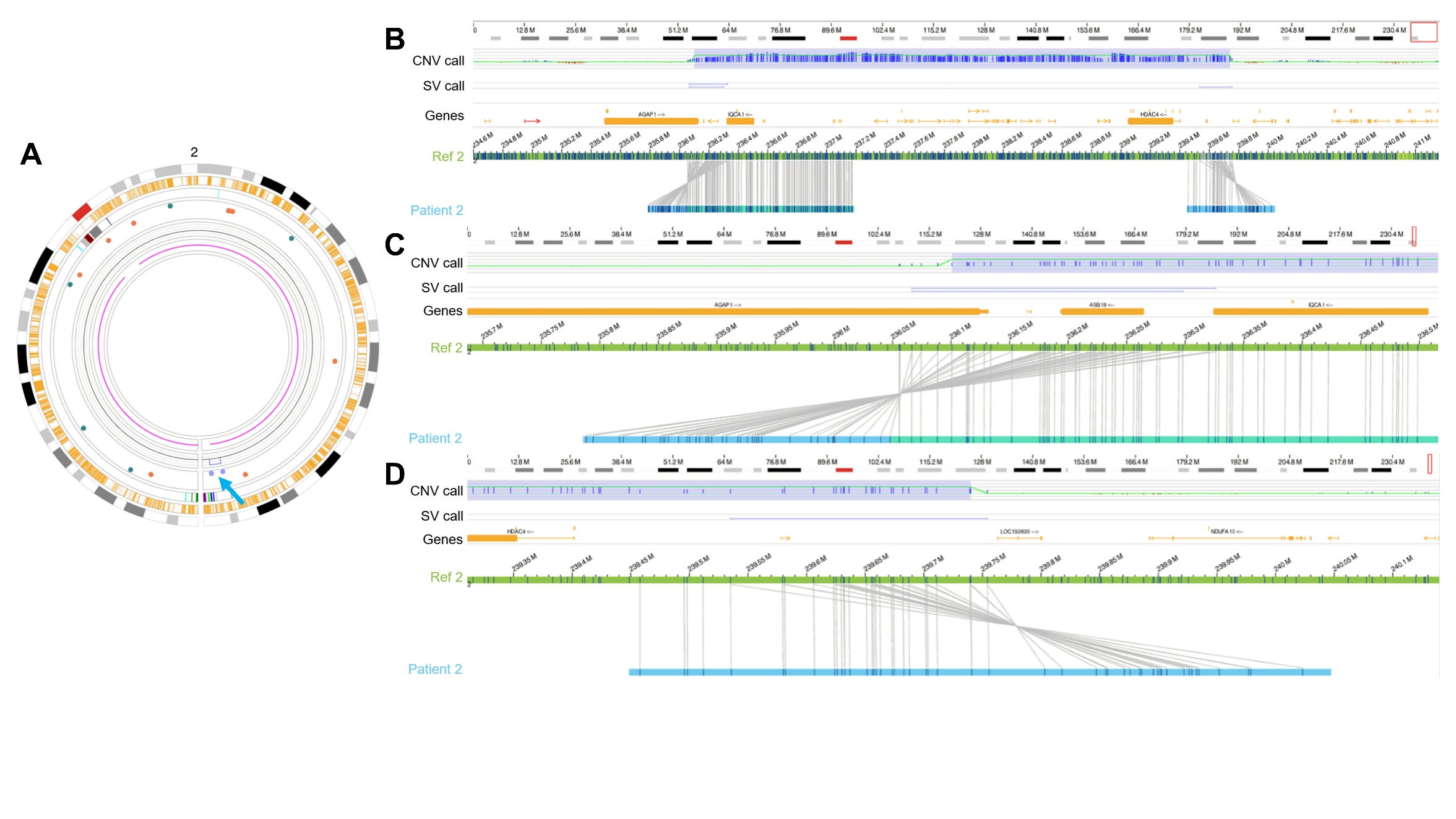

**Supplementary Figure 2. Optical Genome Mapping of Patient 4.**

**(A)** Circos plot showing chromosome 2. The chromosome idiogram can be seen in the periphery. The blue arrow draws attention to the patient’s copy number gains, shown by the lilac spheres (inverted duplications called by the SV pipeline) and the blue line (triplication called by the CNV pipeline). **(B) Chromosome 2 triplication.** From top to bottom, chromosome 2 idiogram, CNV call track, SV call track, Genes track, reference map from chromosome 2 (hg38) in green with labels in blue, patient map in light blue with labels in dark blue. The CNV call track shows a ~3.7 Mb triplication (blue square). The SV call track shows inverted duplications at the edges of the triplication called by the CNV pipeline (horizontal blue lines). **(C) Upstream chromosome 2 triplication breakpoint.** The patient map shows the inverted duplication called by the SV pipeline. **(D) Downstream chromosome 2 triplication breakpoint.** The patient map shows the inverted duplication called by the SV pipeline. Notice the large distance between the labels both in the reference and the patient maps.

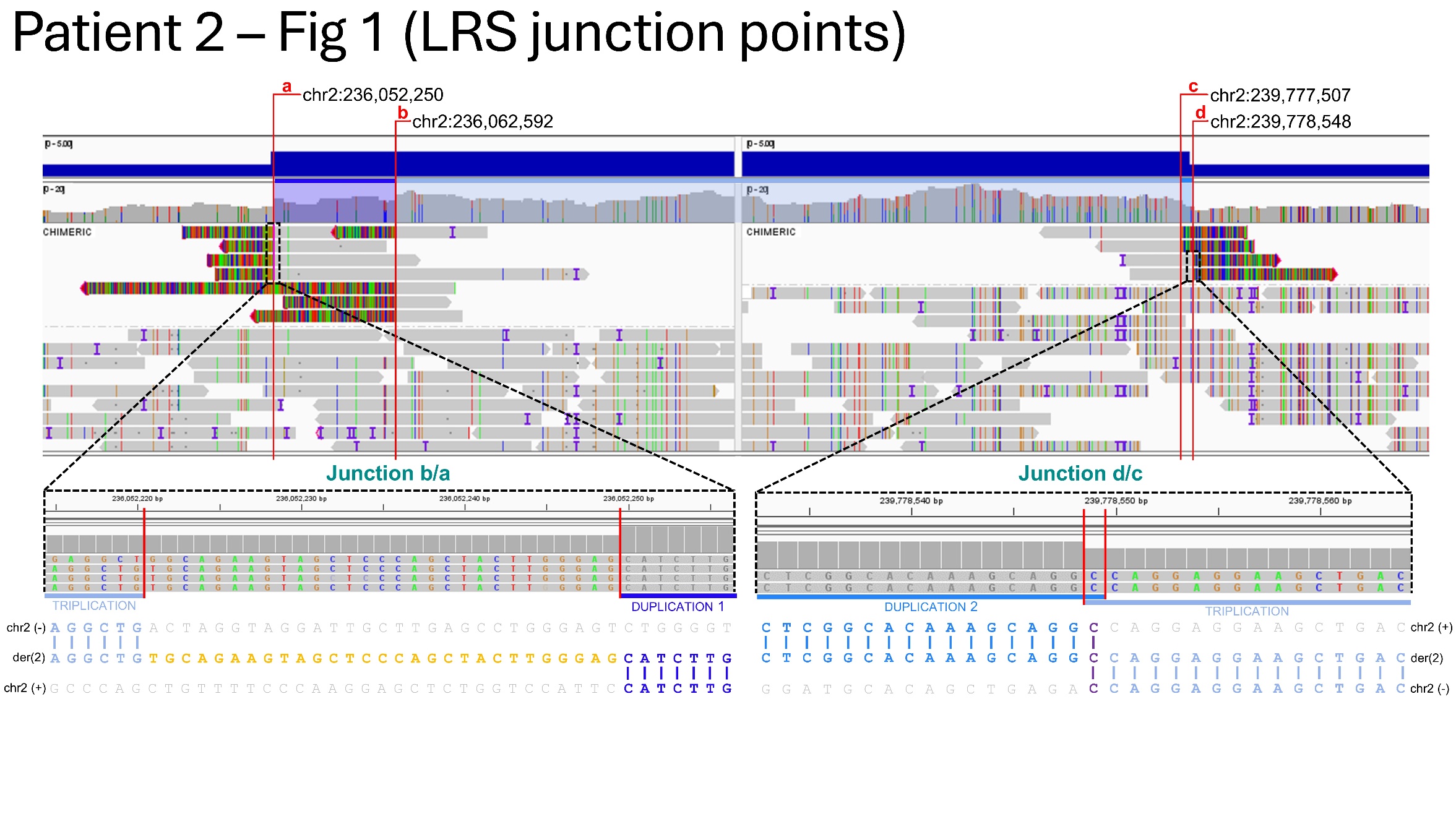

**Supplementary Figure 3. Patient 4’s Junction Points seen through long-read sequencing chimeric reads on IGV.**

At the top, IGV view of the chimeric reads in the regions involved in the CGR. The breakpoints (a, b, c, and d) are indicated with red lines. At the bottom, zoom in on the junction points at the nucleotide level with the sequence alignment and breakpoints below. In junction b/a, a 29-nucleotide insertion is revealed while in junction d/c, a 1-nucleotide microhomology is present.

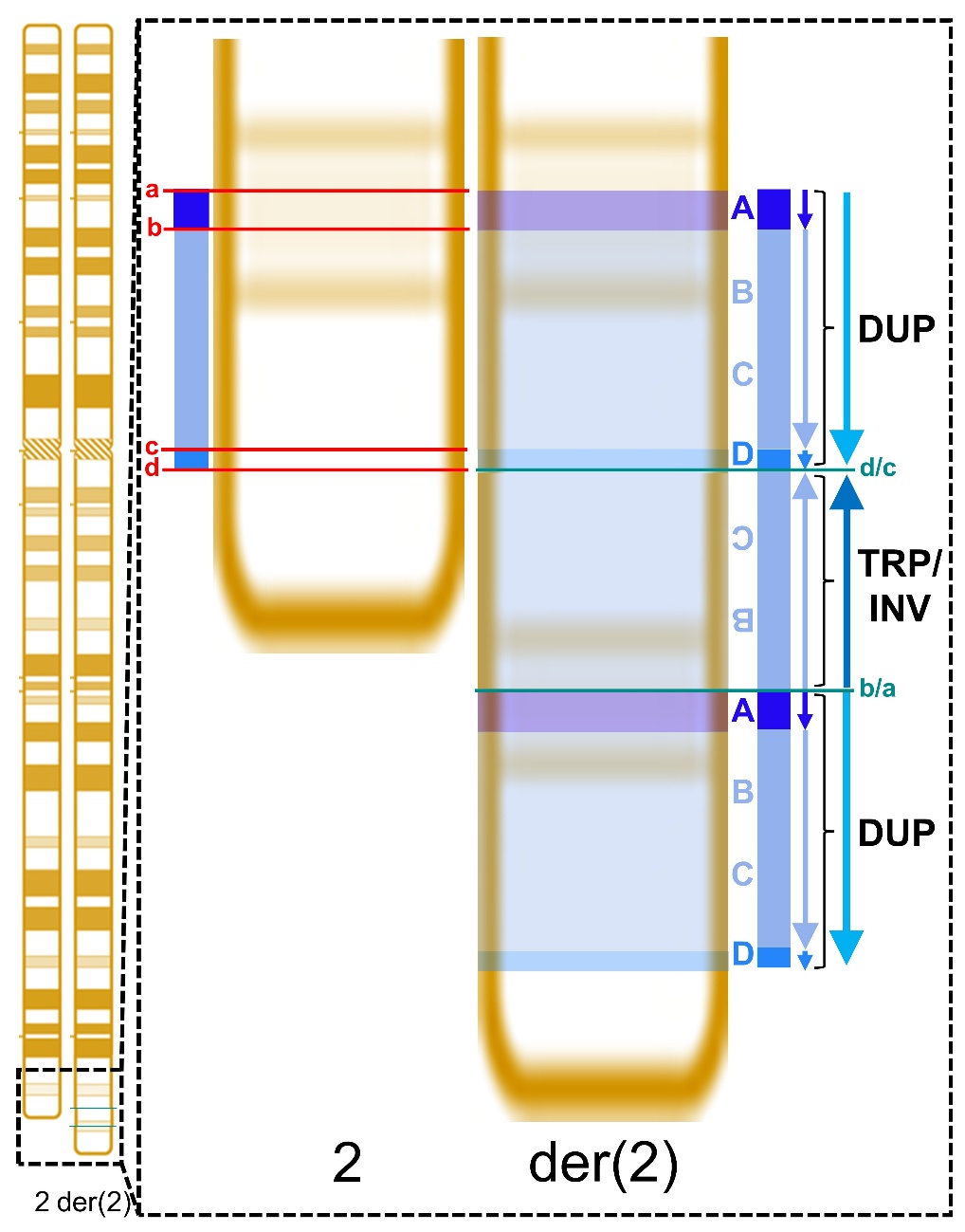

**Supplementary Figure 4. Final rearrangement of Patient 4’s rearranged chromosome 2.**

Idiogram of the chromosome involved in the complex genomic rearrangement. In the rectangle, to the left, a depiction of the breakpoints’ location in red (a, b, c, and d) and a representation of the first duplication between a and b, the triplication between b and c, and the second duplication between c and d. To the right, the characterization of the final DUP-TRP/INV-DUP rearrangement highlighting, in cyan lines, junction d/c between the second duplication and the inverted triplication and junction b/a between the inverted triplication and the first duplication.
