## Supplementary material for "Integrative optical genome mapping and long-read sequencing resolve constitutional complex rearrangements at nucleotide resolution": Supplementary File 5 - Patient 5.docx

Bruna Burssed, Bart van der Sanden, Wolfram Höps, Kornelia Neveling, Eveline Kamping, Ronald van Beek, Amber den Ouden, Ronny Derks, Raoul Timmermans, Eduardo Perrone, Marco Antonio Ramos, Fernanda Teixeira Bellucco, Alexander Hoischen, Maria Isabel Melaragno

**Supplementary File 5 – Additional information in Tables and Figures from Patient 5**

**TABLES.**

**FIGURES.**

**Supplementary Table 1. Patient 5’s phenotypes with Human Phenotype Ontology (HPO) terms**

| **Phenotype** | **HPO** |
| --- | --- |
| **General** |  |
| Neurodevelopmental delay | HP:0012758 |
| Seizure precipitated by febrile infection | HP:0032894 |
| Gastroenteritis-related afebrile seizure | HP:0032893 |
| **Brain/Head** |  |
| Frontal lobe hypoplasia | HP:0007333 |
| Cerebellar atrophy | HP:0001272 |
| Skull asymmetry | HP:0002678 |
| Posterior plagiocephaly | HP:0011327 |
| Blake's pouch cyst | HP:0033140 |
| **Eyes** |  |
| Abnormal retinal morphology | HP:0000479 |
| **Renal** |  |
| Renal insufficiency | HP:0000083 |
| Positive CMV urine nucleic acid test | HP:0034890 |
| **Ears** |  |
| Low-set ears | HP:0000369 |
| Small external ear | HP:0008772 |
| Prominent crus of helix | HP:0009899 |
| Auricular pit | HP:0030025 |
| **Hands/Feet** |  |
| Clinodactyly of the 5th finger | HP:0004209 |
| Fibular deviation of the 4th toe | HP:0100340 |
| Fibular deviation of the 3rd toe | HP:0100342 |
| Decreased palmar creases | HP:0006184 |
| **Facial dysmorphisms** |  |
| Facial hypotonia | HP:0000297 |
| Drooling (Sialorrhea) | HP:0002307 |
| Hypertelorism | HP:0000316 |
| Downslanted palpebral fissures | HP:0000494 |
| Depressed nasal bridge | HP:0005280 |
| Deep philtrum | HP:0002002 |
| Thin upper lip | HP:0000219 |

**Supplementary Table 2. Results of the techniques performed for the patient and final rearrangement.**

| **Technique** | **Result** |
| --- | --- |
| **Karyotype** | 46,XY,add(16)(q13) |
| **Chromosomal Microarray Analysis** | arr[GRCh38] 16p13.3(7097013_7610932)×1, 16q11.2q12.1(46455602_51233521)×3, 16q12.1q21(chr16:52507011_59255682)×3, 16q21q23.1(chr16:64697095_76450931)×3 |
| **Optical Genome Mapping** | ogm[GRCh38] 16p13.3(7083445_7611022)×1, 16q11.2q12.1(46404936_51222712)×3, 16q12.1q21(52526194_59290844)×3, 16q21q23.1(64690708_76452718)×3, 16q23.1(78530249_78575587)×1 |
| **Long-read Sequencing** | seq[GRCh38] 16p13.3(7098043_7606629)×1, 16q11.2q12.1(46381025_51230394)×3, 16q12.1q21(52516986_59290869)×3, 16q21q23.1(64686006_76454258)×3, 16q23.1(78534493_78567573)×1 |
| **Final Rearrangement** | 46,XY,der(16)(16pter→16p13.3(chr16:7,098,043)::16p13.3(chr16:7,606,629)→16q12.1(chr16:51,230,394)::16q11.2  (chr16:46,381,025)→16q21(chr16:59,290,869)::16q12.1(chr16:52,516,986)→16q23.1(chr16:76,454,258)::16q23.2  (chr16:80,357,530)→16q23.2(chr16:80,357,803)::16q21(chr16:64,686,006)→16q23.1(chr16:78,534,493)::16q23.2  (chr16:80,365,240)→16q23.2(chr16:80,365,335)::16q23.1(chr16:78,567,573)→16qter) |
|  | 46,XY,der(16)(16pter→16p13.3::16p13.3→16q12.1::16q11.2→16q21::16q12.1→16q23.1::16q23.2→  16q23.2::16q21→16q23.1::16q23.2→16q23.2::16q23.1→16qter) |

**Supplementary Table 3. Breakpoint (BP) coordinates found through long-read sequencing and their number (#) for identification.**

| **#BP** | **BP coordinate** |
| --- | --- |
| 1 | chr16:7,098,043 |
| 2 | chr16:7,606,629 |
| 3 | chr16:46,381,025 |
| 4 | chr16:51,230,394 |
| 5 | chr16:52,516,986 |
| 6 | chr16:59,290,869 |
| 7 | chr16:64,686,006 |
| 8 | chr16:76,454,258 |
| 9 | chr16:78,534,493 |
| 10 | chr16:78,567,573 |
| 11 | chr16:80,357,530 |
| 12 | chr16:80,357,803 |
| 13 | chr16:80,365,240 |
| 14 | chr16:80,365,335 |

**Supplementary Table 4. Junction points and breakpoints details found on the long-read sequencing analysis and annotation of Repetitive Elements in regions surrounding the breakpoints.**

|  | **Junction Point** | | | | | | **Breakpoint** | | **Repetitive Elements** | | |
| --- | --- | --- | --- | --- | --- | --- | --- | --- | --- | --- | --- |
| **#JP** | **JP ID** | **Ins** | **MH** | **Subjunction** | | |  |  |  |  |  |
|  |  |  |  | **#SJP** | **Ins** | **MH** | **#BP** | **BP coord.** | **BP - 1 kb** | **BP** | **BP + 1 kb** |
| **1** | del16p | 1 (A) | - | - | - | - | 1 | chr16:7,098,043 | LTR (ERVL-MaLR) – MLT1K  chr16:7096953-7097056 (-) | - | SINE (Alu) – AluSz  chr16:7098163-7098467 (-) |
|  |  |  |  |  |  |  |  |  |  |  | SINE (Alu) – AluJo  chr16:7098670-7098785 (+) |
|  |  |  |  |  |  |  |  |  |  |  | Simple Repeat – (AAAT)n  chr16:7098792-7098817 (+) |
|  |  |  |  |  |  |  |  |  |  |  | SINE (MIR) – MIR  chr16:7098882-7098951 (+) |
|  |  |  |  |  |  |  | 2 | chr16:7,606,629 | SINE (Alu) – AluSx3  chr16:7606119-7606420 (-) | - | - |
|  |  |  |  |  |  |  |  |  | SINE (Alu) – AluSz6  chr16:7605842-7606110 (-) |  |  |
| **2** | dup1 | - | - | - | - | - | 4 | chr16:51,230,394 | DNA (TcMar-Tigger) – MER6A  chr16:51229587-51229866 (+) | LINE (L1) – L1PA8  chr16:51229905-51231292 (+) | DNA (TcMar-Tigger) – MER6A  chr16: 51231312-51231817 (+) |
|  |  |  |  |  |  |  |  |  | SINE (MIR) – MIRb  chr16:51229432-51229534 (+) |  |  |
|  |  |  |  |  |  |  |  |  | LTR (ERVL-MaLR) – MLT1O  chr16:51229212-51229427 (-) |  |  |
|  |  |  |  |  |  |  | 3 | chr16:46,381,025 | Human Satellite II – Pericentromeric region | | |
| **3** | dup2 | 1 (A) | - | - | - | - | 6 | chr16:59,290,869 | SINE (MIR) – MIRb  chr16:59290250-59290291 (+) | - | LTR (Gypsy) – MamGypLTR3  chr16:59291059-59291303 (+) |
|  |  |  |  |  |  |  |  |  | SINE (Alu) – AluYa5  chr16:59289829-59290133 (-) |  | SINE (Alu) – AluJr  chr16:59291591-59291712 (-) |
|  |  |  |  |  |  |  | 5 | chr16:52,516,986 | LINE (L1) – L1ME1  chr16:52516730-52516902 (+) | - | - |
|  |  |  |  |  |  |  |  |  | SINE (MIR) – MIR3  chr16:52516047-52516104 (-) |  |  |
| **4** | dup3 | 277  (3.0 Mb from BP7) | - | 4-1 | - | 3 (AG  G) | 8 | chr16:76,454,258 | - | SINE (Alu) – AluSp  chr16:80357314-80357620 (+) | - |
|  |  |  |  |  |  |  | 11 | chr16:80,357,530 | - | SINE (Alu) – AluSz6  chr16:76454104-76454329 (-) | - |
|  |  |  | - | 4-2 | - | 3 (A  T  T) | 12 | chr16:80,357,803 |  | SINE (MIR) – MIRb  chr16:80357769-80357859 (-) | - |
|  |  |  |  |  |  |  | 7 | chr16:64,686,006 | LTR (ERVL-MaLR) – MLT1G1  chr16:64685816-64685908 (+) | - | Simple Repeat – (TATT)n  chr16:64686104-64686152 (+) |
|  |  |  |  |  |  |  |  |  | LINE (L2) – L2a  chr16:64684559-64685084 (+) |  | LINE (L2) – L2  chr16:64686287-64686531 (+) |
| **5** | del16q | 96  (1.8 Mb from BP10) | 1 (G) | 5-1 | - | - | 9 | chr16:78,534,493 | SINE (MIR) – MIRb  chr16:78534168-78534348 (+) | - | SINE (Alu) – AluSx3  chr16:78534701-78535010 (-) |
|  |  |  |  |  |  |  |  |  | LINE (L2) – L2b  chr16:78533666-78533952 (+) |  |  |
|  |  |  |  |  |  |  |  |  | DNA (hAT-Charlie) – MER30  chr16:78533322-78533535 (-) |  |  |
|  |  |  |  |  |  |  | 13 | chr16:80,365,240 | - | SINE (MIR) – MIRb  chr16:80365154-80365284 (-) | - |
|  |  |  | - | 5-2 |  | 1 (A) | 14 | chr16:80,365,335 | - | - | - |
|  |  |  |  |  |  |  | 10 | chr16:78,567,573 | SINE (MIR) – MIRb  chr16:78566985-78567127 (-) | SINE (Alu) – AluY  chr16:78567264-78567578 (+) | SINE (Alu) – AluSx1  chr16:78568475-78568774 (-) |
|  |  |  |  |  |  |  |  |  | Simple Repeat – (AATAGT)n  chr16:78566904-78566948 (+) |  |  |
| **#JP** | **JP ID** | **Ins** | **MH** | **#SJP** | **MH** | **Del** | **#BP** | **BP coord.** | **BP - 1 kb** | **BP** | **BP + 1 kb** |
|  |  |  |  | **Subjunction** | | | **Breakpoint** | | **Repetitive Elements** | | |
| **Junction Point** | | | | | | |  |  |  |  |  |

JP: junction point; JP ID: junction point identification; Ins: insertion; SJP: sub-junction point; MH: microhomology; Del: deletion; BP: breakpoint; coord.: coordinate; Obs.: observation. All genomic coordinates are according to reference genome GRCh38/hg38.

**Supplementary Table 5. Comparison of breakpoints and CNV’s location and size between reference genomes GRCh38/hg38 and T2T-CHM13.**

|  | **GRCh38/hg38** | | | **T2T-CHM13** | | | **Obs.** |
| --- | --- | --- | --- | --- | --- | --- | --- |
| **JP ID** | **BP1** | **BP2** | **Size (bp)** | **BP1** | **BP2** | **Size (bp)** |  |
| **del16p** | chr16:7,098,043 | chr16:7,606,629 | 508,586 | chr16:7,128,212 | chr16:7,638,100 | 509,888 | Coordinates match perfectly according to LiftOver despite the difference in sizes |
| **dup1** | chr16:46,381,025 | chr16:51,230,394 | 4,849,370 | chr16:51,745,471 | chr16:57,028,284 | 5,282,813 |  |
| **dup2** | chr16:52,516,986 | chr16:59,290,869 | 6,773,883 | chr16:58,314,876 | chr16:65,082,941 | 6,768,065 |  |
| **dup3** | chr16:64,686,006 | chr16:76,454,258 | 11,768,252 | chr16:70,474,539 | chr16:82,510,471 | 12,035,932 |  |
| **del16q** | chr16:78,534,493 | chr16:78,567,573 | 33,080 | chr16:84,590,857 | chr16:84,623,924 | 33,067 |  |
| **ins JP4** | chr16:80,357,530 | chr16:80,357,803 | 273 | chr16:86,418,936 | chr16:86,419,209 | 273 | Extra copy inserted in JP4 |
| **ins JP5** | chr16:80,365,240 | chr16:80,365,335 | 95 | chr16:86,426,646 | chr16:86,426,741 | 95 | Extra copy inserted in JP5 |

JP ID: junction point identification; BP: breakpoint; Obs.: observation.

**Supplementary Table 6. Difference between breakpoints found by Optical Genome Mapping (OGM) and Long-read Sequencing (lrGS).**

| **#JP** | **JP ID** | **BP** | **BP OGM** | | **BP lrGS** | **BPs difference (bp)** | **Sum of BPs difference (bp)** | **Size of Uncertain OGM region (bp)** |
| --- | --- | --- | --- | --- | --- | --- | --- | --- |
| 1 | del16p | 1 | chr16:7,083,445 | | chr16:7,098,043 | 14,598 | 18,991 | 19,061 |
|  |  | 2 | chr16:7,611,022 | | chr16:7,606,629 | 4,393 |  |  |
| 2 | dup1 | 4 | chr16:51,222,712 | | chr16:51,230,394 | 7,682 | 31,593 | - |
|  |  | 3 | chr16:46,404,936 | | chr16:46,381,025 | 23,911 |  |  |
| 3 | dup2 | 6 | chr16:59,290,844 | | chr16:59,290,869 | 25 | 9,233 | 9,668 |
|  |  | 5 | chr16:52,526,194 | | chr16:52,516,986 | 9,208 |  |  |
| 4 | dup3 | 8 | chr16:76,452,718 | | chr16:76,454,258 | 1,540 | 6,242 | 6,502 |
|  |  | 11 | NF | | chr16:80,357,530 | - |  |  |
|  |  | 12 | NF | | chr16:80,357,803 | - |  |  |
|  |  | 7 | chr16:64,690,708 | | chr16:64,686,006 | 4,702 |  |  |
| 5 | del16q | 9 | chr16:78,530,249 | | chr16:78,534,493 | 4,244 | 12,258 | 13,156 |
|  |  | 13 | NF | | chr16:80,365,240 | - |  |  |
|  |  | 14 | NF | | chr16:80,365,335 | - |  |  |
|  |  | 10 | chr16:78,575,587 | | chr16:78,567,573 | 8,014 |  |  |
|  | | | | **Average** | | 7,832 |  |  |
|  | | | | **Standard Deviation** | | 7,002 |  |  |

JP: junction point; JP ID: junction point identification; BP(s): breakpoint(s); NF: not found. All genomic coordinates are according to reference genome GRCh38/hg38.

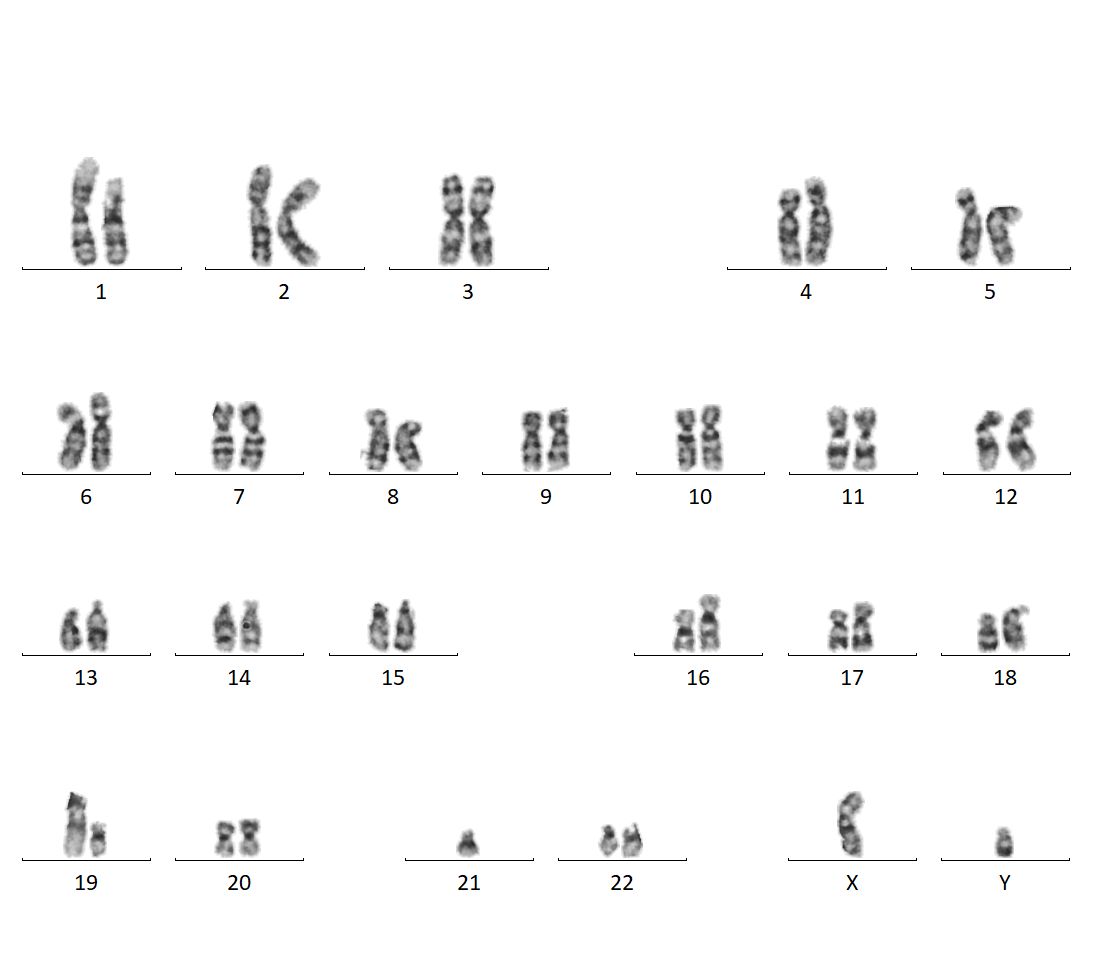

**Supplementary Figure 1. Karyotype of Patient 5.**

The red arrow highlights the chromosome involved in the patient’s rearrangement.

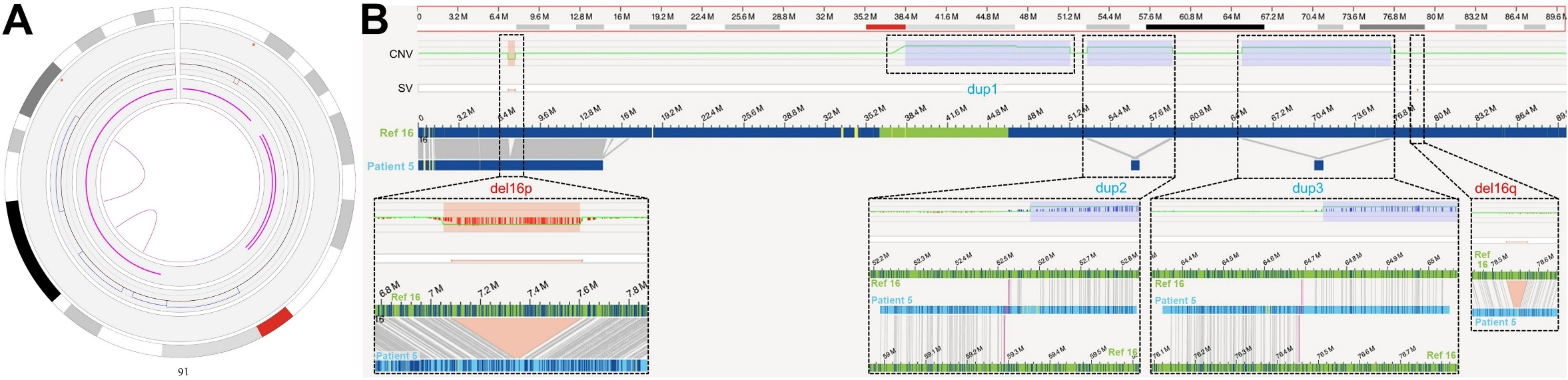

**Supplementary Figure 2. Optical Genome Mapping results of Patient 5.**

To the left, circos plot with OGM results of Patient 5’s chromosome 16 CGR with the chromosome’s idiogram in the periphery, CNVs in the following layer, and the pink lines in the middle representing inter-fusions. To the right, genome browser view of chromosome 16, highlight the CNVs whose position and orientation could be resolve with OGM.
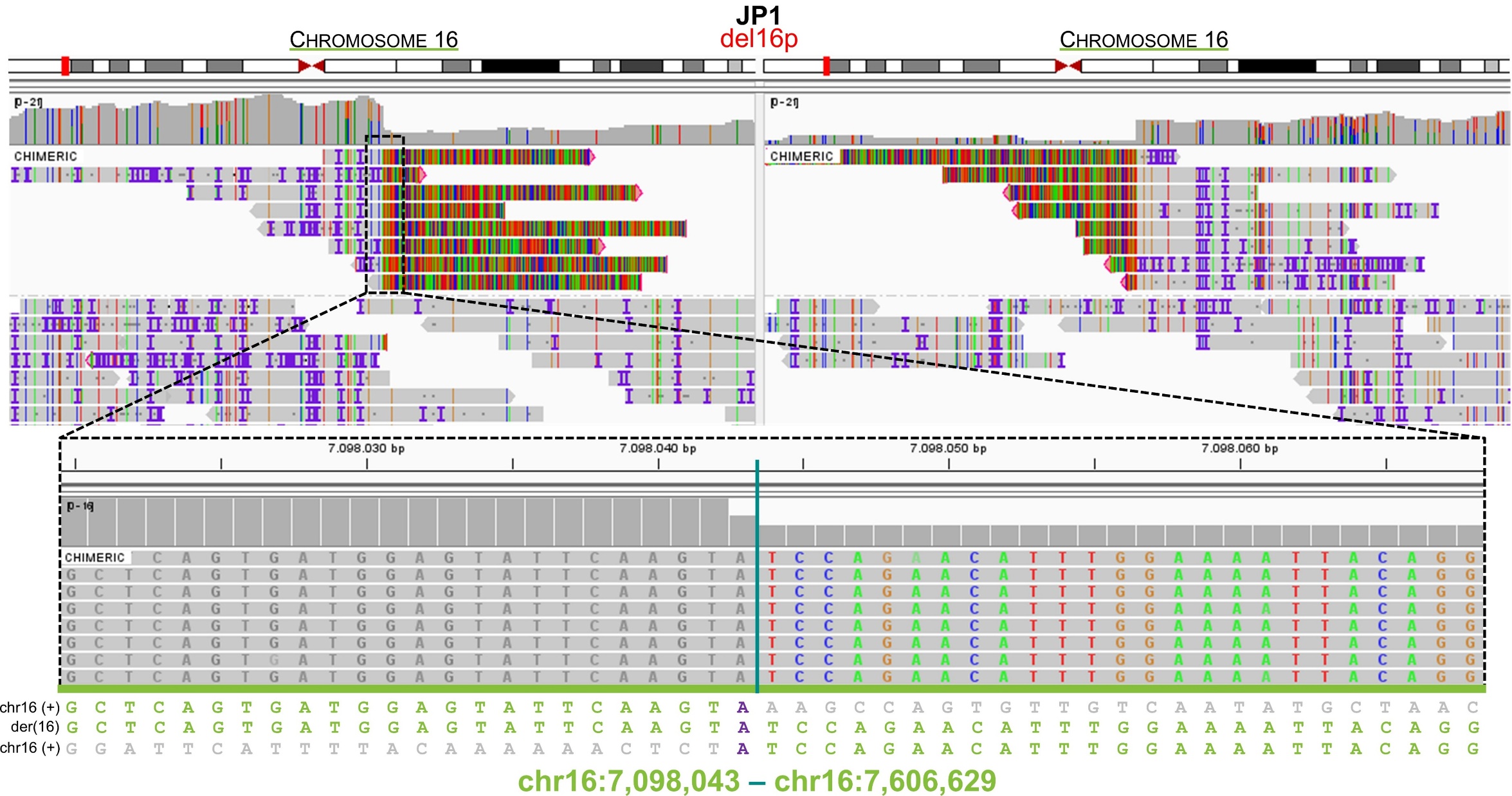

**Supplementary Figure 3. Patient 5’s Junction Point 1 (del16p) seen through long-read sequencing chimeric reads in IGV.**

At the top, IGV view of the chimeric reads in the regions involved in the CGR. At the bottom, zoom in on the junction point at the nucleotide level with the sequence alignment and breakpoints below.

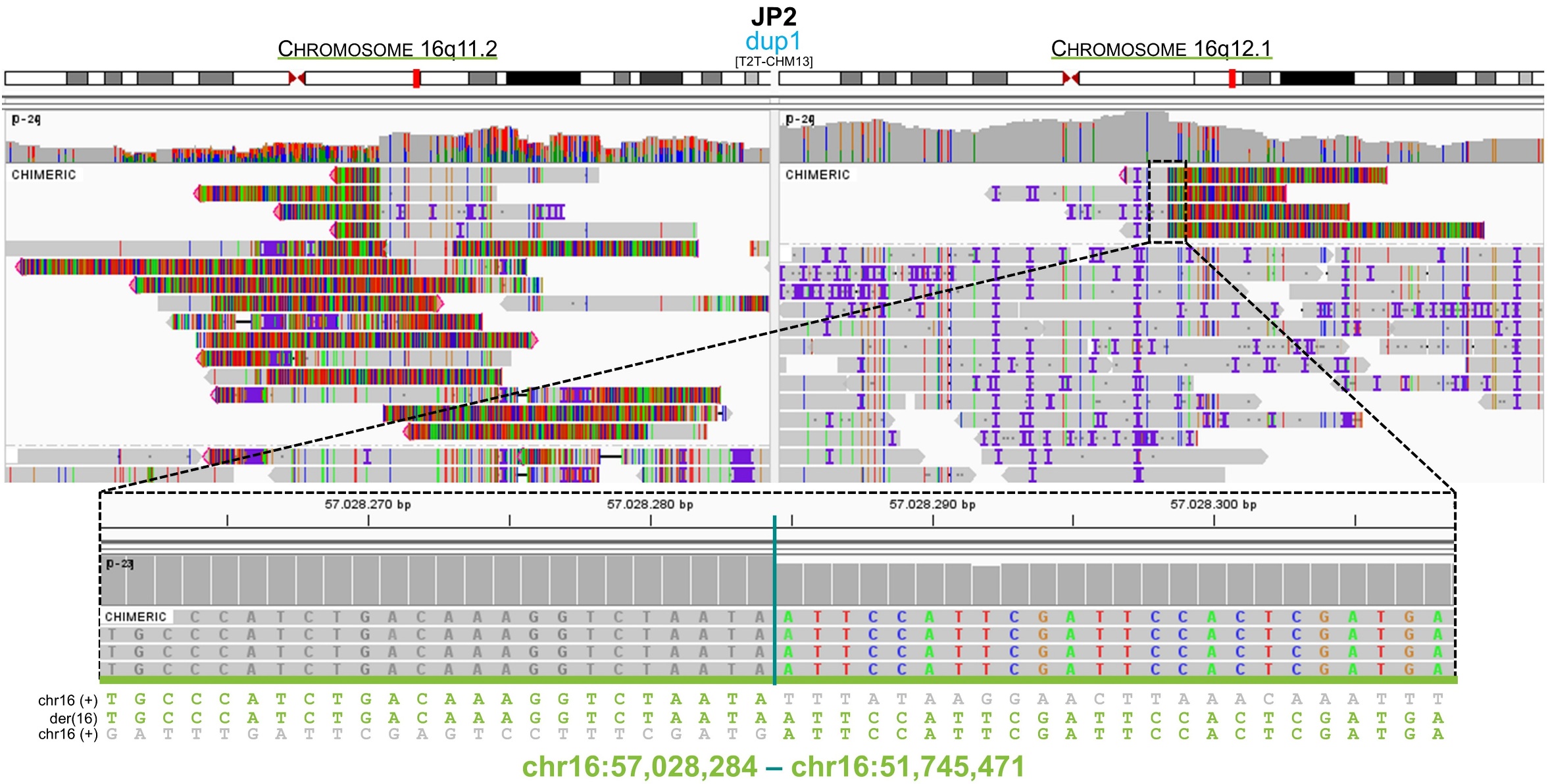

**Supplementary Figure 4. Patient 5’s Junction Point 2 (dup1) seen through long-read sequencing chimeric reads in IGV.**

At the top, IGV view of the chimeric reads in the regions involved in the CGR. At the bottom, zoom in on the junction point at the nucleotide level with the sequence alignment and breakpoints below.

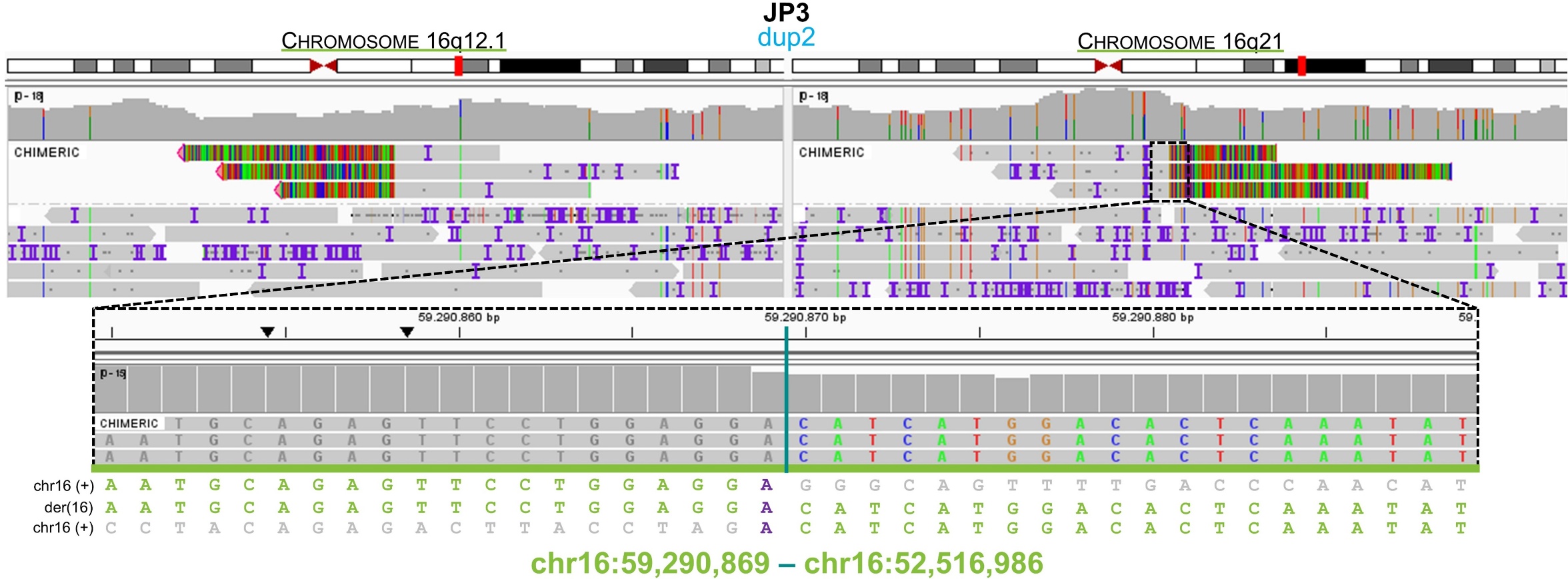

**Supplementary Figure 5. Patient 5’s Junction Point 3 (dup2) seen through long-read sequencing chimeric reads in IGV.**

At the top, IGV view of the chimeric reads in the regions involved in the CGR. At the bottom, zoom in on the junction point at the nucleotide level with the sequence alignment and breakpoints below.

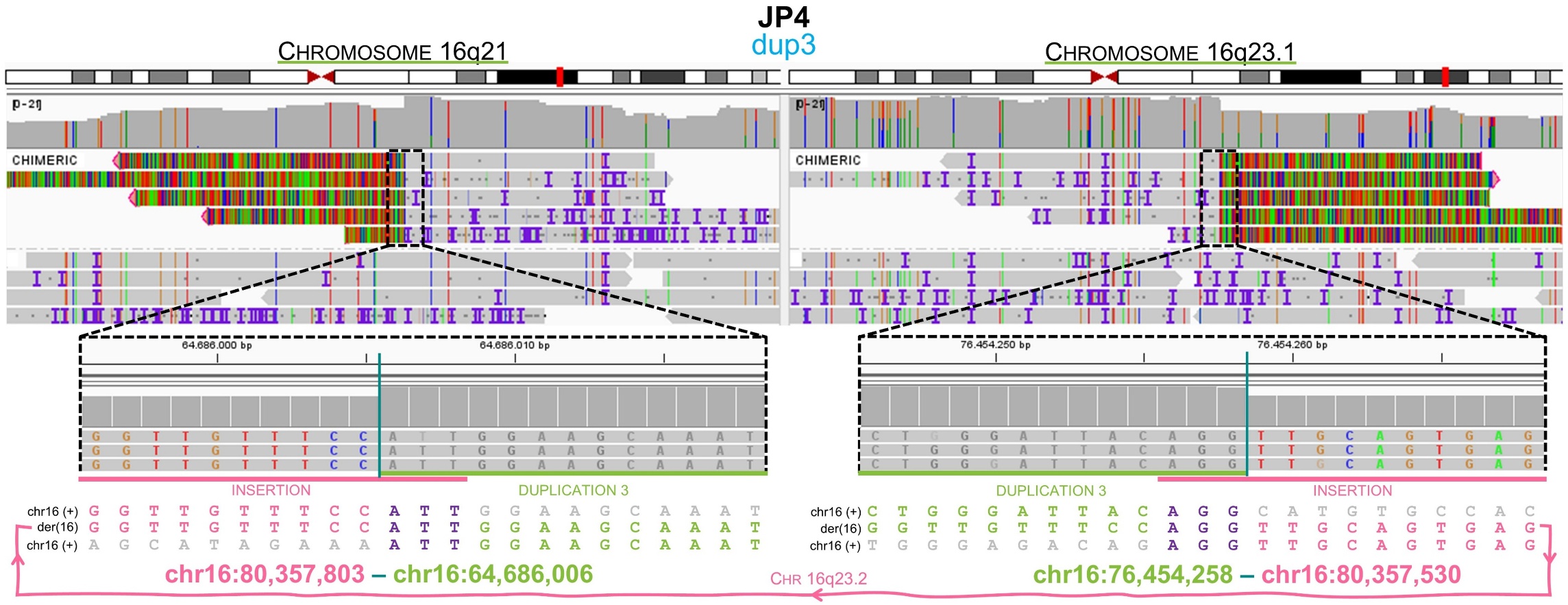

**Supplementary Figure 6. Patient 5’s Junction Point 4 (dup3) seen through long-read sequencing chimeric reads in IGV.**

At the top, IGV view of the chimeric reads in the regions involved in the CGR. At the bottom, zoom in on the junction point at the nucleotide level with the sequence alignment and breakpoints below.

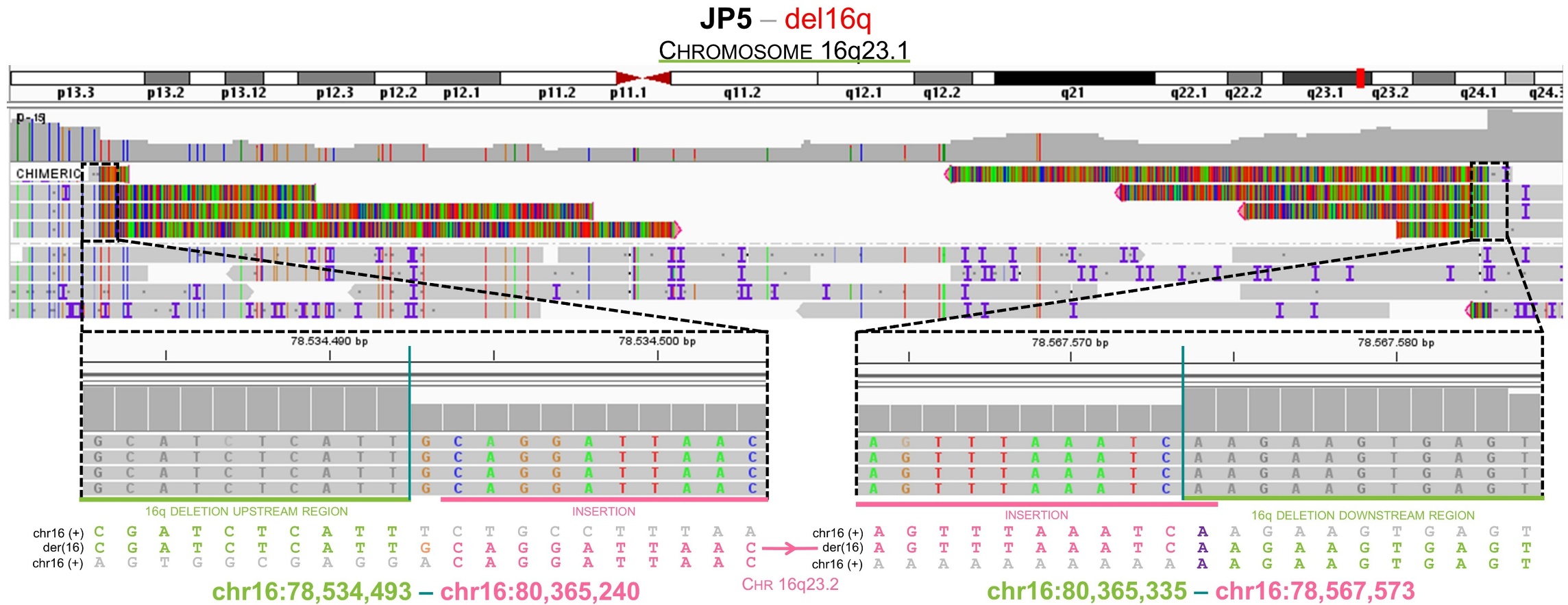

**Supplementary Figure 7. Patient 5’s Junction Point 5 (del16q) seen through long-read sequencing chimeric reads in IGV.**

At the top, IGV view of the chimeric reads in the regions involved in the CGR. At the bottom, zoom in on the junction point at the nucleotide level with the sequence alignment and breakpoints below.

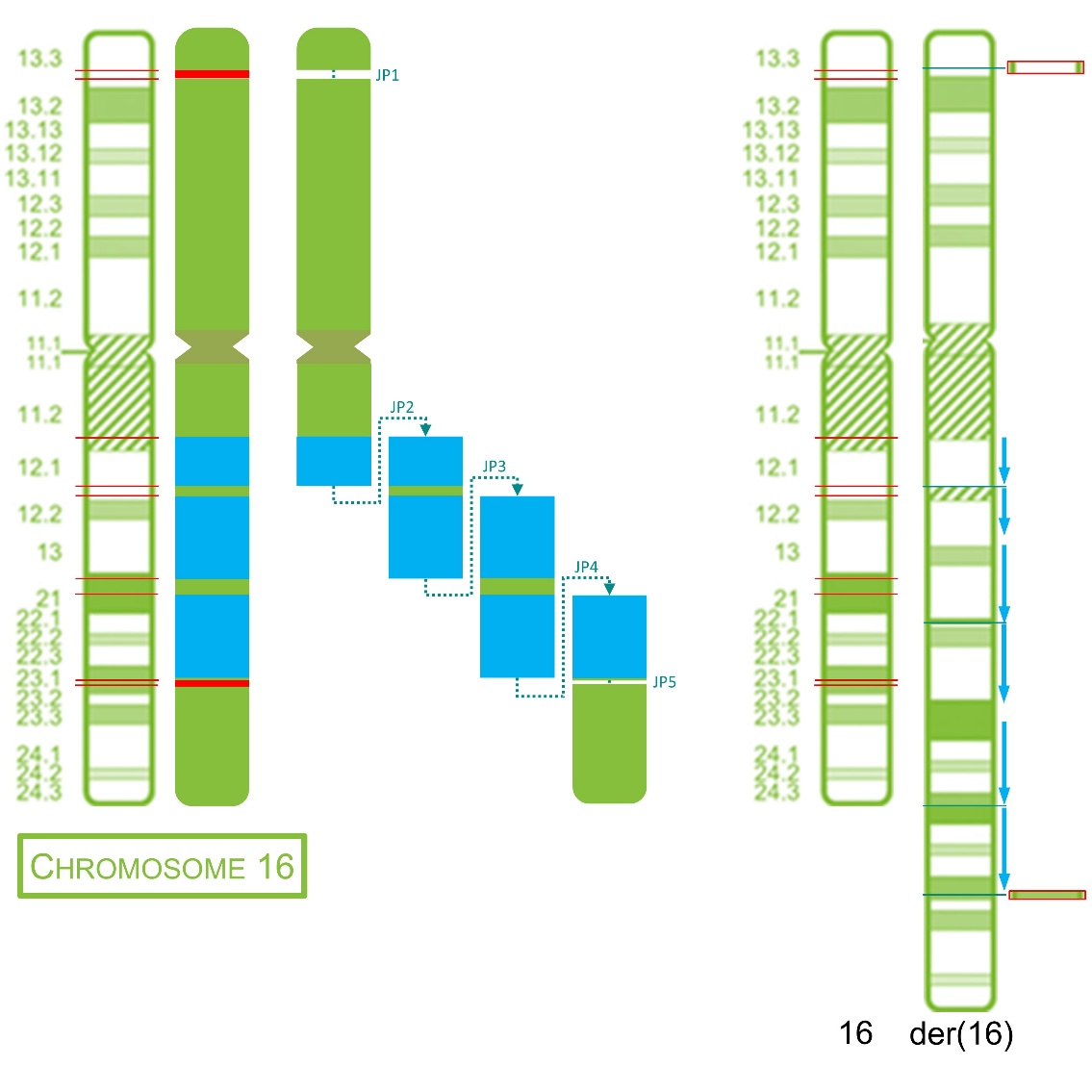

**Supplementary Figure 8. Final rearrangement of Patient 5’ rearranged chromosome 16.**

To the left, a schematic representation of the junctions between all chromosome 16 regions involved in the CGR. To the right, idiogram of the chromosome involved in the complex genomic rearrangement. Red lines show the breakpoints in the normal chromosomes and cyan lines show the junction points in the derivative chromosomes.
