## Supplementary material for "Integrative optical genome mapping and long-read sequencing resolve constitutional complex rearrangements at nucleotide resolution": Supplementary File 6 - Patient 6.docx

Bruna Burssed, Bart van der Sanden, Wolfram Höps, Kornelia Neveling, Eveline Kamping, Ronald van Beek, Amber den Ouden, Ronny Derks, Raoul Timmermans, Eduardo Perrone, Marco Antonio Ramos, Fernanda Teixeira Bellucco, Alexander Hoischen, Maria Isabel Melaragno

**Supplementary File 6 – Additional information in Tables and Figures from Patient 6**

**TABLES.**

**FIGURES.**

**Supplementary Table 1. Patient 6’s phenotypes with Human Phenotype Ontology (HPO) terms**

| **Phenotype** | **HPO** |
| --- | --- |
| **General** |  |
| Mild intellectual disability | HP:0001256 |
| Autism | HP:0000717 |
| Epilepsy | HP:0001250 |

**Supplementary Table 2. Results of the techniques performed for the patient and final rearrangement.**

| **Technique** | **Result** |
| --- | --- |
| **Karyotype** | 46,XY,add(18)(p11.2) |
| **Chromosomal Microarray Analysis** | arr[GRCh38] 18q21.1q21.32(50,255,939_58,918,298)×3, 18q22.2q23(69471830-76253294)×1 |
| **Optical Genome Mapping** | ogm[GRCh38] 18q11.2(26,832,573_27,032,179)×3, 18q12.1q12.2(32,037,307_35,583,505)×1, 18q21.1q21.2(50,215,914_58,857,454)×3, 18q22.2q23(69,441,862_76,269,868)×1, inv(18)(18q21.1)(48,739,965_49,016,148), fus(18)(18p11.32q21.32)(484,763_58,933,688), fus(18)(18p11.32q12.1)(603,123_31,941,501), fus(18)(18q11.2)(12,724,636_27,036,567), fus(18)(18p11.21q21.2)(12,797,857_49,016,148), fus(18)(18q11.2q12.2)(26,845,637_35,642,805), fus(18)(q22.2q23)(69,461,792_76,265,274) |
| **Long-read Sequencing** | seq[GRCh38] 18p11.32[479,581_609,746]×3,18p11.21 [12,720,506_12,799,427]×3, 18q11.2[26,838,028_27,038,666]×3, 18q11.2[27,038,666_27,038,727]×3, 18q11.2[27,039,152_27,039,386]×3, 18q12.1[31,933,516_32,042,120]×3, 18q12.1[31,933,498_31,933,517]×3, 18q12.1[32,042,094_32,042,114]×4, 18q12.1[32,042,118_32,042,145]×3, 18q12.2[35,577,193_35,646,246]×3, 18q21.1[48,730,407_49,020,934]×3, 18q21.2[50,831,095_50,842,336]×4, 18q21.1q21.32 [50,219,079_58,939,655]×3, 18q22.2q23[69,463,589_76,257,772]×1 |
| **Final Rearrangement** | 46,XY,der(18)(pter→18p11.32(chr18:609,746)::18q11.2(chr18:27,039,386)→18q11.2(chr18:27,039,152)::18q12.1(chr18:32,042,093)→18q12.1(chr18:32,042,115)::18q12.1(chr18:31,933,515)→18q12.1(chr18:32,042,120)::18q11.2(chr18:27,038,727)→18q11.2(chr18:27,038,665)::18q12.2(chr18:35,577,193)→18q12.2(chr18:35,646,246)::18q11.2(chr18:26,838,028)→18q11.2(chr18:27,038,666)::18p11.21(chr18:12,720,506)→18p11.21(chr18:12,799,427)::18q21.2(chr18:50,842,336)→18q21.2(chr18:50,831,095)::18q21.1(chr18:49,020,934)→18q21.1(chr18:48,730,407)::18q21.1(chr18:50,219,079)→18q21.32(chr18:58,939,655)::18q12.1(chr18:31,933,497)→18q12.1(chr18:31,933,517)::18q12.1(chr18:32,042,117)→18q12.1(chr18:32,042,145)::18p11.32(chr18:479,581)→18q22.2(chr18:69,463,589)::18q23(chr18:76,257,772)→qter) |
|  | 46,XY,der(18)(pter→18p11.32::18q11.2→18q11.2::18q12.1→18q12.1::18q12.1→18q12.1::18q11.2→18q11.2::18q12.2→18q12.2::18q11.2→18q11.2::18p11.21→18p11.21::18q21.2→18q21.2::18q21.1→18q21.1::18q21.1→18q21.32::18q12.1→18q12.1::18q12.1→18q12.1::18p11.32→18q22.2::18q23→qter) |

**Supplementary Table 3. Microduplications detected in Patient 6's father by CMA^a^.**

| **# Microduplication** | **Chromosome** | **Start** | **End** | **Size (bp)** |
| --- | --- | --- | --- | --- |
| 1 | chr18 | 485,968 | 603,871 | 117,903 |
| 2 | chr18 | 12,694,380 | 12,797,695 | 103,315 |
| 3 | chr18 | 26,844,671 | 27,035,736 | 191,065 |
| 4 | chr18 | 31,952,562 | 32,018,846 | 66,284 |
| 5 | chr18 | 46,497,916 | 46,535,244 | 37,328 |
| 6 | chr18 | 48,715,760 | 48,869,008 | 153,248 |
| 7 | chr18 | 48,956,537 | 49,002,757 | 46,220 |
| 8 | chr18 | 50,227,099 | 50,331,921 | 104,822 |
| 9 | chr18 | 58,865,588 | 58,931,521 | 65,933 |
| 10 | chr18 | 69,417,425 | 69,443,420 | 25,995 |
| 11 | chr18 | 76,251,700 | 76,321,765 | 70,065 |

^a^The platform used for CMA was CytoSNP-850K BeadChip *–* Illumina Technologies. All genomic coordinates are according to reference genome GRCh38/hg38.

**Supplementary Table 4. Breakpoint (BP) coordinates found through long-read sequencing and their number (#) for identification.**

| **#BP** | **BP coordinate** |
| --- | --- |
| 1 | chr18:479,581 |
| 2 | chr18:609,746 |
| 3 | chr18:12,720,506 |
| 4 | chr18:12,799,427 |
| 5 | chr18:26,838,028 |
| 6 | chr18:27,038,665 |
| 7 | chr18:27,038,666 |
| 8 | chr18:27,038,727 |
| 9 | chr18:27,039,152 |
| 10 | chr18:27,039,386 |
| 11 | chr18:31,933,497 |
| 12 | chr18:31,933,515 |
| 13 | chr18:31,933,517 |
| 14 | chr18:32,042,093 |
| 15 | chr18:32,042,114 |
| 16 | chr18:32,042,117 |
| 17 | chr18:32,042,120 |
| 18 | chr18:32,042,145 |
| 19 | chr18:35,577,193 |
| 20 | chr18:35,646,246 |
| 21 | chr18:48,730,407 |
| 22 | chr18:49,020,934 |
| 23 | chr18:50,219,079 |
| 24 | chr18:50,831,095 |
| 25 | chr18:50,842,336 |
| 26 | chr18:58,939,655 |
| 27 | chr18:69,463,589 |
| 28 | chr18:76,257,772 |

**Supplementary Table 5. Junction points and breakpoints details found on the long-read sequencing analysis and annotation of Repetitive Elements in regions surrounding the breakpoints.**

| **Junction Point** | | | | | | **Breakpoint** | | **Repetitive Elements** | | | **Obs.** |
| --- | --- | --- | --- | --- | --- | --- | --- | --- | --- | --- | --- |
| **#JP** | **Ins** | **MH** | **#SJP** | **Ins** | **MH** | **#BP** | **BP coord.** | **BP - 1 kb** | **BP** | **BP + 1 kb** |  |
| **1** | 269 | - | 1-1 | 7 | - | **2** | chr18:609,746 | DNA (TcMar-Tigger) – Tigger4  chr18:609364-609541 (-) | SINE (Alu) – AluSz  chr18:609542-609843 (+) | DNA (TcMar-Tigger) – Tigger4  chr18:609844-609899 (-) | 7 nt ins = AGTGGAG |
|  |  |  |  |  |  |  |  | DNA (TcMar-Tigger) – Tigger4  chr18:608978-609360 (-) |  |  |  |
|  |  |  |  |  |  | **10** | chr18:27,039,386 | - | LINE (L1) – L1MB3  chr18:27039134-27039690 (+) | - |  |
|  |  |  | 1-2 | 6 | - | **9** | chr18:27,039,152 |  |  |  | 6 nt ins =  AAAGTA |
|  |  |  |  |  |  | **14** | chr18:32,042,093 | SINE (Alu) – AluYa5  chr18:32041621-32041929 (-) | - | - |  |
|  |  |  | 1-3 | - | 2 (AT) | **15** | chr18:32,042,115 |  |  |  | - |
|  |  |  |  |  |  | **12** | chr18:31,933,515 | SINE (Alu) – AluY  chr18:31933021-31933320 (+) | SINE (MIR) – MIR  chr18:31933323-31933559 (+) | LINE (L1) – L1ME3Cz  chr18:31933862-31933905 (+) |  |
|  |  |  |  |  |  |  |  |  |  | SINE (Alu) – AluSz  chr18:31933906-31934195 (+) |  |
|  |  |  |  |  |  |  |  | SINE (Alu) – AluSx1  chr18:31932700-31933020 (+) |  | LINE (L1) – L1ME3Cz  chr18:31934196-31934418 (+) |  |
|  |  |  |  |  |  |  |  |  |  | Simple Repeat – (AAAC)n  chr18:31934498-31934521 (+) |  |
| **2** | 93 | - | 2-1 | 9 | - | **17** | chr18:32,042,120 | SINE (Alu) – AluYa5  chr18:32041621-32041929 (-) | - | SINE (Alu) – AluSz6  chr18:32042871-32043162 (+) | 9 nt ins =  TACTTTTAA |
|  |  |  |  |  |  | **8** | chr18:27,038,727 | SINE (Alu) – AluSz  chr18:27038266-27038550 (+) | DNA (hAT-Charlie) – Charlie5  chr18:27038572-27038831 (+) | DNA (hAT-Charlie) – MER33  chr18:27038830-27039005 (+) |  |
|  |  |  | 2-2 | 21 | - | **6** | chr18:27,038,665 |  |  |  | 21 nt ins =  TTAATTTAACTTTCAAGTTAA |
|  |  |  |  |  |  | **19** | chr18:35,577,193 | LINE (L1) – L1MD2  chr18:35576640-35576806 (+) | - | LINE (L1) – L1MD2  chr18:35577464-35577614 (+) |  |
|  |  |  |  |  |  |  |  | LTR (ERVL-MaLR) – THE1A  chr18:35576281-35576639 (-) |  |  |  |
|  |  |  |  |  |  |  |  | LINE (L1) – L1MD2  chr18:35576237-35576280 (+) |  |  |  |
| **3** | 2 (GA) | - | - | - | - | **20** | chr18:35,646,246 | LTR (ERVL-MaLR) – MLT1F1  chr18:35645879-35645978 (-) | LTR (ERVL-MaLR) – MSTA  chr18:35645979-35646361 (-) | LTR (ERVL-MaLR) – MLT1F1  chr18:35646362-35646711 (-) | - |
|  |  |  |  |  |  |  |  | LINE (L1) – L1M5  chr18:35645530-35645873 (-) |  | LINE (L2) – L2a  chr18:35646795-35647010 (+) |  |
|  |  |  |  |  |  |  |  | SINE (Alu) – AluY  chr18:35645249-35645529 (-) |  | LINE (L2) – L2a  chr18:35647074-35647432 (+) |  |
|  |  |  |  |  |  | **5** | chr18:26,838,028 | LINE (RTE-X) – L4_C_Mam  chr18:26837520-26837845 (+) | - | SINE (Alu) – AluY  chr18:26838221-26838515 (-) |  |
|  |  |  |  |  |  |  |  | SINE (MIR) – MIR  chr18:26837279-26837519 (+) |  | LINE (L2) – L2c  chr18:26838863-26839002 (-) |  |
|  |  |  |  |  |  |  |  | LINE (RTE-X) – L4_C_Mam  chr18:26837188-26837278 (+) |  |  |  |
| **Junction Point** | | | | | | **Breakpoint** | | **Repetitive Elements** | | | Obs. |
| **#JP** | **Ins** | **MH** | **#SJP** | **Ins** | **MH** | **#BP** | **BP coord.** | **BP - 1 kb** | **BP** | **BP + 1 kb** |  |
| **4** | - | - | - | - | - | **7** | chr18:27,038,666 | SINE (Alu) – AluSz  chr18:27038266-27038550 (+) | DNA (hAT-Charlie) – Charlie5  chr18:27038572-27038831 (+) | DNA (hAT-Charlie) – MER33  chr18:27038830-27039005 (+) | - |
|  |  |  |  |  |  |  |  |  |  | Simple Repeat – (AAC)n  chr18:27039029-27039063 (+) |  |
|  |  |  |  |  |  |  |  |  |  | LINE (L1) – L1MB3  chr18:27039134-27039690 (+) |  |
|  |  |  |  |  |  | **3** | chr18:12,720,506 | SINE (Alu) – AluYj4  chr18:12720039-12720127 (-) | - | SINE (Alu) – AluYj4  chr18:12720738-12720829 (+) |  |
|  |  |  |  |  |  |  |  | SINE (Alu) – AluSx3  chr18:12719699-12720016 (-) |  | SINE (Alu) – AluY  chr18:12720884-12721195 (+) |  |
|  |  |  |  |  |  |  |  | SINE (Alu) – AluSx1  chr18:12719388-12719686 (-) |  | LINE (L1) – L1MB4  chr18:12721363-12721970 (-) |  |
| **5** | - | 282 | - | - | - | **4** | chr18:12,799,427 | DNA (hAT-Charlie) – Charlie9  chr18:12798931-12799047 | SINE (Alu) – AluSg  chr18:12799137-12799443 (+) | SINE (Alu) – AluSz  chr18:12799579-12799930 (-) | 83% homeology |
|  |  |  |  |  |  |  |  | LINE (L1) – L1PA13  chr18:12798248-12798922 |  |  |  |
|  |  |  |  |  |  | **25** | chr18:50,842,336 | SINE (MIR) – MIRb  chr18:50841759-50841929 (+) | SINE (Alu) – AluSz  chr18:50842065-50842346 (-) | LINE (L2) – L2b  chr18:50842364-50842436 (+) |  |
|  |  |  |  |  |  |  |  |  |  | Simple Repeat – (T)n  chr18:50842569-50842594 (+) |  |
|  |  |  |  |  |  |  |  | SINE (Alu) – AluY  chr18:50841269-50841570 (-) |  | LINE (L1) – L1ME4a  chr18:50842729-50842876 (-) |  |
|  |  |  |  |  |  |  |  |  |  | LTR (ERVL-MaLR) – MLT1B  chr18:50842905-50842925 (-) |  |
|  |  |  |  |  |  |  |  |  |  | LTR (ERVL) – MLT2D  chr18:50842926-50843332 (-) |  |
| **6** | 8 | - | - | - | - | **24** | chr18:50,831,095 | DNA (hAT-Tip100) – MER96  chr18:50830617-50830696 (-) | SINE (Alu) – AluSg4  chr18:50830849-50831151 (-) | LINE (L2) – L2c  chr18:50831321-50831415 (+) | 8nt ins =  TGTACTGT |
|  |  |  |  |  |  |  |  | SINE (Alu) – AluY  chr18:50830235-50830536 (+) |  |  |  |
|  |  |  |  |  |  |  |  | SINE (Alu) – AluSx  chr18:50829934-50830233 (+) |  |  |  |
|  |  |  |  |  |  | **22** | chr18:49,020,934 | - | SINE (MIR) – MIRb  chr18:49020860-49020993 (+) | DNA (hAT-Tip100) – MER91C  chr18:49021956-49022022 (-) |  |
| **Junction Point** | | | | | | **Breakpoint** | | **Repetitive Elements** | | | **Obs.** |
| **#JP** | **Ins** | **MH** | **#SJP** | **Ins** | **MH** | **#BP** | **BP coord.** | **BP - 1 kb** | **BP** | **BP + 1 kb** |  |
| **7** | 25 | - | - | - | - | **21** | chr18:48,730,407 | DNA (hAT-Charlie) – MER3  chr18:48730039-48730178 (-) | - | - | 25nt ins =  CAGGCAGGTGGTACGGGCAGGGAGT |
|  |  |  |  |  |  |  |  | SINE (MIR) – MIR3  chr18:48729553-48729644 (-) |  |  |  |
|  |  |  |  |  |  | **23** | chr18:50,219,079 | LTR (ERVL?) – LTR55  chr18:50218921-50219038 (-) | LTR (ERVL) – LTR16E1  chr18:50219060-50219364 (+) | SINE (MIR) – MIR3  chr18:50219369-50219434 (+) |  |
|  |  |  |  |  |  |  |  | LTR (ERVL?) – LTR55  chr18:50218436-50218814 (+) |  |  |  |
|  |  |  |  |  |  |  |  | LINE (L2) – L2b  chr18:50218125-50218163 (-) |  |  |  |
|  |  |  |  |  |  |  |  | LTR (ERVL-MaLR) – MLT1B  chr18:50217809-50218124 (-) |  |  |  |
| **8** | 51 | - | 8-1 | - | - | **26** | chr18:58,939,655 | SINE (Alu) – AluSz6  chr18:58938965-58939275 (+) | - | SINE (Alu) – AluSc8  chr18:58939896-58940192 (-) |  |
|  |  |  |  |  |  |  |  |  |  | SINE (MIR) – MIRb  chr18:58940488-58940675 (+) |  |
|  |  |  |  |  |  | **11** | chr18:31,933,497 | SINE (Alu) – AluY  chr18:31933021-31933320 (+) | SINE (MIR) – MIR  chr18:31933323-31933559 (+) | LINE (L1) – L1ME3Cz  chr18:31933862-31933905 (+) |  |
|  |  |  | 8-2 | - | - | **13** | chr18:31,933,517 |  |  |  |  |
|  |  |  |  |  |  | **16** | chr18:32,042,117 | SINE (Alu) – AluYa5  chr18:32041621-32041929 (-) | - | - |  |
|  |  |  | 8-3 | 1 (G) | - | **18** | chr18:32,042,145 |  |  |  |  |
|  |  |  |  |  |  | **1** | chr18:479,581 | Simple Repeat – (CT)n  chr18:479527-479551 (+) | Simple Repeat – (AC)n  chr18:479552-479593 (+) | Low_complexity – A-rich  chr18:479715-479747 (+) |  |
|  |  |  |  |  |  |  |  |  |  | LTR (ERVL-MaLR) – MLT1D  chr18:479779-480260 (-) |  |
| **9** | 9 | - | - | - | - | **27** | chr18:69,463,589 | LTR (ERVL-MaLR) – MLT1A0  chr18:69463006-69463352 (-) | - | Simple Repeat – (TGTGTG)n  chr18:69463639-69463824 (+) | 9nt ins =  TCCAGTATC |
|  |  |  |  |  |  | **28** | chr18:76,257,772 | - | - | Simple Repeat – (A)n  chr18:76258004-76258024 (+) |  |
| **#JP** | **Ins** | **MH** | **#SJP** | **Ins SJP** | **MH** | **#BP** | **BP coord.** | **BP - 1 kb** | **BP** | **BP + 1 kb** | **Obs.** |
| **Junction Point** | | | | | | **Breakpoint** | | **Repetitive Elements** | | |  |

JP: junction point; JP ID: junction point identification; Ins: insertion; SJP: sub-junction point; MH: microhomology; Del: deletion; BP: breakpoint; coord.: coordinate; SNVs: single nucleotide variants; Obs.: observation. All genomic coordinates are according to reference genome GRCh38/hg38.

**Supplementary Table 6. Copy number variants found on Patient 6's lrGS and comparison with the microduplications found on his father's CMA.**

| **Alteration** | **Chromosome** | **Start** | **End** | **Size (bp)** | **Relation with the father’s microduplications** |
| --- | --- | --- | --- | --- | --- |
| Microduplication | chr18 | 479,581 | 609,746 | 130,165 | Overlaps microduplication 1 |
| Microduplication | chr18 | 12,720,506 | 12,799,427 | 78,921 | Overlaps microduplication 2 |
| Microduplication | chr18 | 26,838,028 | 27,038,666 | 200,638 | Overlaps microduplication 3 |
| Microduplication | chr18 | 27,038,666 | 27,038,727 | 62 | None |
| Microduplication | chr18 | 27,039,152 | 27,039,386 | 234 | None |
| Microduplication | chr18 | 31,933,516 | 32,042,120 | 108,604 | Overlaps microduplication 4 |
| Microduplication | chr18 | 31,933,498 | 31,933,517 | 20 | None |
| Microduplication | chr18 | 32,042,094 | 32,042,114 | 21 | None |
| Microduplication | chr18 | 32,042,118 | 32,042,145 | 28 | None |
| Microduplication | chr18 | 35,577,193 | 35,646,246 | 69,053 | None |
| Microduplication | chr18 | 48,730,407 | 49,020,934 | 290,527 | Overlaps microduplications 6 and 7 |
| Microduplication | chr18 | 50,831,095 | 50,842,336 | 11,241 | None |
| Duplication | chr18 | 50,219,079 | 58,939,655 | 8,720,576 | Start ~8 kb upstream of microdup 8 then overlaps;  End ~8kb downstream of microdup 9 after overlap |
| Deletion | chr18 | 69,463,589 | 76,257,772 | 6,794,183 | Start ~20 kb downstream of microdup 10 (no overlap);  End overlaps ~6 kb of microdup 11 |

All genomic coordinates are according to reference genome GRCh38/hg38.

**Supplementary Table 7. Difference between breakpoints found by Optical Genome Mapping (OGM) and Long-read Sequencing (lrGS).**

| **#JP** | **BP** | **BP OGM** | | **BP lrGS** | **BPs difference (bp)** | **Sum of BPs difference (bp)** | **Size of Uncertain OGM region (bp)** |
| --- | --- | --- | --- | --- | --- | --- | --- |
| 1 | 1 | chr18:603,123 | | chr18:609,746 | 6,623 | 14,609 | 14,930 |
|  | 2 | NF | | chr18:27,039,386 | - |  |  |
|  | 3 | NF | | chr18:27,039,152 | - |  |  |
|  | 4 | NF | | chr18:32,042,093 | - |  |  |
|  | 5 | NF | | chr18:32,042,115 | - |  |  |
|  | 6 | chr18:31,941,501 | | chr18:31,933,515 | 7,986 |  |  |
| 2 | 7 | chr18:32,037,307 | | chr18:32,042,120 | 4,813 | 11,125 | 11,125 |
|  | 8 | NF | | chr18:27,038,727 | - |  |  |
|  | 9 | NF | | chr18:27,038,665 | - |  |  |
|  | 10 | chr18:35,583,505 | | chr18:35,577,193 | 6,312 |  |  |
| 3 | 11 | chr18:35,642,805 | | chr18:35,646,246 | 3,441 | 11,043 | 11,260 |
|  | 12 | chr18:26,845,637 | | chr18:26,838,028 | 7,609 |  |  |
| 4 | 13 | chr18:27,036,567 | | chr18:27,038,666 | 2,099 | 2,031 | 6,260 |
|  | 14 | chr18:12,724,636 | | chr18:12,720,506 | 4,130 |  |  |
| 5 | 15 | chr18:12,797,857 | | chr18:12,799,427 | 1,570 | 6,356 | 18,143 |
|  | 16 | NF | | chr18:50,842,336 | - |  |  |
| 6 | 17 | NF | | chr18:50,831,095 | - |  |  |
|  | 18 | chr18:49,016,148 | | chr18:49,020,934 | 4,786 |  |  |
| 7 | 19 | chr18:48,739,965 | | chr18:48,730,407 | 9,558 | 14,826 | 14,916 |
|  | 20 | chr18:50,224,347 | | chr18:50,219,079 | 5,268 |  |  |
| 8 | 21 | chr18:58,933,688 | | chr18:58,939,655 | 5,967 | 11,149 | 11,652 |
|  | 22 | NF | | chr18:31,933,497 | - |  |  |
|  | 23 | NF | | chr18:31,933,517 | - |  |  |
|  | 24 | NF | | chr18:32,042,117 | - |  |  |
|  | 25 | NF | | chr18:32,042,145 | - |  |  |
|  | 26 | chr18:484,763 | | chr18:479,581 | 5,182 |  |  |
| 9 | 27 | chr18:69,461,792 | | chr18:69,463,589 | 1,797 | 9,299 | 9,310 |
|  | 28 | chr18:76,265,274 | | chr18:76,257,772 | 7,502 |  |  |
|  | | | **Average** | | 5,290 |  |  |
|  | | | **Standard Deviation** | | 2,317 |  |  |

JP: junction point; BP(s): breakpoint(s); bp: base pair; NF: not found. All genomic coordinates are according to reference genome GRCh38/hg38.

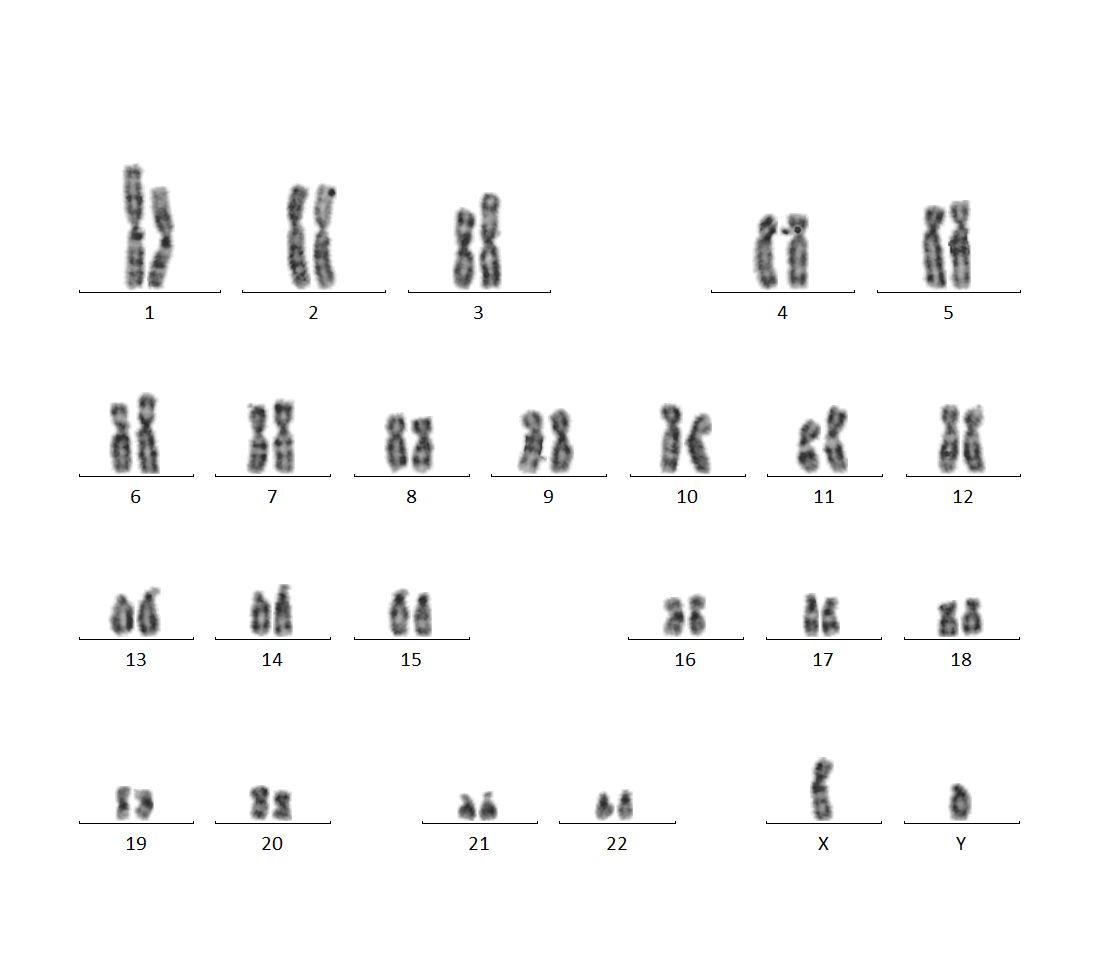

**Supplementary Figure 1. Karyotype of Patient 6.**

The red arrow highlights the chromosome involved in the patient’s rearrangement.

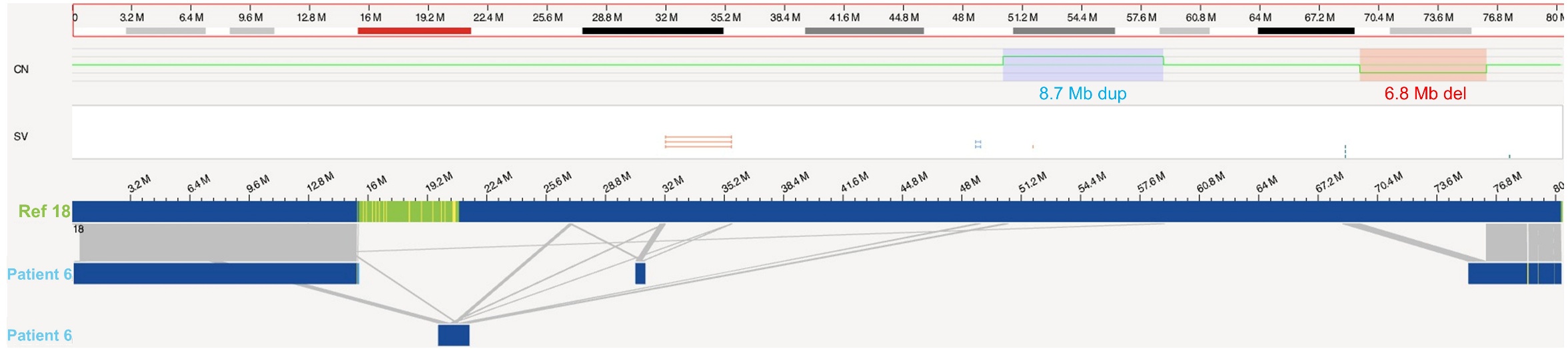

**Supplementary Figure 2. Optical Genome Mapping of Patient 6.**

The image shows the genome browser view of the patient’s chromosome 18. Both the 8.7 Mb duplication and the 6.8 Mb deletion in 18q were called by the CNV pipeline. Patient maps show alignments to multiple regions across the chromosome.

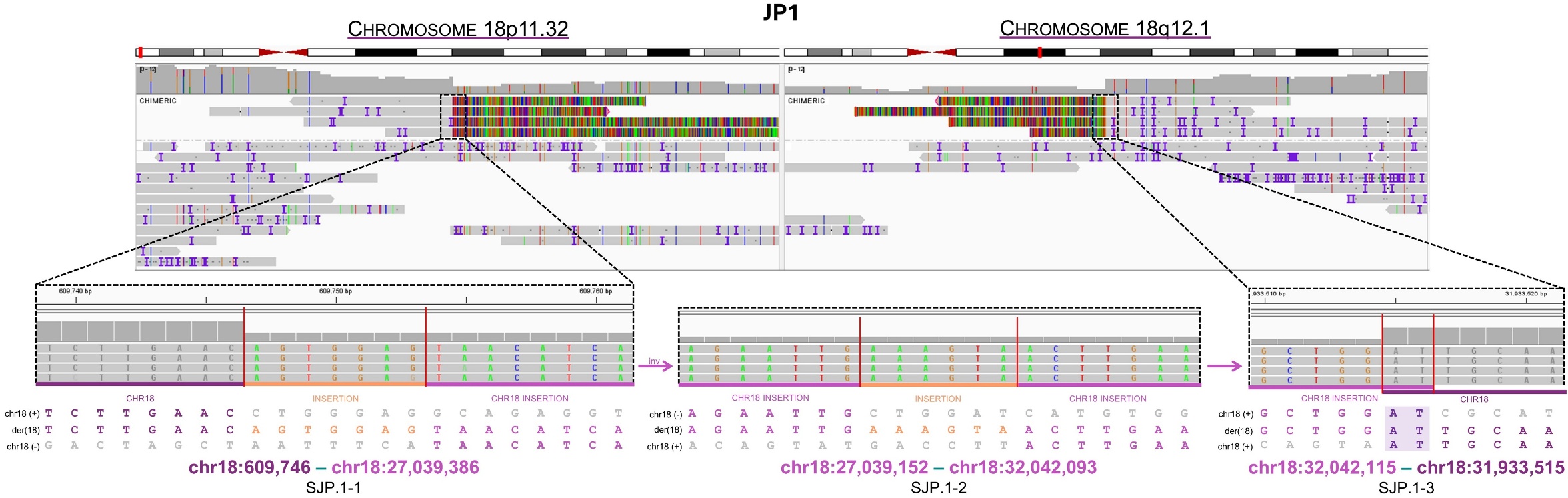

**Supplementary Figure 3. Patient 6’s Junction Point 1 seen through long-read sequencing chimeric reads on IGV.**

At the top, IGV view of both involved regions showing the chimeric reads. At the bottom, zoom in on the junction point at the nucleotide level with the sequence alignment and breakpoints below.

The patient’s JP1 presents two other regions of chromosome 18 between the breakpoints, thus forming three subjunctions. SJP-1 and SJP.1-2 show insertions of seven and six nucleotides, respectively, while SJP.1-3 presents two nucleotides of microhomology.

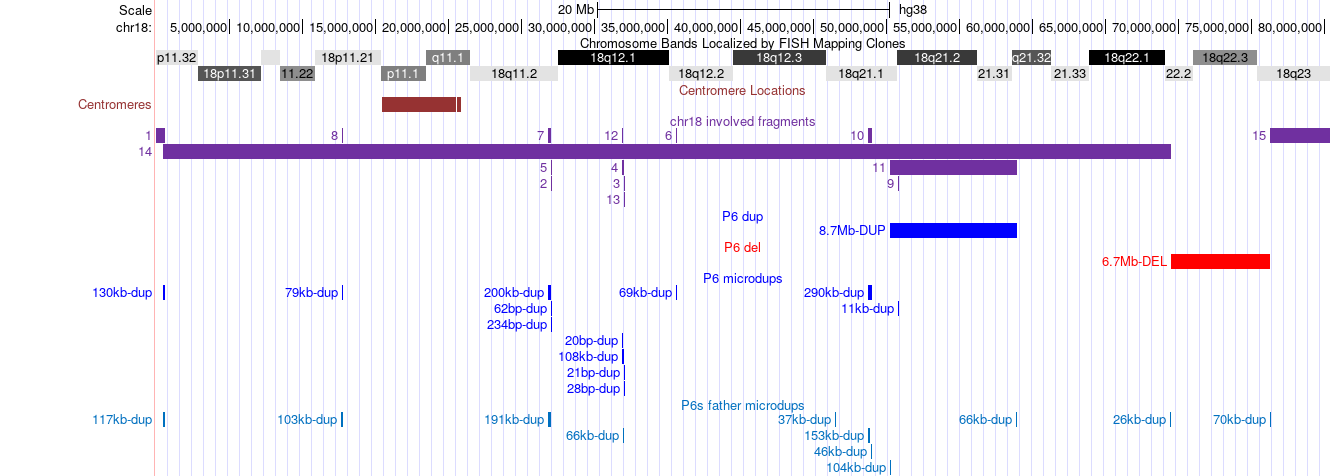

**Supplementary Figure 4. UCSC Genome Browser view of chromosome 18 showing Patient 6’s and his father’s CNVs.**

At the top, coordinates, chromosomal bands, and centromere location of chromosome 18. Below, we see five different tracks with their respective annotations. The first one, “chr18 involved fragments”, show which regions of the chromosome are present in the der(18) and how many copies they have. The second and the third ones, “P6 dup” and “P6 del”, show the location of the 8.7 Mb duplication and the 6.7 Mb deletion, respectively. The fourth, “P6 microdups”, highlights the location of Patient 6’s microduplications with their respective sizes. The fifth, “P6s father microdups”, exhibit the location and size of his father’s microduplications. The image shows a general view of chromosome 18 to show the patient’s CNVs and to compare his microduplications with his father’s.

Patient results are according to lrGS. Father results are according to CMA.

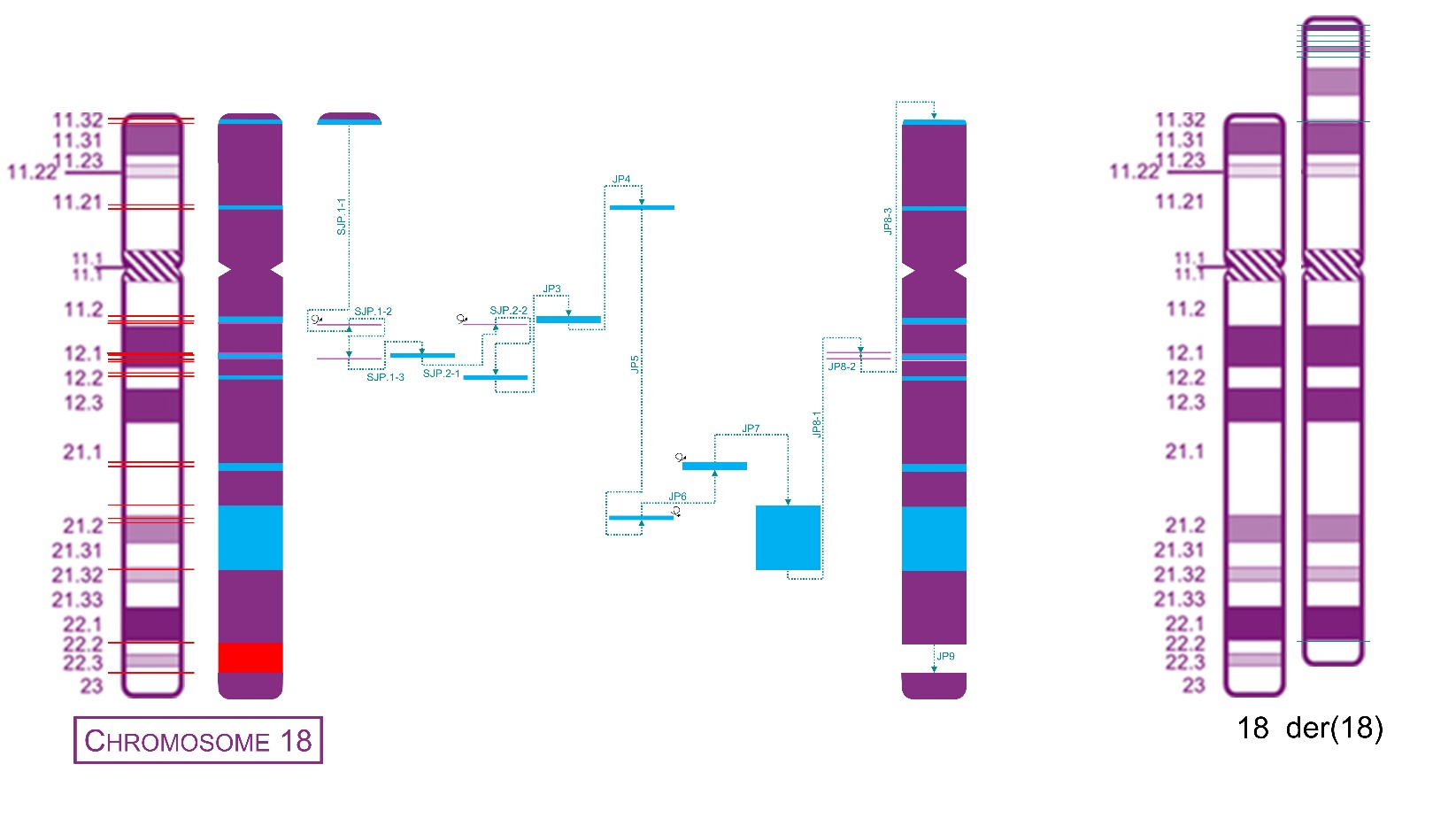

**Supplementary Figure 5. Final rearrangement of Patient 6’s rearranged chromosome 18.**

To the left, a schematic representation of the junctions between all chromosome 18 regions involved in the CGR. To the right, idiogram of the chromosome involved in the complex genomic rearrangement. Red lines show the breakpoints in the normal chromosomes and cyan lines show the junction points in the derivative chromosomes.
